# A unified benchmark of synthetic data generation for clinical transcriptomic cancer cohorts

**DOI:** 10.64898/2026.05.13.724858

**Authors:** The-Chuong Trinh, Jean-Baptiste Woillard, Guido Uguzzoni, Christophe Battail

## Abstract

Achieving a trade-off between biological utility and patient privacy remains a key challenge for secure data sharing when applying transcriptomic clinical datasets to artificial intelligence in precision oncology. Here, we introduce the first benchmarking study tailored to high-dimensional clinical transcriptomic cancer data, comparing synthetic data generation methods across three clinical cancer trials. Our framework, SynOmicsBench, combines standardized preprocessing with multidimensional evaluation, prioritizing downstream biological validation alongside statistical fidelity and attack-based privacy assessment. Results indicate that no single method dominated all dimensions, with Gaussian Copula achieving the most balanced performance, followed by Avatar, demonstrating that metric-based similarity alone is insufficient to ensure preservation of higher-order molecular dependencies. Synthetic data consistently reproduced biomedical signal directionality but with attenuated effect sizes and inter-replicate variability, supporting hypothesis generation when multi-seed synthesis is adopted. Collectively, this framework provides a reproducible decision-support tool for method selection and promotes biologically informed, privacy-aware adoption of synthetic data in precision oncology.

## 1 Introduction

High-throughput transcriptomic profiling generated within drug clinical trials has become a cornerstone of contemporary cancer research, enabling genome-wide quantification of gene expression across well-characterized patient cohorts.^1^ These datasets support a broad spectrum of translational applications, including biomarker discovery, patient stratification, and the prediction of clinical outcomes.^1-4^ Because they are collected in controlled clinical settings, these datasets constitute high-quality resources that underpin artificial intelligence (AI) approaches in precision oncology that increasingly inform therapeutic decision-making.^5^

Progress in this area critically depends on the ability to share and reuse data across institutions.^6^ However, clinical transcriptomic datasets are inherently sensitive, as they capture both high-dimensional molecular phenotypes and potentially identifiable patient information.^6^ Their access and use are therefore governed by stringent regulatory frameworks, including the General Data Protection Regulation (GDPR)^7^ in the European Union and the Health Insurance Portability and Accountability Act (HIPAA)^8^ in the United States. In practice, these regulations impose substantial constraints on data access, requiring complex governance procedures such as data transfer agreements and data protection impact assessments. As a result, cross-institutional data sharing remains limited, hindering collaborative research, restricting external validation of artificial intelligence (AI) models, and slowing the assembly of sufficiently large and diverse cohorts needed for robust AI-based developments. Conventional privacy-preserving strategies, such as pseudonymization and anonymization, provide only partial mitigation.^9^ While pseudonymization reduces direct identifiability, it does not eliminate re-identification risks under regulatory definitions. Conversely, full anonymization often requires substantial aggregation at the cohort level, leading to the loss of individual-level signal and rendering the data largely unusable for precision oncology applications.^10^

Synthetic data generation (SDG) has emerged as a promising strategy to alleviate constraints on data access and sharing in biomedical research.^10^ In this context, synthetic data refers to artificially generated datasets designed to replicate the statistical properties of real patient-level data while avoiding direct exposure to personal information. In the healthcare domain, SDG approaches have been primarily investigated in medical imaging and tabular clinical data.^11,12^ Their potential applications are diverse beyond data sharing, ranging from enabling controlled data access without exposing raw patient records to augmenting limited cohorts of rare clinical or molecular traits that are under-represented in real-world datasets.^10,13^ Advances in deep generative modeling, including Generative Adversarial Networks (GANs),^14^ Variational Autoencoders (VAEs),^15^ and copula-based statistical frameworks,^16^ have substantially improved the quality and applicability of synthetic biomedical data in recent years. Despite these advances, the generation and benchmarking of synthetic high-dimensional omics datasets, such as transcriptomics, remain complex and unaddressed challenges.

Indeed, transcriptomic data presents a unique combination of statistical and biological challenges that are not adequately captured by existing SDG evaluation frameworks.^10,17^ These datasets are inherently high-dimensional, typically comprising tens of thousands of genes. At the same time, clinical cancer cohorts are often limited to a few hundred patients, resulting in an extreme p ≫ n setting. In addition, gene expression data exhibit strong correlation structures driven by co-expression programs, pathway activity, and underlying gene regulatory networks. These features coexist with heterogeneous clinical variables, including continuous measurements, categorical covariates, and time-to-event outcomes, within complex, multimodal datasets. Furthermore, these datasets are often affected by noise, sparsity, and missingness arising from both technical and clinical sources. In this context, synthetic datasets may reproduce marginal statistical properties while failing to preserve high-order biological organization, thereby limiting their downstream utility. Collectively, these considerations underscore the need for rigorous, context-aware benchmarking strategies specifically tailored to the complexity of transcriptomic data in clinical trial settings.

To address these challenges, we present SynOmicsBench, a systematic and application-driven benchmarking framework for SDG methods applied to transcriptomic cancer datasets from clinical trials. We evaluate a representative set of approaches spanning deep generative models and statistical methods across multiple complementary dimensions, including statistical fidelity, the preservation of biological and clinical utility, and privacy risk. ***Statistical fidelity*** quantifies how closely synthetic data recapitulate the distributional and correlational properties of the original dataset. ***Biological utility*** assesses whether synthetic data preserve scientifically relevant signals present in the real data, including differential gene expression (DGE), pathway enrichment, immune cell composition, and survival-associated transcriptomic patterns. The distinction between these two dimensions is methodologically critical: while statistical fidelity can be evaluated using established, domain-agnostic metrics, biological utility is inherently context-dependent and cannot be reduced to a single generalizable measure. For privacy evaluation, rather than relying on distance-based metrics commonly used in prior benchmarks, we implement regulatory-grade metrics aligned with practical risk assessment frameworks.^7,18^ In addition, computational cost and scalability, which are important considerations for real-world deployment, are systematically reported. Our approach is therefore grounded in a multi-metric evaluation strategy aiming to capture the complexity of transcriptomic data and the trade-offs between fidelity, utility, and privacy. To achieve this, we introduce a rank-based scoring system to aggregate multiple metrics into an interpretable comparative framework, thereby facilitating method selection in practice.

We apply this framework to benchmark five representative SDG methods spanning generative deep learning and statistical modelling paradigms, using three independent transcriptomic datasets collected from clinical immunotherapy trials in kidney cancer, melanoma, and lung cancer.^3,19,20^ Beyond comparative evaluation, this framework provides structured guidance on the current reliability of synthetic transcriptomic data, identifying the biological and clinical questions for which existing methods are adequate, the contexts where caution is warranted, and the key directions for future methodological development. Validated synthetic transcriptomic data have the potential to transform translational cancer research by enabling broader access to high-value clinical datasets, and accelerate the development of digital twins and in silico trials.

## 2 Results

### 2.1 Overview of benchmarking protocol

We established a standardized benchmarking protocol evaluating five synthetic data generation (SDG) methods: CTGAN,^21^ TVAE,^21^ Gaussian Copula,^16^ Synthpop,^22^ and Avatar (K5/K10).^12^ These methods were evaluated across three cancer cohorts: CheckMate (CM) 009, 010, and 025 for clear cell renal cell carcinoma (ccRCC) from Braun et al.,^3^ melanoma from Liu et al.,^19^ and Stand Up To Cancer-Mark Foundation (SU2C-MARK) for non-small cell lung cancer (NSCLC) from Ravi et al.^20^ All clinical trial cohorts included patients treated with immune checkpoint blockade and comprised sensitive clinical outcomes alongside high-dimensional bulk transcriptomic profiles (Table 1). The protocol applied a standardized preprocessing pipeline including quality control and multivariate imputation to ensure comparable model inputs and minimize confounding from data quality discrepancies. Models were then trained using five independent random seeds, generating 90 synthetic datasets to assess the robustness to stochastic training variability. Quality was evaluated across three pillars: statistical fidelity (univariate and bivariate fidelity), biological and clinical utility (downstream analyses including differential gene expression (DGE),^23^ gene set enrichment analysis (GSEA),^24^ single-sample GSEA (ssGSEA),^25^ cell type deconvolution,^26^ and survival analysis), and privacy risk (singling-out, linkability, and inference attacks) aligned with the European Data Protection Board (EDPB).^27^ Computational cost was reported alongside quality metrics to assess practical feasibility and scalability. Finally, a rank-derived meta-score was calculated to identify methods providing the best trade-off between utility and privacy preservation (Figure 1).

**Table 1.** Clinical and transcriptomic characteristics of three cancer datasets.

| Dataset characteristic |  | ccRCC | Melanoma | NSCLC |
| --- | --- | --- | --- | --- |
| Number of patients |  | 311 | 121 | 152 |
| Number of clinical features |  | 52 | 47 | 14 |
| Number of transcriptomics features |  | 40934 | 18760 | 21969 |
| Gene expression level |  | TPM | TPM | TPM |
| Age (year), median (range) |  | 63 (30-88) |  | 64 (40-89) |
| Sex, no. | Male | 229 | 71 | 87 |
|  | Female | 82 | 50 | 65 |
| Clinical outcomes | PR/CR | 44 | 47 | 60 |
|  | PD | 106 | 56 | 50 |
|  | SD | 131 | 16 | 42 |
|  | Others | 30 | 2 | 0 |

TPM: Transcripts Per Kilobase Million; PR: Partial Response; CR: Complete Response; SD: Stable Disease; PD: Progressive Disease

**Figure 1.**
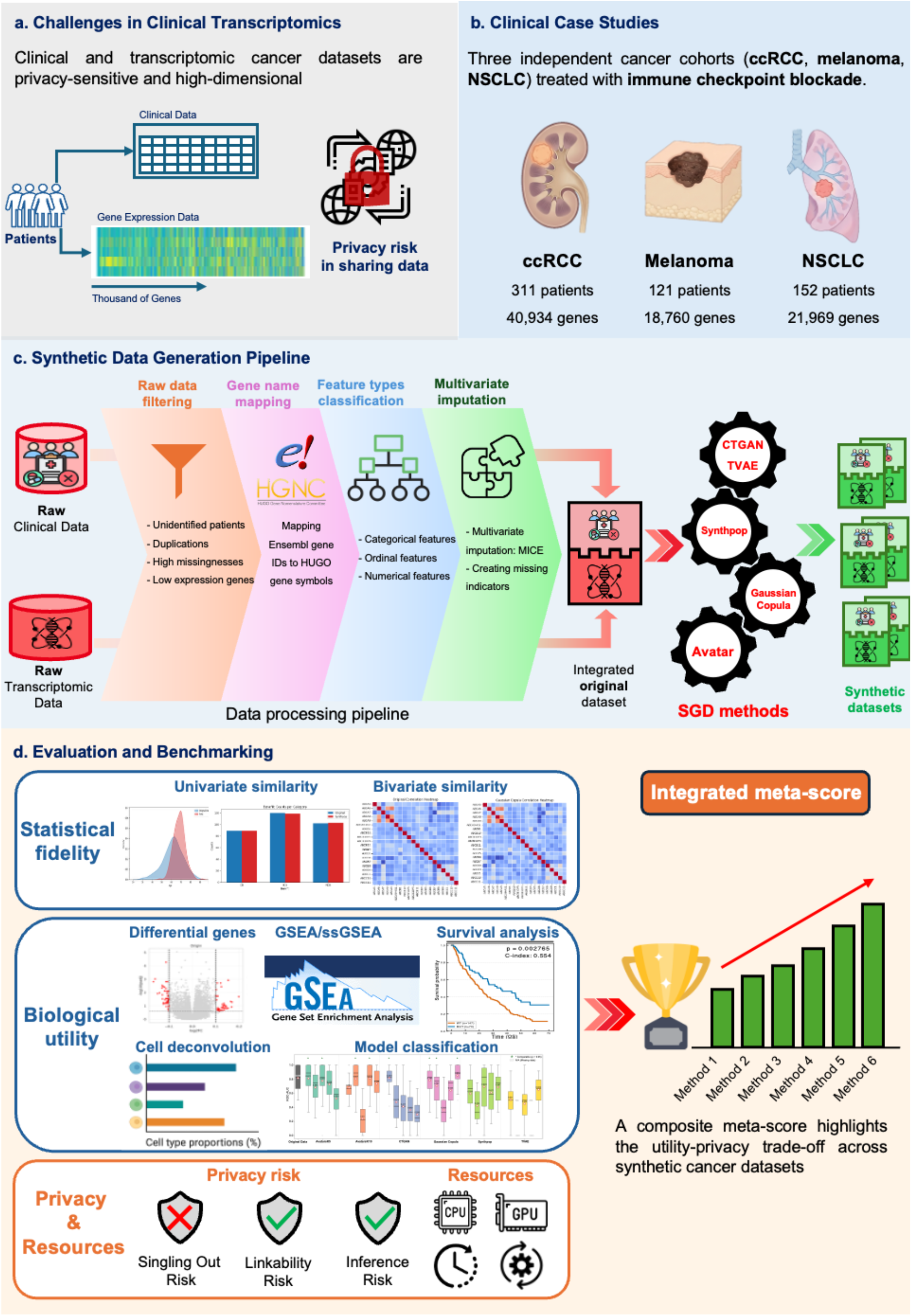
Overview of the benchmarking protocol for synthetic clinical transcriptomic data. **a) Data sensitivity and dimensionality**. Clinical transcriptomic datasets comprise sensitive patient-level information alongside with high-dimensional gene expression profiles (thousands of genes per sample). While cross-institutional data sharing is essential for collaborative oncology research, these modalities pose significant privacy risks. **b) Oncology case studies**. Three cancer types were selected as benchmarking case studies: ccRCC, Melanoma, NSCLC. The included clinical cohorts differ in sample size and transcriptomic dimensionality, reflecting realistic heterogeneity across oncology datasets. **c) Synthetic data generation pipeline**. Clinical and transcriptomic data were first harmonized and integrated through a standardized data processing pipeline. The processed dataset was subsequently used to train SDG models, which generated patient-level synthetic multimodal datasets. **d) Evaluation framework**. The benchmarking framework comprises three evaluation pillars. Statistical fidelity assesses global statistical fidelity through univariate and bivariate similarity metrics. Biological utility evaluates task-specific performance in clinically and biologically relevant downstream analyses, including DGE, GSEA, ssGSEA, cell deconvolution, survival analysis, and predictive model classification. Privacy risk quantifies disclosure vulnerability via singling-out, linkability, and inference attacks. Computational resource consumption is reported alongside synthetic data quality metrics. Finally, a rank-derived meta-score is calculated for each SDG model to identify the optimal trade-off between utility and privacy preservation.

### 2.2 Statistical fidelity

Statistical fidelity assessments quantified how well synthetic data mirrored the marginal distributions and inter-variable relationships of the original cohorts. Table 2 summarizes the performance of the six SDG approaches; mathematical and computational details are provided in the Methods section.

**Table 2.** Statistical fidelity assessment of synthetic datasets across three cancer cohorts based on univariate and bivariate similarity scores. Higher scores (approaching 1.0) indicate a stronger ability of the synthetic data to mirror the statistical distributions and pairwise relationships of the original data.

| Synthetic dataset | ccRCC | Melanoma | NSCLC |
| --- | --- | --- | --- |
| **Univariate similarity score** |  |  |  |
| Avatar K5 | 0.890 ± 0.001 | 0.857 ± 0.002 | 0.867 ± 0.003 |
| Avatar K10 | 0.869 ± 0.002 | 0.843 ± 0.002 | 0.853 ± 0.002 |
| CTGAN | 0.818 ± 0.010 | 0.767 ± 0.016 | 0.762 ± 0.021 |
| Gaussian copula | 0.889 ± 0.003 | 0.893 ± 0.001 | 0.901 ± 0.005 |
| Synthpop | 0.952 ± 0.001 | 0.924 ± 0.005 | 0.925 ± 0.004 |
| TVAE | 0.627 ± 0.027 | 0.802 ± 0.017 | 0.747 ± 0.022 |
| **Bivariate similarity score\*** |  |  |  |
| Avatar K5 | 0.954 ± 0.001 | 0.934 ± 0.001 | 0.940 ± 0.002 |
| Avatar K10 | 0.952 ± 0.001 | 0.932 ± 0.000 | 0.918 ± 0.015 |
| CTGAN | 0.849 ± 0.000 | 0.916 ± 0.000 | 0.875 ± 0.001 |
| Gaussian copula | 0.935 ± 0.003 | 0.938 ± 0.001 | 0.938 ± 0.002 |
| Synthpop | 0.905 ± 0.001 | 0.925 ± 0.001 | 0.900 ± 0.003 |
| TVAE | 0.805 ± 0.011 | 0.908 ± 0.004 | 0.863 ± 0.019 |

Each cell presents the mean scores and their standard deviations from five independent replicates for each method.
\* The bivariate scores are derived from all feature pairs $\frac{N(N-1)}{2}$ , where N is the total number of features (clinical, transcriptomics)

Most methods achieved high univariate similarity scores across cohorts, with the exception of TVAE and CTGAN, which showed the lowest values and highest variability (Table 2). Clinical attributes were generally slightly better reproduced than transcriptomic features (Figure 2a and Figure S2), reflecting the challenges posed by the high dimensionality of gene expression data. Bayesian pairwise comparisons between SDG models across three cohorts revealed that Gaussian Copula and Synthpop achieved the highest statistical fidelity, followed by the Avatar method (Figure 2b and Tables S1, S2, and S3).

**Figure 2.**
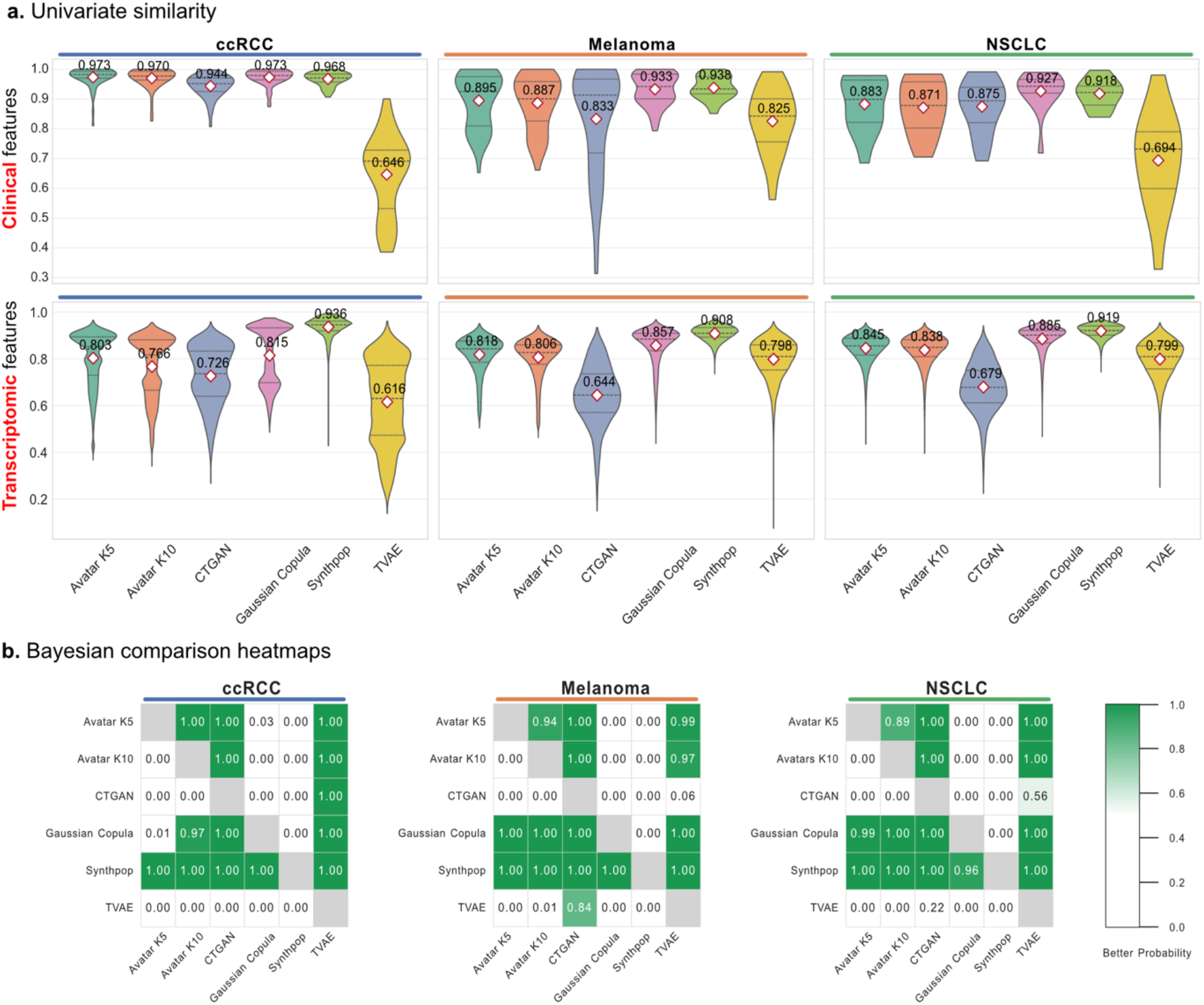
Comparative analysis of univariate statistical similarity across cancer cohorts. **a) Violin plot distribution of univariate similarity scores**. Violin plots display the distribution of univariate similarity scores for clinical variables and transcriptomic features across the ccRCC, melanoma, and NSCLC cohorts. Numerical attributes were evaluated using the KS statistic, whereas categorical attributes were assessed using TVD. Each violin represents the feature-level distribution of similarity scores within a cohort and data modality. The broader distributions and lower mean scores of gene expression features compared to clinical features highlight the increased complexity of preserving high-dimensional transcriptomic data. Shown panels represent one representative replicate; results from additional replicates are provided in Figure S2 in the Supplementary Data. **b) Bayesian pairwise comparative probability heatmap**. Heatmaps illustrate pairwise “better probabilities” estimated using a Bayesian comparison analysis. Each cell represents *P*(*row* > *column*), defined as the posterior probability that the SDG method in the row achieves superior univariate similarity compared with the method in the column for the corresponding cohort. Values range from 0 to 1, with darker green indicating stronger evidence of superiority.

Bivariate statistics were highly informative for discriminating between methods: Gaussian Copula and Avatar (K5/K10) consistently outperformed the other approaches in preserving overall pairwise relationships between synthetic and original datasets (Table 2, Figure 3a, and Tables S4, S5, and S6). However, the high mean bivariate score reported in Table 2 was largely driven by the large number of weakly correlated feature pairs, which inflate overall similarity and can obscure the meaningful performance differences. Restricting the analysis to strongly correlated pairs of features (|*ρ*| ≥ 0.5) revealed clearer disparities between methods (Figure 3b and Figure S3): Avatar (K5/K10) and Gaussian Copula most consistently preserved clinical-clinical and transciptomic-transcriptomic relationships across cohorts, while Synthpop performed only on transcriptomic ones. All methods showed reduced capacity to reproduce clinical-transcriptomic associations compared to within-type associations (clinical-clinical and transcriptomic-transcriptomic).

**Figure 3.**
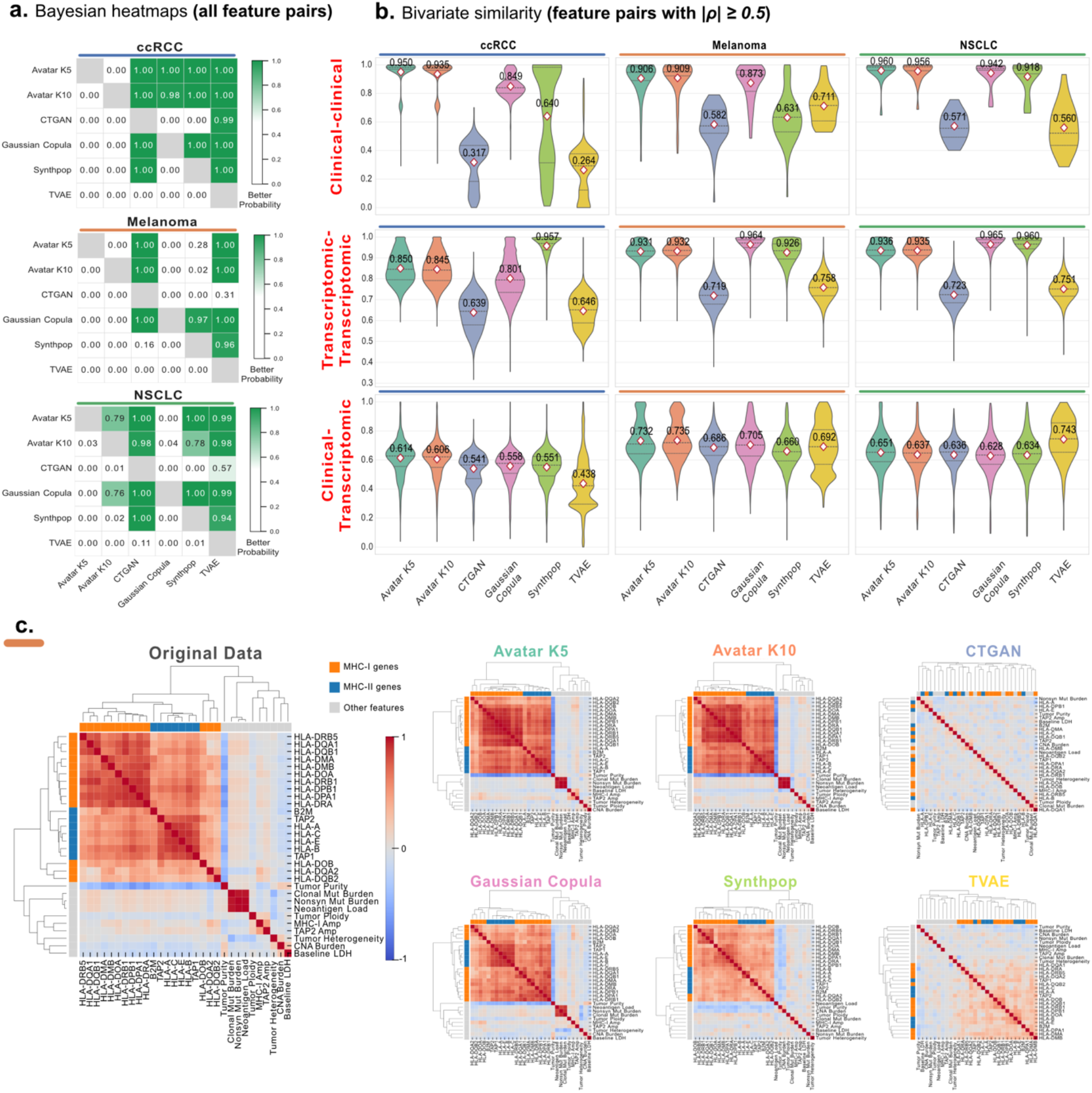
Comparative evaluation of bivariate similarity and structural relationship preservation across cancer cohorts. **a) Bayesian pairwise comparative probability heatmaps, derived from bivariate similarity scores of all feature pairs**. Heatmaps display pairwise “better probabilities” derived from Bayesian estimation. Each cell corresponds *P*(*row* > *column*) to representing the posterior probability that the SDG method in the row achieves superior bivariate similarity compared with the method in the column for the specified cohort. Values range from 0 to 1; darker green indicates stronger evidence of superior preservation of pairwise relationships. **b) Violin plot distribution of pairwise similarity for strongly correlated feature pairs (**|***ρ***| ≥ **0. 5)**. Violin plots depict the distribution of bivariate similarity scores for feature pairs with high correlation in the original data (|*ρ*| ≥ 0.5), stratified by clinical-clinical, transcriptomic-transcriptomic, and clinical-transcriptomic associations across the ccRCC, melanoma, and NSCLC cohorts. Each violin represents the distribution of similarity scores across selected feature pairs, with central markers indicating mean values. Higher and more compact distributions reflect improved preservation of strong inter-variable associations. **c) Hierarchical clustering of feature correlation matrices in the melanoma cohort (orange strip)**. Correlation heatmaps with hierarchical clustering depict the structural organization of feature-feature associations in the melanoma dataset. The “Original data” panel presents the correlation matrix of the real data and reveals a grouping immune-related genes according to MHC class I and II, a separate cluster of mutation and neoantigen load features, and a negative association between tumor purity and immune infiltration. Subsequent panels display the reconstructed correlation matrices generated by each SDG method.

To assess whether the statistical fidelity reflected biologically meaningful structure, following Liu et al, we performed hierarchical clustering of correlation matrices from the melanoma dataset (Figure 3c and Figure S4). The original data displayed clear biological organization, including immune-related clusters with MHC class I and II substructures, a separate cluster of mutation and neoantigen load features, and a negative association between tumor purity and immune infiltration. Avatar (K5/K10) and Gaussian Copula largely recapitulated these structures followed by Synthpop, whereas CTGAN and TVAE produced noisier or overly smoothed patterns. Together, these findings indicate that accurate reproduction of marginal distributions does not guarantee preservation of higher-order relationships, underscoring the need to evaluate bivariate structure and downstream biological utility.

### 2.3 Biological utility

Preserving biologically and clinically meaningful signals is arguably the most critical requirement for synthetic transcriptomic data intended for oncology research, yet it remains rarely evaluated in existing benchmarks. Here, we assessed concordance between original and synthetic datasets across six downstream bioinformatic tasks that represent standard analytical workflows in transcriptomic cancer studies: differential gene expression (DGE), gene-set enrichment analysis (GSEA), single-sample gene-set expression analysis (ssGSEA), cell type deconvolution, survival analysis, and machine learning-based modeling. We re-implemented bioinformatic findings from the original studies to test whether known biology could be rediscovered in synthetic data. Evaluation scores from biological utility metrics applied to downstream bioinformatic analyses are summarized in Table 3 and detailed in the Methods section.

**Table 3.**
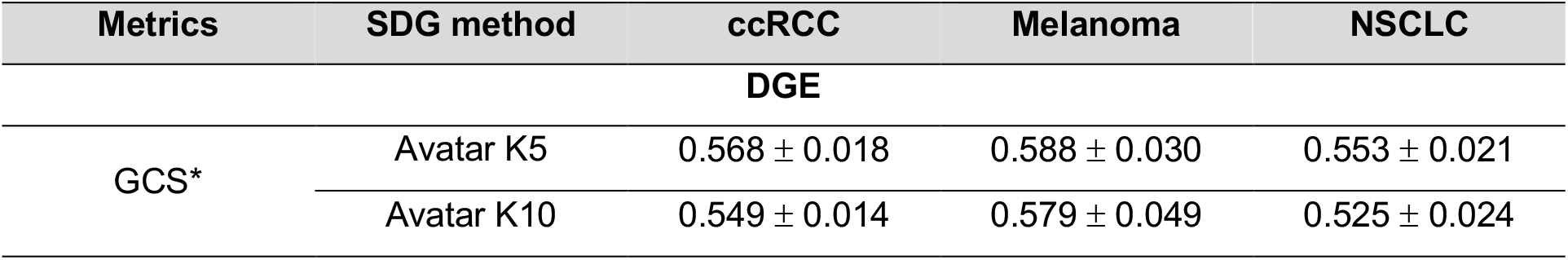
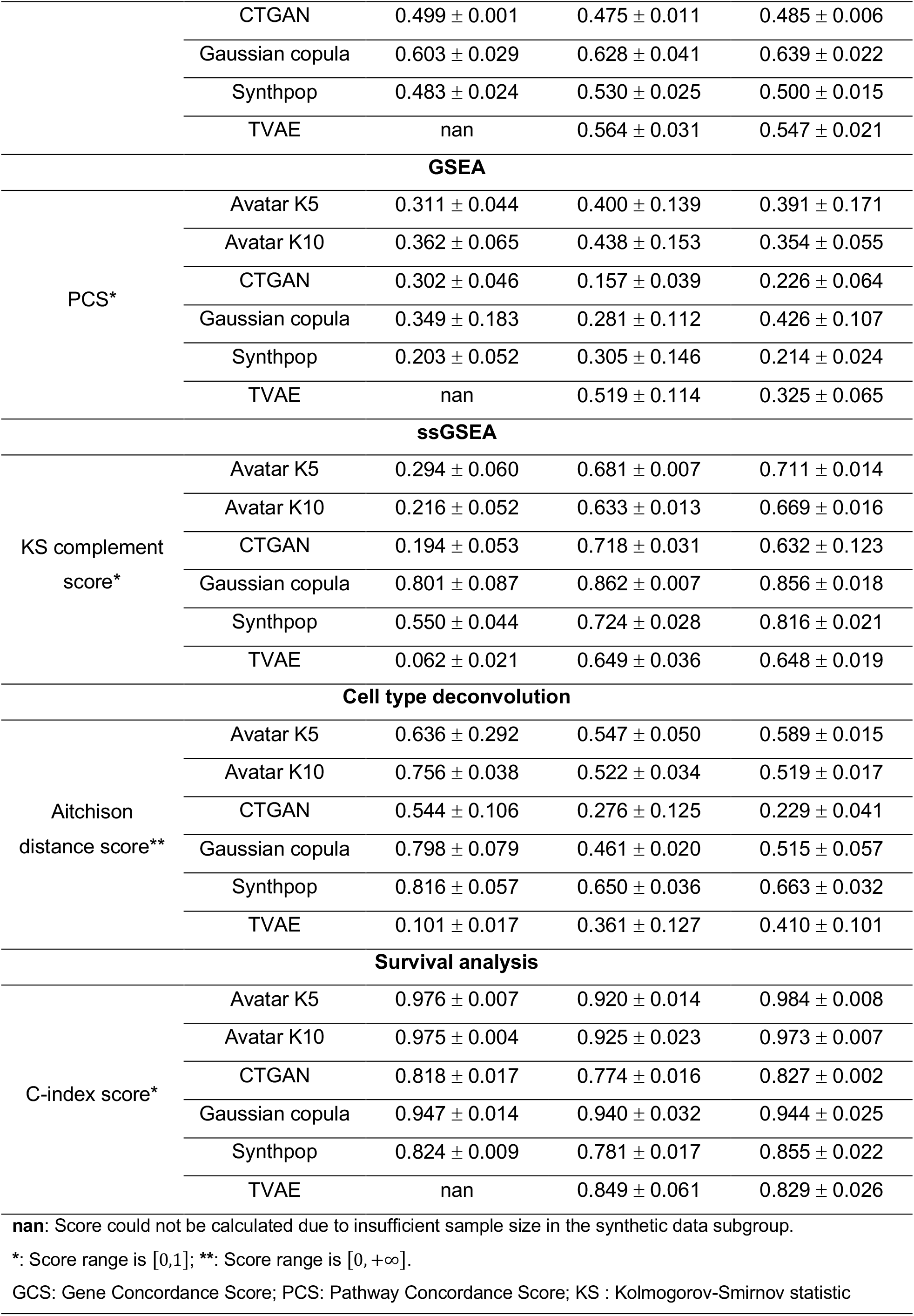
Biological utility assessment of synthetic datasets across three cancer cohorts based on specialized transcriptomic and clinical analysis scores. Higher scores indicate a stronger ability of the synthetic data to mirror the original data.

Identification of differentially expressed genes whose abundance levels differ systematically between biological conditions constitutes one of the most fundamental bioinformatic analyses in transcriptomic studies. Performance was evaluated via the Gene Concordance Score (GCS), which quantifies the weighted proportion of genes for which synthetic data preserves both directionality and statistical significance of the original data (Methods). Gaussian Copula consistently achieved the highest GCS across all cohorts, as confirmed by Bayesian estimation over five replicates (Figure 4a and Tables S7, S8, and S9); Spearman correlations of gene-wise *log*_2_*FC* values followed the same ordering (Figure S8 and Table S10). Biological rediscovery on synthetic data revealed selective patterns: higher angiogenesis score associated with kidney cancer samples harboring the PBRM1 mutation (Figure 4b, *P* < 0.01), MHC class II HLA gene upregulation in Melanoma responders-though with attenuation in Avatar K10/Gaussian Copula and inflated false positives in Avatar K5/TVAE (Figure 4c), and immunoproteasome enrichment (PSME1, PSME2, PSMB8, PSMB9, and PSMB10) over canonical IFN-*γ* targets and proteasome components in NSCLC were successfully recovered by Avatar (K5/K10) and Gaussian Copula (Figure 4d, *P* < 0.01). On the contrary, these biological patterns could not be recovered from synthetic data generated by the CTGAN and TVAE methods. Results from additional replicates are provided in the Supplementary Data (Figures S9, S10, and S11).

**Figure 4.**
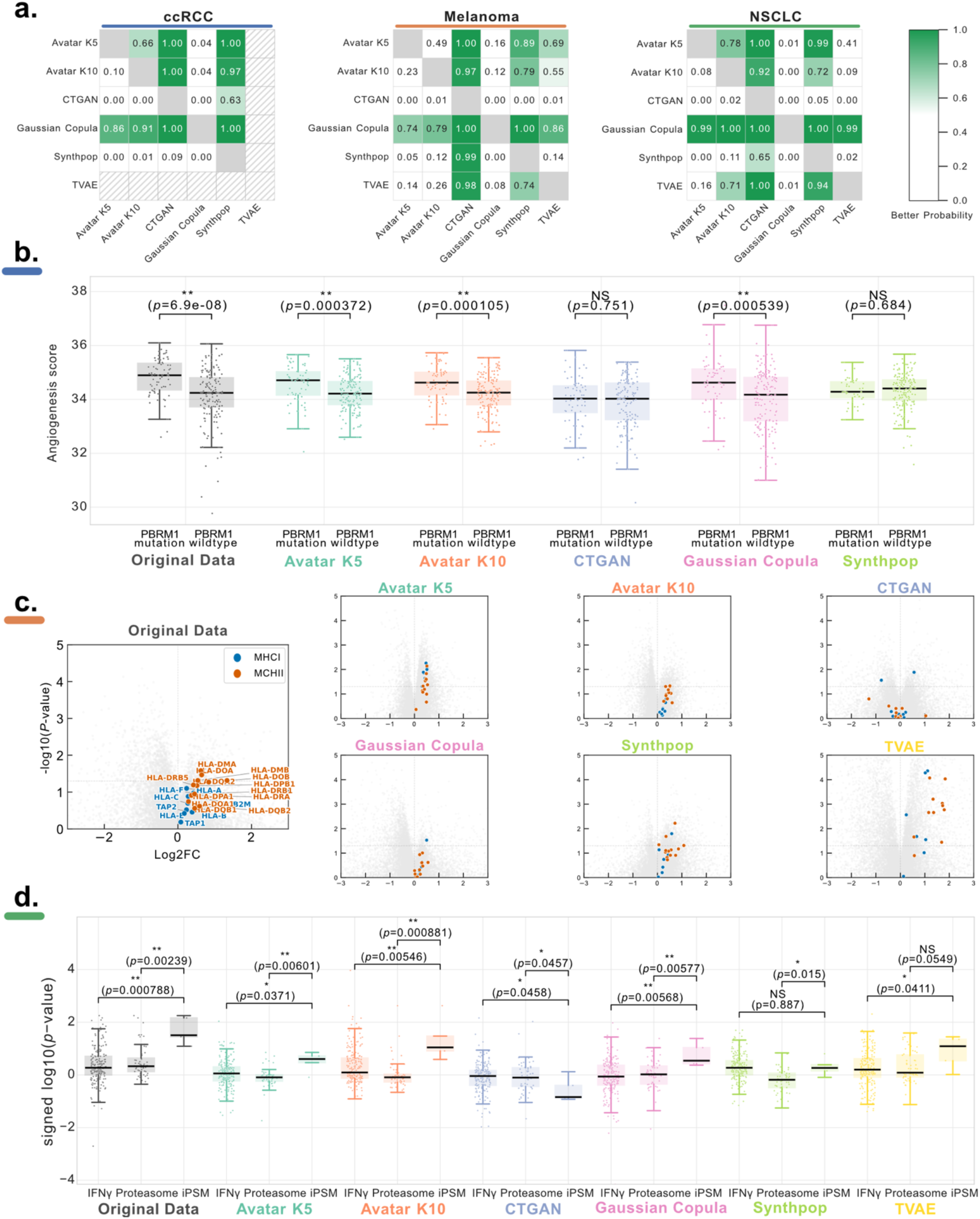
Preservation of differentially expressed genes and downstream biological re-discoveries in synthetic transcriptomic data. **a) Bayesian comparison of GCS**. Pairwise probability heatmaps display the posterior probability *P*(*row* > *column*) that a SDG method in a given row achieves a higher GCS than the method in the corresponding column. GCS quantifies the weighted proportion agreement between original and synthetic datasets in both regulation direction (sign of log_2_ *FC*) and statistical significance of individual gene. **b) PBRM1 mutation-associated angiogenesis signal in ccRCC (blue strip)**. Angiogenesis scores stratified by PBRM1 mutation status in patients are shown. In the original data, higher angiogenesis was significantly associated with PBRM1 mutated cancer samples. Statistical significance was assessed using a two-sided Wilcoxon rank-sum test. **c) Preservation of responder-associated MHC gene signatures in melanoma (orange strip)**. Volcano plots compare responders and non-responders with highlighted points corresponding to MHC class I and MHC class II HLA genes. The original data showed an upregulation of MHC class II genes in responders. **d) Re-discovery of immunoproteasome enrichment in NSCLC (green strip)**. Signed − log_10_(*p* − *value*) distributions compare IFN-*γ* response genes, immunoproteasome (iPSM) components, and canonical proteasome genes between responders and non-responders. The original hierarchical enrichment pattern is characterized by the upregulation of iPSM relative to the IFN-*γ* targets and general proteasome components. Statistical significance was assessed using a two-sided Wilcoxon rank-sum test. Shown panels represent one representative replicate; results from additional replicates are provided in Figures S9, S10, and S11 in the Supplementary Data.

Interpretation of gene-level information at the pathway level with gene set enrichment analysis (GSEA) allow to test whether predefined gene sets show coordinated expression changes between conditions. Similarly to gene-level, performance was evaluated using the Pathway Concordance Score (PCS) alongside Bayesian comparison (Figure 5a, Figures S12, S13, and S14 and Methods section). We found that no single method dominated pathway-level concordance across cohorts (Table 3, Figure 5c, and Tables S11, S12, and S13). Immune-related pathways (IFN-*γ* response, allograft rejection, complement, inflammatory response, and IL6-JAK-STAT3 signaling) originally enriched in treatment responders in melanoma were broadly recovered by most methods except CTGAN (Figure 5b), though reproducibility across replicates varied substantially. In ccRCC, Avatar (K5/K10) and Gaussian Copula reproducibly recapitulated downregulated pathways in 9p21.3 deletion patients compared to controls, though upregulated pathways were less consistently preserved across replicates (Figure 5d). Furthermore, the treatment-stratified IFN-*γ*/IFN-*α* enrichment pattern in ipilimumab-treated but not ipilimumab-naive melanoma patients, were robustly reproduced by Gaussian Copula and partially by Avatar (K5/K10) and Synthpop methods (Figure 5e). This highlights that single-run assessments can overestimate synthetic data quality; reproducibility across random seeds is essential for reliable biological inference.

**Figure 5.**
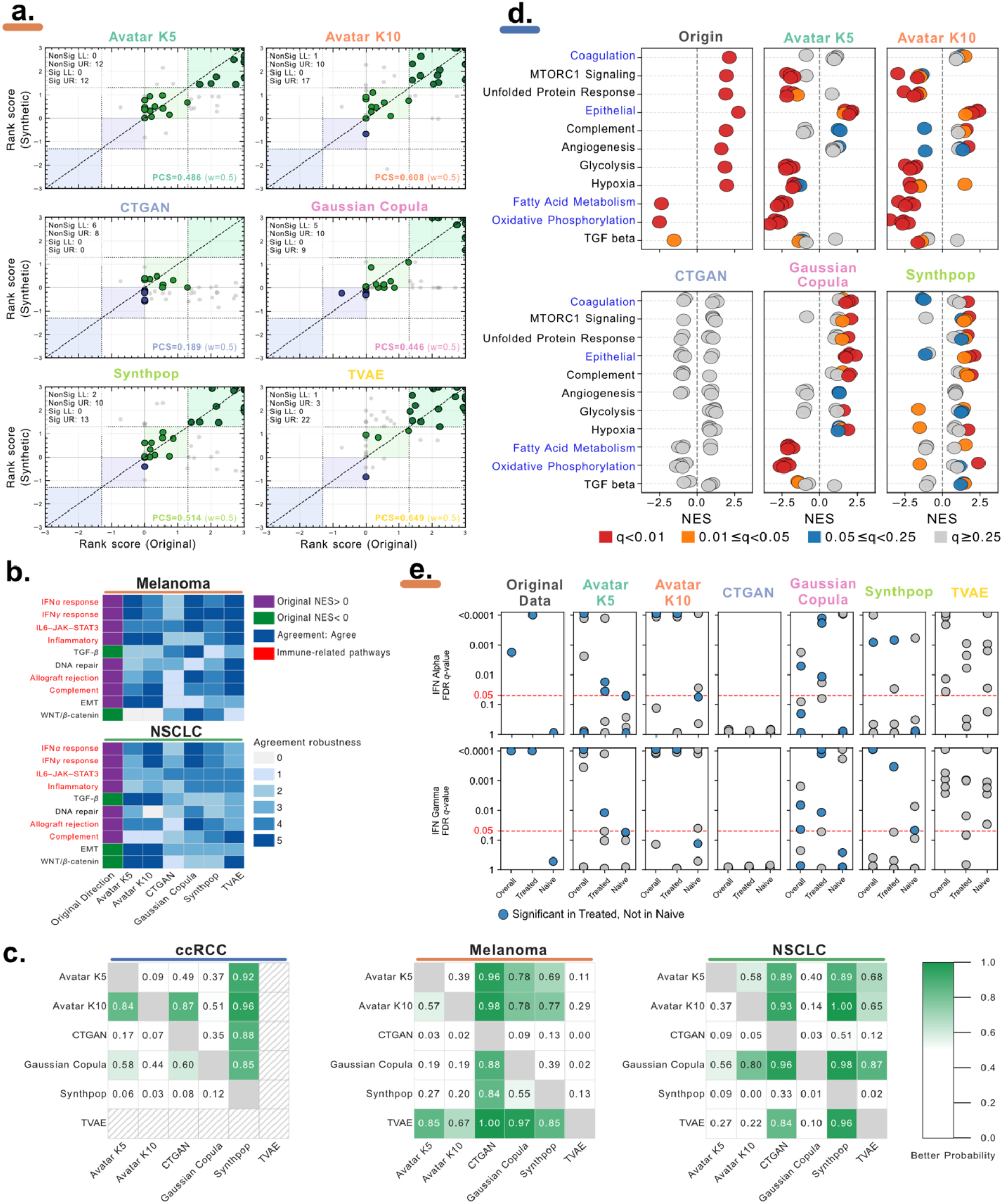
The concordance of pathway-level signals and biomarker recovery across synthetic datasets. **a) PCS scatter plots of Melanoma cohort**. These scatter plots compare pathway rank score between the original data and synthetic datasets of each SDG method. Rank scores combine the sign of the NES and the False discovery rate (FDR), *Q* value. Pathways located in the upper-right (UR, light green) and lower-left (LL, light purple) quadrants indicate concordant regulation direction and the level of significance (FDR, *Q* < 0.05). **b) Directional agreement of key cancer pathways**. Heatmaps display enrichment direction and agreement patterns for selected functional pathways in Melanoma and NSCL. Immune-related pathways enriched in responders (IFN-*γ* response, IFN-*α* response, Allograft rejection, Complement, Inflammatory response, IL6-JAK-STAT3 signaling) were largely recapitulated by Avatar (K5/K10), Gaussian Copula, Synthpop, and TVAE, but inconsistently by CTGAN. Reproducibility across random-seed replicates is summarized alongside. **c) Bayesian comparison of PCS**. Pairwise probability heatmaps display the posterior probability *P*(*row* > *column*) that an SDG method in a given row achieves a higher PCS than the method in the corresponding column. **d) GSEA result of 9p21.3 deletion in ccRCC cohort (blue strip)**. Bubble plots show NES values for Hallmark pathways in the original and synthetic datasets. Color indicates statistical significance. Avatar (K5/K10) and Gaussian Copula reproducibly captured downregulated pathways (fatty acid metabolism, oxidative phosphorylation), whereas upregulated signals showed greater variability. Gaussian Copula demonstrated the highest cross-replicate stability. **e) Treatment-stratified IFN response in the melanoma cohort (orange strip)**. Dot plots present FDR *q*-values for IFN-*γ* and IFN-*α* response pathways stratified by prior ipilimumab exposure.

The assignment of a pathway activity score to each individual sample by single-sample GSEA (ssGSEA) method enables patient-level characterization of biological programs. Gaussian Copula achieved the highest KS-Complement scores for ssGSEA NES distributions across all cohorts, with Bayesian posterior probabilities exceeding 87%, followed by Synthpop (Figure 6a-b, Figure S15, and Tables S14, S15, and S16). In ccRCC, biological signals such as PBRM1-associated IL6-JAK-STAT3 reduction alongside estrogen response, apoptosis, allograft rejection, and UV response pathways were robustly reproduced by Gaussian Copula, and to a lesser extent by Avatar, while other methods showed pronounced inter-replicate variability (Figure 6c). In Melanoma, the subgroup-specific pattern of higher MHC-II ssGSEA scores in ipilimumab-treated responders was recapitulated only by Avatar K10 and Gaussian Copula, although recovery was limited to a single replicate per method (Figure S16).

**Figure 6.**
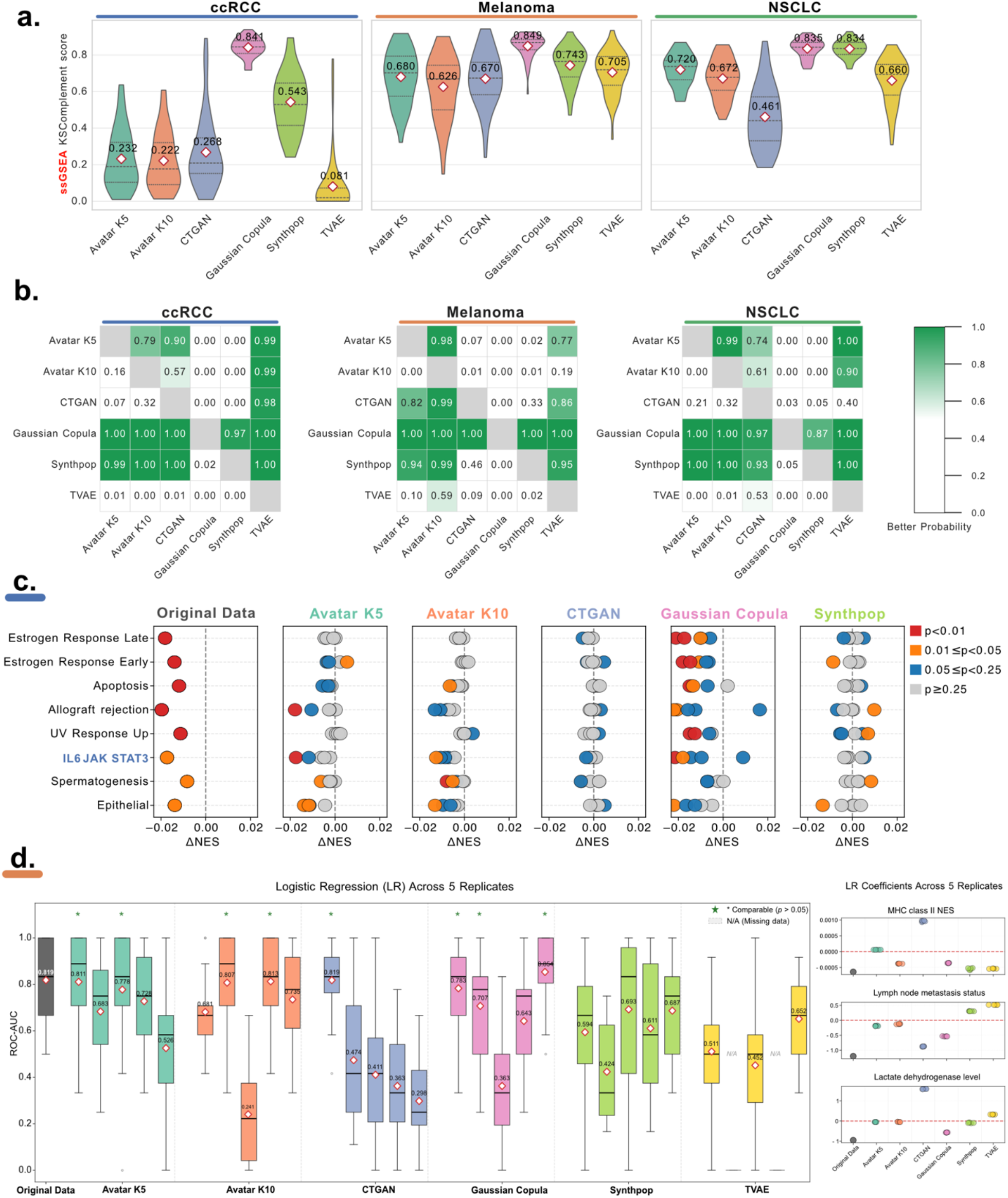
The concordance of single-sample pathway-level signal, biomarker recovery, and predictive transferability in synthetic data. **a) Distribution of KS-Complement scores of 50 Hallmark pathways**. Violin plots summarize pathway-level KS-Complement values across datasets. Higher values reflect greater similarity of single-sample GSEA (ssGSEA) score distributions between original and synthetic data. **b) Bayesian comparison of KS-Complement**. Pairwise probability heatmaps display the posterior probability P(row>column) that a SDG method in a given row achieves a higher pathway-wise KS-Complement scores than the method in the corresponding column. **c) Association between ssGSEA signals and PBRM1 phenotype in the ccRCC dataset (blue strip)**. Bubble plots compare differential NES between PBRM1 mutation patients and PBRM1 wildtype patients in original and synthetic datasets. Bubble color denotes *P* value thresholds (Wilcoxon rank-sum test). PBRM1 mutations were associated with reduced IL6-JAK-STAT3 signaling in the original cohort. Additional pathways (estrogen response, apoptosis, allograft rejection, UV response) are shown for comparison. **d) Predictive modeling transferability in Melanoma cohort (orange strip)**. Boxplots depict cross-validated ROC-AUC for logistic regression (LR) models trained on synthetic data replicates and evaluated on held-out original data (5-fold cross-validation, repeated three times). The original data benchmark is shown for reference. Colored dots indicate denote statistically non-inferior performance versus the original model (Wilcoxon signed-rank test *P* > 0.05). Right panels display learned coefficients for MHC class II score, lymph node metastasis status, and LDH level. **Results for additional classifiers and replicates are provided in Figure S17**.

In the original melanoma study, Liu et al. developed a logistic regression (LR) model to predict progression status under combined anti-PD1 and anti-CTLA-4 therapy. We subsequently trained classifiers (LR, random forest, MLP) exclusively on synthetic data and evaluated on held-out original data. Gaussian Copula synthetic dataset achieved performance comparable to real-data baselines (Wilcoxon signed-rank test *P* > 0.05) and was the most consistent across replicates (Figure 6d left panel and Figure S17). Furthermore, Gaussian Copula’s LR models preserved the coefficient directionality for prognostic features MHC class II score, lymph node metastasis status and LDH level (Figure 6d right panel). Taken together, these findings highlight an important limitation: high marginal pathway-level agreement does not guarantee reliable biological interpretation or predictive transferability, as substantial inter-replicate variability can still arise in downstream analyses.

A cell type deconvolution method estimates the relative abundance of immune and stromal cell populations from bulk transcriptomic profiles, providing a proxy for the tumor microenvironment composition. The global immune landscape was predicted using CIBERTSORTx and its default LM22 immune cell type signatures.^26^ Its average concordance level with original data, measured by Aitchison distance,^28^ was highest in ccRCC and progressively lower in Melanoma and NSCLC across all methods (Table 3, Figure S18, and Tables S17, S18, and S19). Recovery of biologically meaningful immune contrasts was cohort and signal-dependent. In ccRCC, Avatar (K5/K10) and Gaussian Copula reconstructed immune contrasts between infiltrated and excluded/desert tumors, with reproducible recovery at a relaxed significance level (Wilcoxon rank-sum test, False discovery rate (FDR) *Q* < 0.25) confined to cell types with robust signals in the original data (Figure 7a). Interestingly, in melanoma and NSCLC, differential analysis of LM22 immune cell types in the original data revealed no significant cell-type differences; accordingly, no SDG method produced stable results under these settings (Figure S19). For single-cell-derived signatures collected from the melanoma publication,^19,29^ most methods except CTGAN preserved directional enrichment in responders, with a notable statistical significance attenuation from the original data to synthetic datasets across cell types, signatures, and cohorts (Figure 7b). TVAE nominally recovered several signatures but should be interpreted cautiously, given its extreme responder-to-progressor ratio distortion across replicates (10.1%-692.9% vs. 83.9% in the original), consistent with its poor univariate and bivariate fidelity. In NSCLC, the higher-order co-regulation structure of wound-healing/immunosuppressive and immune activation/exhaustion clusters was preserved only by Gaussian Copula and Avatar K5, partially by Avatar K10, and largely lost in the remaining methods (Figure 7c). The synthetic heatmaps of other replicates are provided in the Supplementary Data (Figure S20). A general trend emerged: compositional immune signals were reproducible only when the original effect size was large, and statistical significance was systematically attenuated in synthetic data, irrespective of the SDG method.

**Figure 7.**
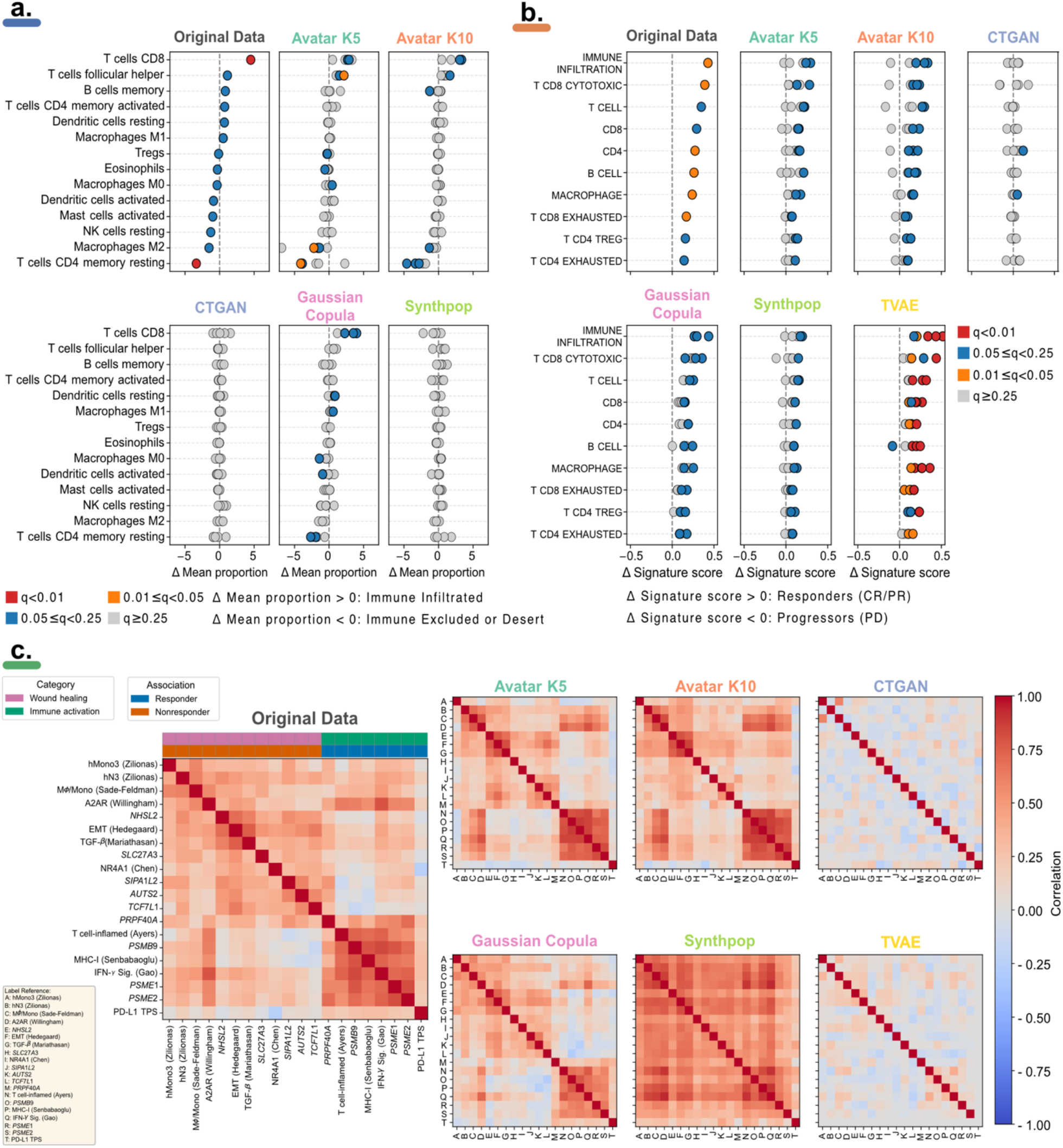
Preservation of immune cell composition and higher-order biological signatures in synthetic data. **a) Differential LM22 immune cell proportions in ccRCC (blue strip)**. Dot plots compare mean differences in immune cell fractions inferred by CIBERSORTx between immune-infiltrated and immune-excluded/desert tumors across the original and synthetic datasets generated by each SDG method. Colors indicate statistical significance based on a two-sided Wilcoxon rank-sum test with FDR adjustment. **b) Recovery of single-cell-derived immune signatures in Melanoma (orange strip)**. Dot plots show the mean differences of signature score between responders and progressors for immune programs derived from single-cell RNA-seq. The colors represent FDR-adjusted significance thresholds from a two-sided Wilcoxon rank-sum test (same color scheme as in panel a). **c) Biological signature correlation matrices in the NSCLC cohort (green strip)**. Correlation heatmaps display pairwise Pearson correlations among signatures in the original data and each synthetic dataset.

The color scale ranges from negative (blue) to positive (red) correlations. Clustering highlights two principal modules: a resistance-associated wound-healing/immunosuppressive, stromal cluster (C1) and a response-associated immune activation/exhaustion cluster (C2). The synthetic heatmaps of other replicates are provided in Figure S20 in the Supplementary Data.

Survival analysis evaluates the association between clinical or molecular features and patient outcomes, representing the most direct measure of clinical relevance in oncology datasets. Survival similarity was assessed using a C-index score derived from original and synthetic datasets, complemented by log-rank tests on Kaplan-Meier survival curves (Methods, Figures S22, S23, S24, S25, S26, and S27). Global survival discrimination was well preserved by the best-performing methods (Avatar K5/K10 and Gaussian Copula), with C-index scores above 0.9 confirmed by Bayesian estimation (Table 3, Figure S21, and Tables S20, S21, and S22).

Clinically interpretable biomarker stratification, including PBRM1-associated survival benefit in ccRCC (Figure 8a, *P* < 0.05), classifier-based risk score stratification (see ssGSEA section) in melanoma (Figure 8b, Figure S29 *P* < 0.05), and macrophage infiltration-associated poor prognosis in PD-L1-high patients in NSCLC (Figure 8c, *P* < 0.05) were investigated across all methods. Results from additional replicates are provided in the Supplementary Data (Figures S28, S29, S30, and S31). The recovery of downstream clinical stratification patterns proved more demanding and method-dependent, with only Gaussian Copula consistently achieving significant patient classification across all three cohorts.

**Figure 8.**
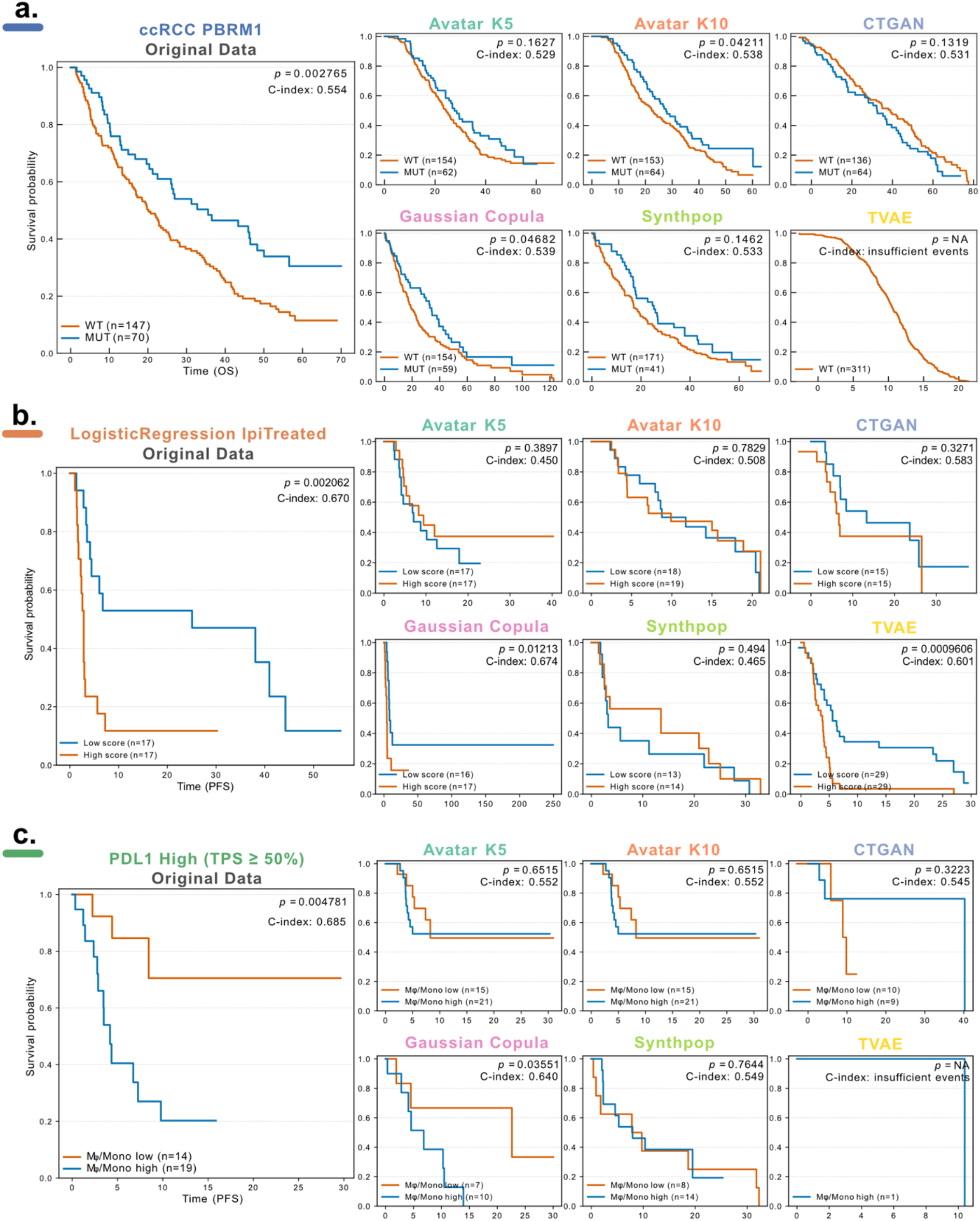
Survival signal preservation and clinical risk stratification in synthetic cohorts. **a) PBRM1-associated overall survival in ccRCC (blue strip)**. Kaplan-Meier curves compare overall survival between PBRM1-mutant (blue) and wild-type (orange) groups in the original dataset and across synthetic cohorts generated by each SDG method. The original survival advantage of PBRM1-mutant patients is reproduced in selected synthetic datasets, with log-rank *P* values C-indices reported in each panel. **b) Preservation of survival stratification based on a predictive model in the ipilimumab-treated Melanoma cohort (orange strip)**. Kaplan-Meier curves of PFS stratified by predicted risk groups (low risk: blue; high risk: orange) derived from a logistic regression model classifying progressors versus non-progressors. The model was retrained on each synthetic dataset, and predicted risk scores were used for survival stratification. Log-rank *P* values and C-indices are reported in each panel. **c) Patient stratification using a macrophage/monocyte signature in the PD-L1 high NSCLC cohort (green strip)**. Kaplan-Meier curves evaluate progression-free survival differences between tumors with low (orange) versus high (blue) macrophage/monocyte signature scores within the PD-L1 high (TPS ≥ 50%) subgroup. Recovery of the adverse prognostic association observed in the original dataset is assessed across SDG methods. Log-rank *P* values and C-indices are shown in each panel. Shown panels represent one representative replicate; results from additional replicates are provided in Figures S28, S29, S30, and S31 in the Supplementary Data.

### 2.4 Privacy risk evaluation

Beyond statistical fidelity and biological utility, privacy preservation is a critical requirement for synthetic clinical transcriptomic data, given the high sensitivity of this patient-level information. To quantify privacy risk, we applied the Anonymeter framework to simulate three canonical attack scenarios: singling out, linkability, and attribute inference.^18^ Results were summarized into an overall privacy score, where higher values indicate stronger privacy preservation (Table 4 and Methods section).

**Table 4.**
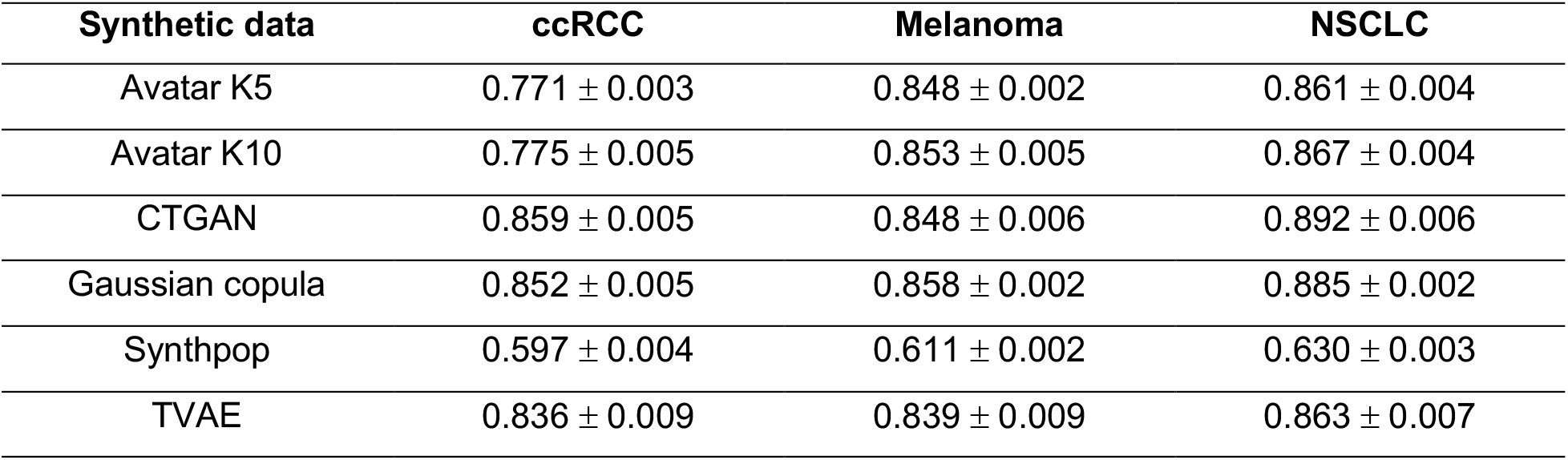
Overall privacy score integrating singling out risk, linkability and attribute inference risk of 6 synthetic dataset across three cancer cohorts. Higher scores indicate a stronger ability of the synthetic data to preserve the privacy.

| Synthetic data | ccRCC | Melanoma | NSCLC |
| --- | --- | --- | --- |
| Avatar K5 | 0.771 ± 0.003 | 0.848 ± 0.002 | 0.861 ± 0.004 |
| Avatar K10 | 0.775 ± 0.005 | 0.853 ± 0.005 | 0.867 ± 0.004 |
| CTGAN | 0.859 ± 0.005 | 0.848 ± 0.006 | 0.892 ± 0.006 |
| Gaussian copula | 0.852 ± 0.005 | 0.858 ± 0.002 | 0.885 ± 0.002 |
| Synthpop | 0.597 ± 0.004 | 0.611 ± 0.002 | 0.630 ± 0.003 |
| TVAE | 0.836 ± 0.009 | 0.839 ± 0.009 | 0.863 ± 0.007 |

For univariate singling out, most methods produced attack success rates at or below the naive random baseline, corresponding to negligible risk. Synthpop was a clear exception, exhibiting singling-out risk close to 1.0 across all three cohorts, consistent with its near-perfect marginal distribution reproduction (Figure 9a). Multivariate singling-out risk decreased with predicate dimensionality across all methods, as expected, yet Synthpop retained the highest risk in all cohorts (Figure 9b). Linkability risk was consistently low across all methods and clinical cohorts indicating that synthetic records cannot be reliably re-identified through attribute bridging, even when large transcriptomic feature subsets were used (Figure 9c).

**Figure 9.**
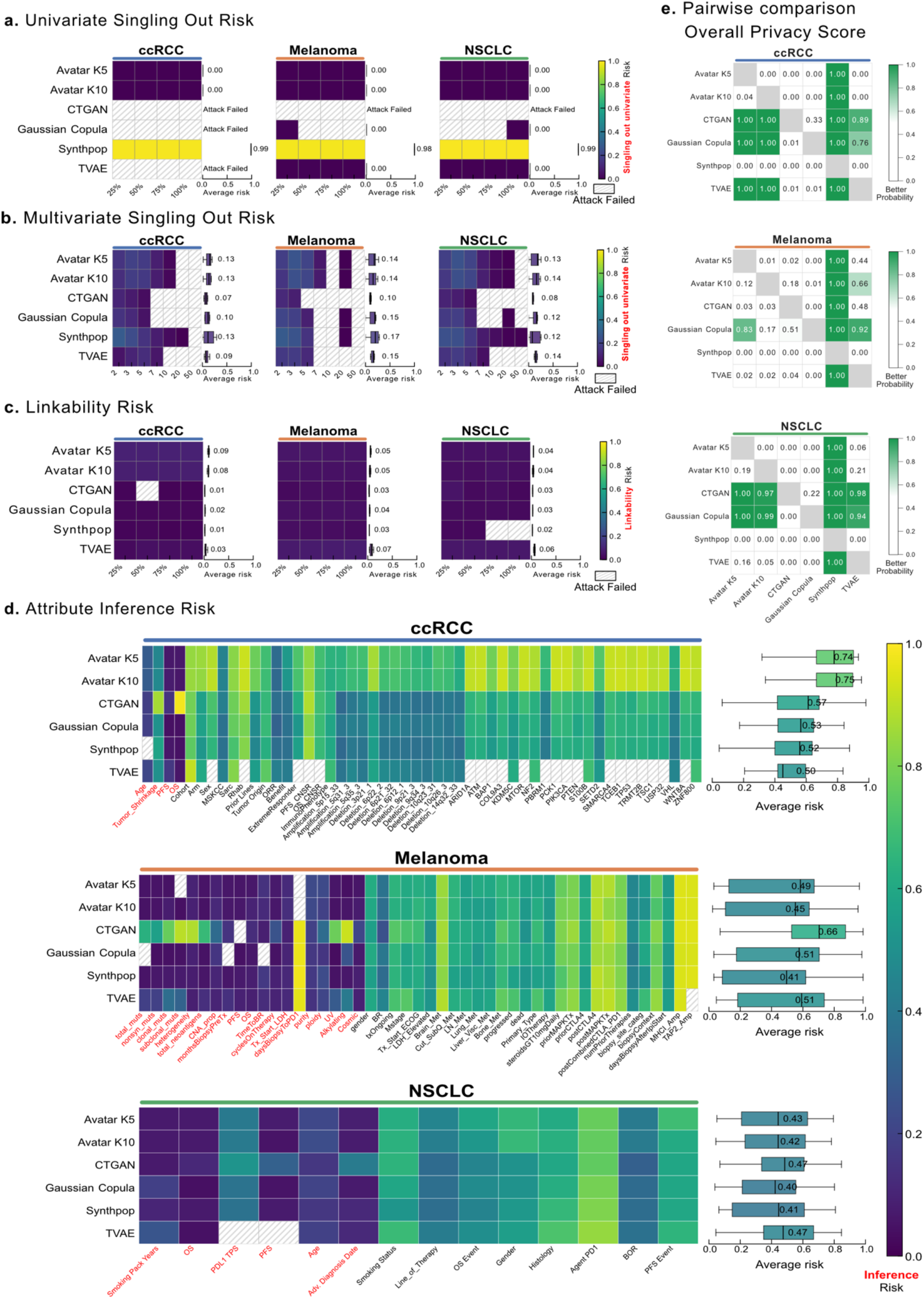
Privacy risk evaluation of synthetic datasets utilizing the Anonymeter framework. **a) Univariate singling-out risk**. Heatmaps display the singling out risk that predicates constructed from synthetic data can uniquely identify a record in the original dataset using random feature subsets (25-100%). Rows correspond to SDG methods. Each cell represents the mean risk across five replicates for a given attack scenario. Boxplots adjacent to each heatmap summarize the average risk per method across all scenarios and replicates.. **b) Multivariate singling out risk**. Heatmaps show singling out risk when predicates combine increasing numbers of attributes. As predicate dimensionality increases, attack effectiveness decreases across methods, reflected by lower risk intensities. Boxplots adjacent to each heatmap summarize the average risk per method across all scenarios and replicates. **c) Linkability risk**. Heatmaps represent the linkability risk that synthetic records enable re-linking of disjoint attribute groups belonging to the same original individual. Boxplots adjacent to each heatmap summarize the average risk per method across all scenarios and replicates. **d) Attribute inference risk**. Heatmaps illustrate the risk of inferring hidden “secret” attributes (e.g., clinical outcomes) from auxiliary transcriptomic features. Boxplots to the right summarize mean inference risk per method across scenarios and replicates. Color intensity indicates risk magnitude, with warmer colors denoting higher risk. **e) Bayesian comparison of overall privacy score**. Pairwise probability heatmaps show the posterior probability *P*(*row* > *column*) that the SDG method in a given row achieves a higher aggregated privacy score than the method in the corresponding column.

Attribute inference represented the most substantial residual privacy risk, with success rates ranging from 0.4 to 0.75 depending on the dataset and method (Figure 9d). Risk levels varied across cohorts and methods, with no single approach performing uniformly well. Notably, numerical clinical attributes consistently showed lower inference risk than categorical ones, a pattern attributable to Anonymeter’s evaluation design: categorical attributes are assessed by exact-match classification, whereas numerical attributes require regression-based inference under a relative error threshold.

Overall, aggregated privacy scores and Bayesian comparative analysis identified CTGAN and Gaussian Copula as the best-performing methods for privacy preservation, while Synthpop ranked last across all cohorts, driven by its extremely high singling-out risk (Figure 9e and Tables S23, S24, and S25).

### 2.5 Computational resources

Computational resource requirements critically shape the real-world feasibility of synthetic data generation in biomedical settings. Table 5 reports hardware configuration, peak memory usage, execution time, energy consumption, output size, and setup complexity for each evaluated SDG method.

**Table 5.**
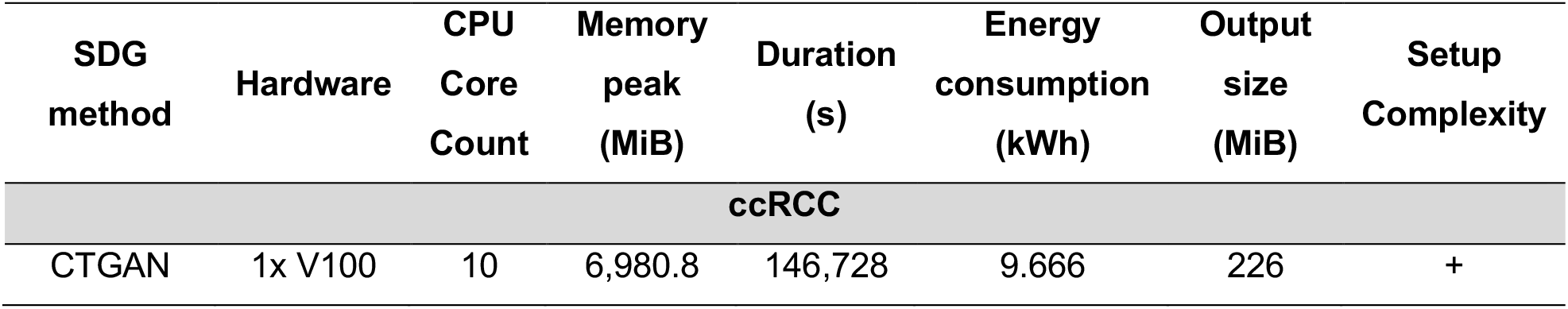
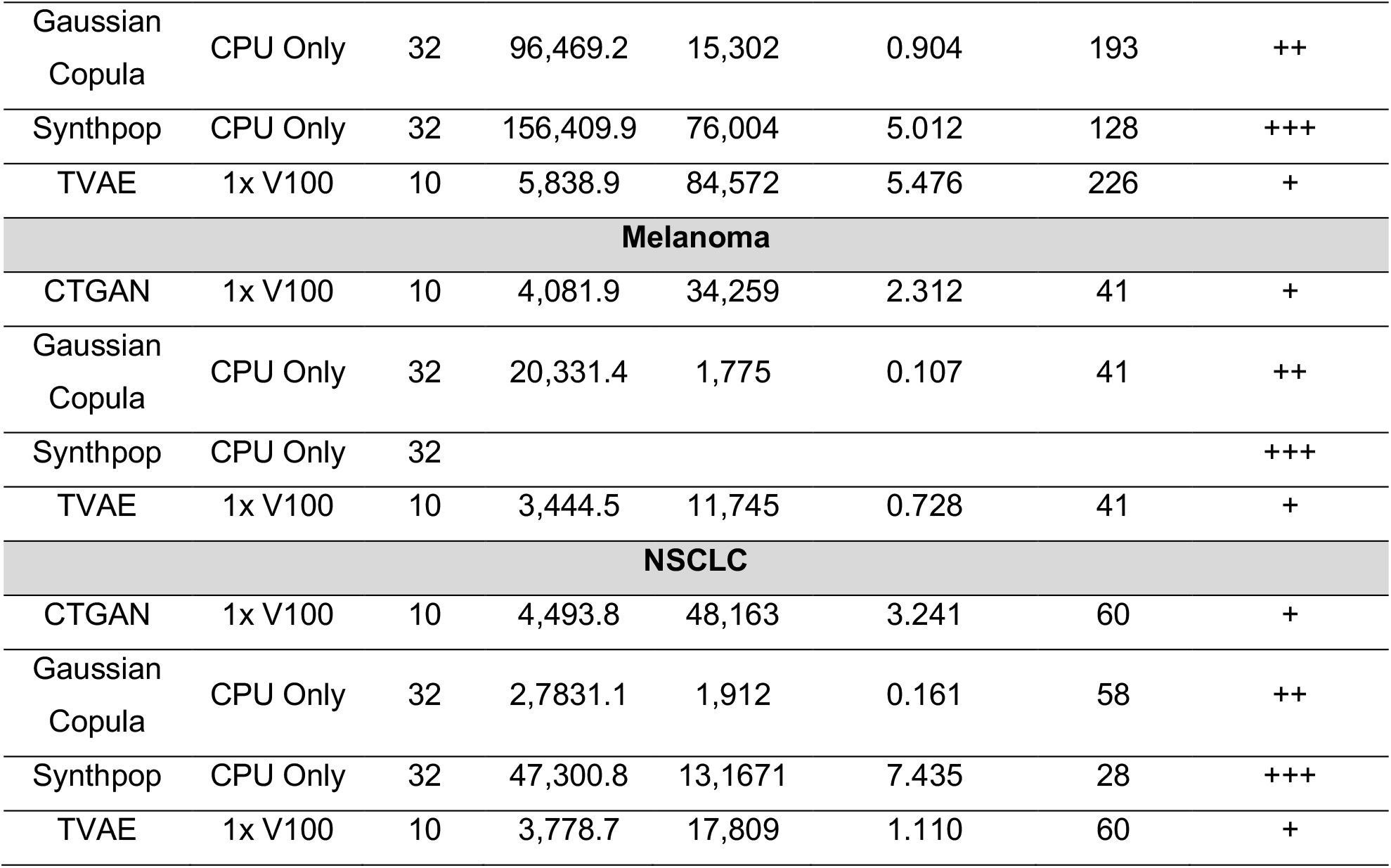
Computational cost and deployment complexity of SDG methods across oncology datasets.

Gaussian Copula was the least resource-intensive method overall: it ran on CPU-only infrastructure, completed within 0.5 to 4 hours depending on cohort size, and consumed less than 1 kWh. Peak memory reached approximately 96 GiB in ccRCC, which requires a well-provisioned server but remains practical in standard HPC environments. Setup complexity was moderate, primarily due to the need for careful encoding of categorical variables. CTGAN and TVAE required GPU acceleration and substantially longer runtimes (3 to 40 hours), with energy consumption reaching approximately 10 kWh in ccRCC. These requirements may limit deployment in resource-constrained settings. Synthpop presented an unusual profile: despite running on CPU only, it required the highest peak memory of all methods (up to approximately 156 GiB in ccRCC) and runtimes of up to 36 hours for larger cohorts. Crucially, these figures were obtained under a carefully configured setup in which predictor matrices were manually constrained to limit variable interactions and reduce memory burden. In a default configuration, resource consumption would likely be higher. This customization also increases setup complexity considerably. Avatar is the only proprietary method evaluated. It can be accessed via a web interface or the Octopize-Avatar Python API. Still, the API requires partitioning the feature space into blocks, as the full data matrix cannot be processed simultaneously. This constraint increases setup complexity and may affect preservation of cross-block feature dependencies.

### 2.6 Meta-ranking methods

To provide an integrated comparison across all evaluated dimensions, we computed a rank-derived meta-score for each SDG method following the protocol of Yan et al.^30^ These meta-scores aggregate performance across statistical fidelity, biological utility, and privacy into a single summary ranking. Results are shown per clinical cohort and across the combined dataset in Figures 10a and 10b, respectively.

**Figure 10.**
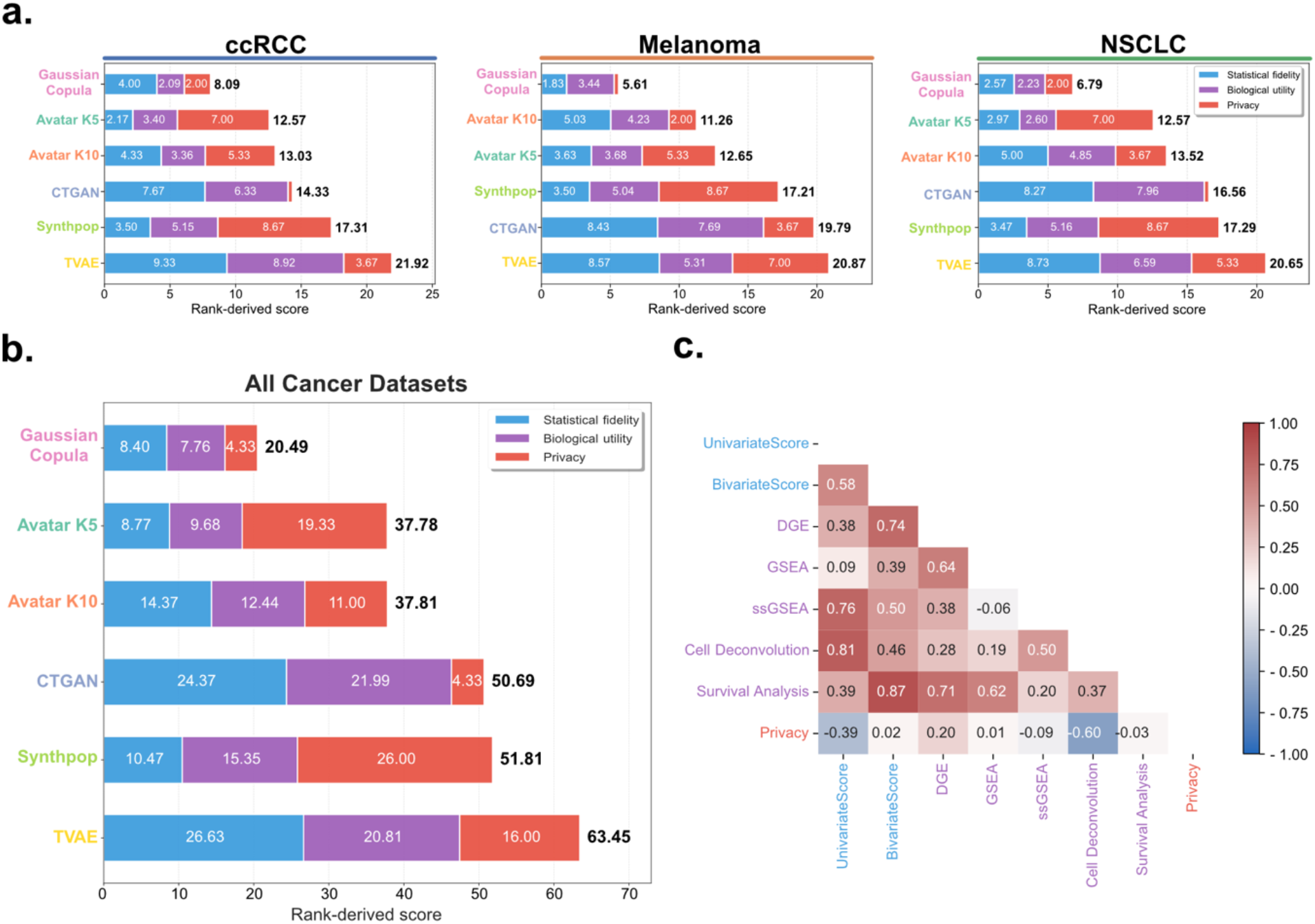
Rank-derived meta-score comparison and metric correlation analysis of SDG methods. **a) Meta-score comparison within each cancer cohort**. Stacked bar plots display the rank-derived meta-scores for each SDG method across ccRCC, melanoma, and NSCLC datasets. For each method, the total bar length represents the aggregated meta-score, computed by integrating three evaluation pillars: statistical fidelity, biological utility, and privacy. The three colored segments correspond to these pillars and indicate their respective contributions to the overall score. Because the metric is rank-derived, lower meta-scores indicate better overall performance. **b) Aggregated meta-score across all datasets**. Stacked bar plots summarize rank-derived meta-scores across all three cancer datasets. Each bar aggregates the three dimensions (statistical fidelity, biological utility, and privacy), represented by distinct color segments, into a composite meta-score. **c) Pairwise correlation of evaluation metrics**. Heatmap of Spearman correlation coefficients among all evaluation metrics across SDG methods and cancer datasets.

Gaussian Copula ranked first overall, both within individual cancer types and in the aggregated analysis, followed by Avatar K5 and K10. A consistent utility-privacy trade-off was observed: methods with stronger statistical fidelity and biological utility scores, notably Gaussian Copula and Avatar, tended to score lower on privacy, while CTGAN achieved higher privacy scores at the cost of lower utility. Synthpop presented a distinctive profile, reaching competitive scores in statistical fidelity while ranking near last overall, penalized by its poor singling-out performance.

To assess the complementarity of the individual metrics, we computed pairwise correlations across all methods and cohorts (Figure 10c). Several informative patterns emerged. The utility-privacy trade-off was clearly reflected in weak or negative correlations between privacy and most biological utility metrics. Among utility measures, GSEA and ssGSEA showed surprisingly weak mutual correlation despite both being pathway-based: GSEA correlated more strongly with DGE and survival analysis, consistent with its focus on population-level transcriptional effects, whereas ssGSEA correlated more closely with univariate scores and cell-type deconvolution metrics, reflecting its sample-level enrichment quantification. Univariate and bivariate scores were moderately correlated, indicating that bivariate analysis captures additional discriminative structure not fully reflected in marginal distribution similarity alone. Taken together, these findings reinforce the need for a multi-dimensional evaluation framework: no single metric provides a sufficient summary of synthetic data quality across the range of downstream use cases considered here.

## 3 Discussion

This study introduces SynOmicsBench (https://trinhthechuong.github.io/SynOmicsBench) as a unified benchmarking framework to systematically evaluate synthetic data generation (SDG) methods for clinical transcriptomic data across three oncology cohorts (ccRCC, Melanoma, and NSCLC).^3,19,20^ The framework integrates three complementary evaluation axes: statistical fidelity, biological utility, and privacy risk to assess whether current SDG approaches can address the statistical and structural challenges of large-scale gene expression data combined with heterogeneous clinical variables. A central design choice is to separate data quality into two distinct dimensions. Statistical fidelity quantifies distributional agreement at the level of marginal and joint feature distributions. Biological utility, by contrast, assesses whether results from established transcriptomic analyses (differential gene expression, pathway enrichment, immune deconvolution, and survival modeling) can be faithfully recovered from synthetic data. This separation is motivated by the observation that distributional similarity alone does not guarantee preservation of the structured, higher-order dependencies that underpin biological interpretation.

Beyond comparative evaluation, the benchmark provides a standardized, reproducible end-to-end pipeline spanning preprocessing, transcriptomic harmonization, multivariate imputation, model training adaptations, and downstream validation. Our framework is designed to be reusable and extensible, enabling the community to evaluate emerging SDG methods or to select approaches suited to specific biological and privacy requirements.

We benchmarked five methods spanning the main families of the SDG landscape: Synthpop (sequential conditional modeling),^22^ Gaussian Copula (parametric copula-based),^16^ CTGAN and TVAE (deep generative models),^21^ and Avatar (local-neighborhood swapping).^12^ No single algorithm excelled across all evaluation dimensions, highlighting inherent trade-offs between statistical fidelity, biological utility, and privacy preservation. Gaussian Copula consistently emerged as the strongest overall performer, achieving a balanced profile across dimensions and datasets, followed by Avatar method (Figure 10a, b). Conversely, deep generative models (TVAE and CTGAN) were computationally demanding and exhibited instability in clinical cohorts of a few hundred patients, manifested as structural degradation and distortion of responder-to-progressor ratios.

Globally, method performance was substantially shaped by dataset characteristics, particularly feature dimensionality and sample size (Table 1). In the very high-dimensional ccRCC cohort (>40,000 transcriptomic features for 311 patients), all methods showed reduced gene-wise fold-change concordance, underscoring the fundamental difficulty of modeling such spaces (Figure S8 and Table S10).

Existing benchmarks have focused almost exclusively on statistical fidelity, leaving the biological utility of synthetic transcriptomic data largely uncharacterized. A distinguishing value of this study is its emphasis on biological utility as a first-class evaluation criterion. By developing task-specific metrics aligned with established transcriptomic workflows and by re-implementing the biological analyses from original studies on synthetic data, this framework directly tests whether clinically and molecularly relevant signals can be rediscovered. This positions biological utility not as an auxiliary consideration but as a core requirement for evaluating SDG quality in translational settings. Indeed, the divergence between statistical fidelity and biological utility emerged as a key finding. Synthpop achieved the highest univariate similarity across all cohorts (Table 2), yet failed to preserve higher-order feature co-dependencies such as MHC gene cluster-mutational burden architecture (Figure 3) and wound-healing-immune activation programs (Figure 7). By contrast, Gaussian Copula and Avatar maintained these complex biological structures and successfully rediscovered clinically relevant biomarkers, including PBRM1-associated angiogenesis in ccRCC and immunoproteasome enrichment in NSCLC (Figure 4).

These results reinforce the necessity of task-specific downstream evaluation: high statistical fidelity does not guarantee preservation of biologically meaningful relationships.

A utility-privacy trade-off was equally apparent. More stringent privacy guarantees tend to reduce the utility of synthetic data, eroding the statistical and biological signals relevant for downstream research. Conversely, synthetic datasets that closely resemble the original may retain residual re-identification risk, particularly in small clinical cohorts where rare feature combinations can be informative. Methods that closely mirror marginal distributions, such as Synthpop, incurred elevated singling-out risks, whereas Gaussian Copula and CTGAN offered comparatively stronger privacy protection (Table 4; Figures 9-10). We also observed consistent attenuation of statistical significance across synthetic datasets. Although the directionality of gene-level, pathway-level, survival, and immune signals was generally preserved, effect sizes and statistical strength were more conservative than in the original data (Figures 4, 7, 8). This suggests that synthetic data are well-suited for hypothesis generation but may underestimate subtle biological effects.

Multi-seed synthesis is essential when conducting downstream biological analyses on synthetic data, as results can vary substantially across replicates and single-run assessments may yield misleading conclusions. Given the computational burden of multi-seed synthesis, robustness was assessed across five independent replicates per method. For statistical comparison, we adopted a Bayesian framework rather than relying solely on null hypothesis significance testing (NHST). With only five replicates, traditional *P* value-based testing can become overly rigid, often leading to dichotomous interpretation when the null hypothesis cannot be rejected. The Bayesian approach estimates the probability that one method outperforms another across the replicate distribution, enabling uncertainty-aware comparison appropriate for small-sample settings.

Regarding privacy evaluation, no consensus currently exists on standard metrics or acceptable risk thresholds for synthetic data. Similarity-based measures are insufficient on two grounds: they do not reliably reflect inference risk, and reporting them may itself facilitate reconstruction by adversaries. Moreover, overlap between similarity and utility metrics complicates the interpretation of the utility-privacy relationship. Accordingly, we employed attack-based evaluations via the Anonymeter framework, directly quantifying singling-out, linkability, and attribute inference risks in alignment with the European Data Protection Board (EDPB) principles. This attack-based approach provides operationally meaningful privacy estimates rather than proxy similarity scores.

Several limitations should be acknowledged. The framework, while multi-dimensional, does not exhaustively cover all possible utility and privacy metrics, particularly for highly task-specific applications, nor all available SDG paradigms. The limited size of the cancer cohorts, consistent with current Phase I and II clinical drug trials, may have constrained generative model performance and reduced the precision of privacy risk estimation. Without large external control datasets, distinguishing population-level signal from unintended memorization remains challenging; privacy risks should therefore be interpreted as relative, scenario-dependent indicators rather than absolute guarantees. Computational constraints limited hyperparameter exploration, and alternative configurations may alter relative performance. Finally, rank-based meta-scores, while enabling integrative comparison, compress absolute performance and should be interpreted alongside absolute metrics and biological validation results.

In summary, rigorous evaluation of synthetic clinical transcriptomic data requires a multi-dimensional framework that jointly assesses statistical fidelity, biological validity, and privacy risk. We encourage researchers to apply this framework to their own datasets, to prioritize preservation of biologically meaningful dependencies over metric-based similarity alone, and to perform task-specific validation when selecting SDG methods. Continued development of biologically grounded metrics and principled privacy assessment strategies will strengthen future benchmarking efforts and support transparent, context-aware adoption of synthetic data in precision oncology.

## 4 Methods

### Benchmark datasets

To ensure the general applicability of the benchmarking framework, we selected paired clinical and transcriptomic data from three independent clinical trials: Braun et al.,^3^ Liu et al.,^19^ and Ravi et al.^20^ These three cohorts address three different cancer types: clear cell renal cell carcinoma (ccRCC), melanoma, and non-small cell lung cancer (NSCLC). Together, these data sets represent a variety of clinical settings and transcriptomic signatures.

Clinical features encompass demographic information (age, sex), clinical outcomes (overall survival, progression-free survival, best objective response), treatment-related variables (treatment arm, prior lines of therapy), and tumor characteristics (performance status, metastasis sites). The ccRCC and melanoma datasets additionally include genomic features such as somatic mutation status (e.g., PBRM1, VHL, TP53 for ccRCC; tumor mutational burden, neoantigen load, and mutational signatures for melanoma) and chromosomal alterations. Molecular features consist of genome-wide bulk RNA-seq gene expression profiles quantified as transcripts per kilobase million (TPM), comprising 40,934, 18,760, and 21,969 genes for ccRCC, melanoma, and NSCLC, respectively.

The ccRCC cohort includes patients treated with nivolumab (anti-PD-1) or everolimus (mTOR inhibitor). The melanoma cohort comprises patients treated with pembrolizumab or nivolumab (anti-PD-1 monotherapy), stratified by prior ipilimumab (anti-CTLA-4) exposure. The NSCLC cohort includes patients treated predominantly with anti-PD-(L)1 monotherapy (81%), with a minority receiving combination therapy with anti-CTLA-4 (17%) or chemotherapy (2%).

### Data processing pipeline

To minimize the confounding impact of data quality on generative model effectiveness, a standardized and reproducible preprocessing pipeline was designed, implemented, and consistently applied across all three datasets. This harmonized processing ensures unbiased comparability among the evaluated synthetic data generation methods. The pipeline consists of five sequential stages: (1) raw data filtering to remove duplicates, unexpressed genes, near-zero variance features, and samples or features exceeding missingness thresholds; (2) transcriptomic harmonization mapping Ensembl IDs to HUGO gene symbols via MyGeneInfo Python library;^31^ (3) feature type classification into categorical, ordinal, and numerical variables; (4) multivariate imputation using MICE,^32^ with missingness indicator flags preserved to retain missingness patterns in synthetic data; and (5) integration of clinical and transcriptomic data via patient-specific identifiers into a unified dataset for downstream synthesis. A schematic overview and full details of each stage are provided in the Supplementary Methods and Figure S1.

### Synthetic data generation protocol

The benchmarking standard was used to assess five methods for the generation of synthetic data: CTGAN,^21^ TVAE,^21^ Gaussian Copula,^16^ Synthpop,^22^ and Avatar^12^ across the three cancer types. The core algorithms in Avatar method uses dimensionality reduction and k-nearest neighbors as its main algorithms.The key settings are the number of nearest neighbors and the number of components. We ran Avatar with k set to 5 and 10, without changing the number of components. These configurations are called Avatar K5 and Avatar K10. These two values were selected to represent a low and a moderate neighborhood size, reflecting the privacy-utility trade-off inherent to k-nearest neighbor anonymization: smaller k values enforce stricter locality constraints and may better preserve fine-grained data structure, whereas larger k values promote greater generalization and typically offer stronger privacy guarantees. The choice of k was informed by a prior benchmarking study applying Avatar to clinical pharmacogenetics data,^33^ and larger values (e.g., *k* = 20) could not be evaluated due to API memory constraints encountered with the high-dimensional datasets used in this study.

We applied all six synthetic data generation methods to each cancer dataset. This process was repeated five times using different random seeds. In total, we created 90 synthetic datasets, with 30 for each cancer type.

Naïve configurations were insufficient for the high-dimensional and heterogeneous nature of this data modality; method-specific adaptations were therefore implemented for each SDG approach. These included feature-type-aware encoding combined with parallelized marginal fitting and vectorized chunking for Gaussian Copula, association-guided feature partitioning into memory-feasible blocks for Avatar, and a graph-centrality-based predictor matrix for Synthpop to constrain cumulative predictor expansion during sequential synthesis. Full implementation details are provided in the Supplementary Methods.

### Statistical fidelity

We measured the statistical fidelity of synthetic data by looking at two main statistical properties: how similar the distributions of individual features are, and how well the relationships between pairs of features are preserved.

#### Univariate similarity

- For each numerical feature *k*, the statistical fidelity between the original and synthetic datasets was quantified using a Kolmogorov-Smirnov(KS)-based similarity score, defined as: 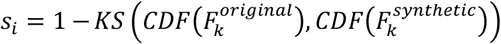, Here, *CDF*() denotes the empirical cumulative distribution function. Scores close to 1 indicate high similarity, while values near 0 indicate low agreement between distributions.
- For each categorical feature *k*, the distributional similarity between the original and synthetic datasets was calculated using a Total Variation Distance (TVD) similarity score defined as: *s*_*j*_ = 1 − *TVD*(*F*_*k,original*_, *F*_*k,synthetic*_). Here, *TVD*() measures the dissimilarity between the empirical distributions of the original and synthetic data. Scores close to 1 indicate high similarity, while values near 0 reflect substantial divergence between distributions.
- These scores were calculated by using the Synthetic Data Vault (SDV) library.^34^

Then, to have a balanced evaluation between the two data modalities (clinical and transcriptomic), the final univariate score for Method A is the average of the mean score of each modality:

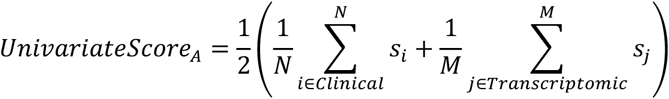

Where *N* and *M* denote the total number of clinical features and genes, respectively. *s*_*j*_ and *s*_*i*_ are the similarity scores calculated via KS or TVD, depending on the nature of the specific feature.

#### Bivariate similarity

To evaluate how well the synthetic data preserves relationships between features, we calculated a bivariate similarity score (*s*_*pair*_) based on the feature types:

- For numerical-numerical pairs: we used Spearman’s rank correlation (*ρ*). The similarity score for a pair (*i, j*) is:

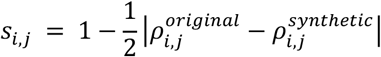
- For categorical-categorical and mixed (numerical-categorical) pairs: Cramér’s V was used. For mixed pairs, numerical features were discretized into ten bins. The similarity score for a pair (*i,j*) is:

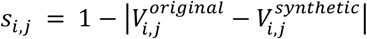

To get the overall bivariate similarity score of Method A, we calculated the mean score for each of the three functional modalities and then averaged them to ensure equal weighting:

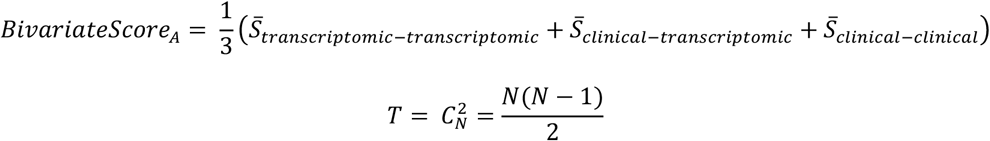

Where:

- 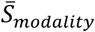 is the average similarity score of all pairs within that category:
  - Clinical-Clinical: 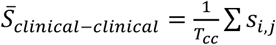 Where 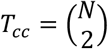
  - Transcriptomic-Transcriptomic: 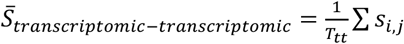 Where 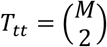
  - Clinical-Transcriptomic: 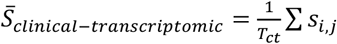 Where *T*_*ct*_ = *N* × *M*
- *N* and *M* denote the total number of clinical features and genes, respectively

### Biological utility

In this evaluation dimension, we assessed whether synthetic transcriptomic data preserved important clinical and biological signals by comparing results from several downstream analyses (DGE, GSEA, ssGSEA, cell type deconvolution, survival analysis, and machine learning-based modeling) on both the original and synthetic datasets. For each task, analyses were run independently on each dataset and results were compared for concordance.

Clinical subgroups were defined for each cohort following the original study definitions. In ccRCC, patients with objective response (complete or partial response) or stable disease with tumor shrinkage and PFS ≥ 6 months were classified as clinical benefit (CB), while those with progressive disease and PFS < 3 months were classified as non-clinical benefit (NCB). Only nivolumab-treated patients were included in the ccRCC analysis. In melanoma and NSCLC, responders were defined as patients achieving complete or partial response (CR/PR) per RECIST v1.1; progressors in melanoma were limited to progressive disease (PD), whereas non-responders in NSCLC additionally included stable disease (SD). Additional subgroups from the original publications, such as genomic subtypes and prior therapy groups, were also incorporated where relevant. The following sections describe the analytical procedures for each task.

#### Differential gene expression (DGE)

To evaluate synthetic data in the DGE task, we introduced the Gene Concordance Score (GCS). This metric quantifies the agreement in gene or pathway significance and directionality between original and synthetic datasets. We calculated GCS in three steps:

- Step 1: Rank Score Calculation: each gene *i* is assigned a rank score (*RS*) that combines its effect size and statistical significance:

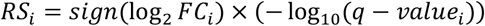 The scatter plots comparing gene RS between the original data and the synthetic datasets generated by each SDG method across all three cancer cohorts are provided in Figures S5, S6, and S7 in the Supplementary Data.
- Step 2: Concordance Zone Identification: A significance boundary is defined as:

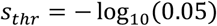 Using this threshold, four concordance zones are defined in the *RS*_*original*_ versus *RS*_*synthetic*_ plane:
  - Significant Zones: Genes appearing in the same extreme quadrants (both significantly up-regulated or both significantly down-regulated in both datasets). Let *N*_*sig*_ be the count of genes in these zones.
  - Non-significant Zones: Genes that remain non-significant and maintain the same direction of *Log*_2_*FC* in both datasets. Let *N*_*non-sig*_ be the count of genes in these zones.
- Step 3: GCS Calculation: The score is calculated by this formula:

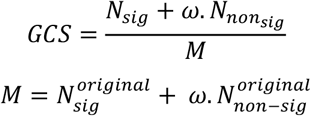

Where 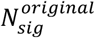 and 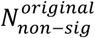 are computed from the full set of genes in the original dataset.

We set a weight *ω* = 0.5 to assign a lower weight to non-significant genes, thereby prioritizing preservation of biologically meaningful differential signals.

#### Gene set enrichment analysis (GSEA)

We implemented GSEA with the GSEAPy Python package. Pre-ranked GSEA was performed with 10,000 gene set permutations to calculate enrichment *P* values, FDR *Q* values and NES for each pathway. We analyzed all three cancer cohorts using the Hallmark Gene Sets from the Molecular Signatures Database (MSigDB).

To measure how well the original and synthetic data matched, we employed the Pathway Concordance Score (PCS). Analogous to GCS, PCS evaluates the agreement in enrichment significance and directionality at the pathway level. For each pathway *p*, a rank score (*RS*_*p*_) was defined as:

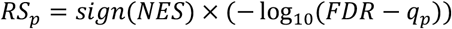

Concordance was subsequently assessed within the same signed significance framework and weighting scheme described for GCS.

#### Single sample gene set enrichment analysis (ssGSEA)

We performed ssGSEA using GSEAPy with the log-rank method and the Hallmark gene sets from MSigDB. To assess preservation of pathway-level activation signals at the single-sample level, we quantified the similarity between the distributions of NES in the original and synthetic datasets.

For each pathway *p*, similarity was measured using the KS-complement score:

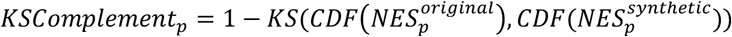

The overall performance for a given method was defined as the mean KS complement score across all 50 Hallmark pathways.

#### Predictive modeling

To assess if the synthetic data preserved the predictive utility of the original data, we compared the performance of classification models on synthetic datasets against those trained on the original dataset. The model was used to distinguish progressors from non-progressors within the Ipilimumab-treated subgroup of the melanoma cohort. Three classifiers, including logistic regression (applied in the original study by Liu et al), random forest, and multilayer perceptron, were evaluated using repeated stratified cross-validation (5 folds, 3 repeats, *RepeatedStratifiedKFold* function). The computations were implemented using the scikit-learn Python library.^35^

For each cross-validation split, models were trained either on the original training set or on the corresponding synthetic training set, while performance was consistently evaluated on the same held-out test folds from the original data. This design ensured that all comparisons were conducted on identical real-data test distributions. Model performance was quantified using the ROC-AUC metric.

To statistically compare predictive performance between models trained on original versus synthetic data, Wilcoxon signed-rank tests were applied across cross-validation folds. A non-significant difference combined with comparable or improved mean performance was interpreted as evidence that the synthetic data preserved predictive utility.

#### Immune cell deconvolution

We used the CIBERSORTx algorithm for cell deconvolution to match bulk transcriptomic data. The LM22 signature estimated the fractions of 22 immune cell types. We ran the analyses in absolute mode with B-mode batch correction, turned off quantile normalization, and performed 1,000 permutations. We excluded samples with CIBERSORTx deconvolution *P* > 0.05. This procedure was consistently applied across all three cancer cohorts.

To quantify preservation of inferred immune cell compositions, we measured the similarity using the Aitchison distance, which is appropriate for compositional data.^28^

Let *x*_*ij*_ denote the estimated fraction of cell type *j* in the sample *i*. After constraining to the top *k* (*k* = 10) most abundant cell types (identified from the original data), zero values were handled using multiplicative replacement.

The compositional center (geometric mean) for each cell type *j* was computed as:

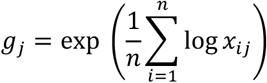

The centered log-ratio (CLR) transformation of the compositional center vector *g*_*j*_ is:

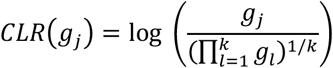

The Aitchison distance between original and synthetic datasets is then defined as the Euclidean distance in CLR space:

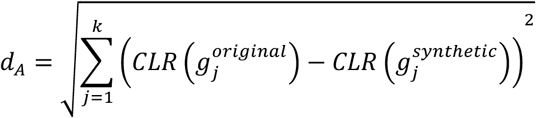

To ease of comparison, the distance was converted into the Aitchison distance score:

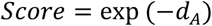

where a higher value indicates the better compositional concordance.

Next, we did differential analysis on the immune cell proportion matrices. For ccRCC, we used the original study design and CD8 immunofluorescence assays to classify tumors as immune-infiltrated, immune-desert, or immune-excluded, and compared these groups directly. For melanoma and NSCLC, we compared responders and non-responders or progressors. Regarding single-cell signatures analysis in the Melanoma and NSCLC cohort, we utilized the corresponding signature following the original studies.^19,20,29,36,37^

#### Survival analysis

To quantify the survival similarity between the original and synthetic datasets, the C-index was first estimated independently for each dataset using a Cox proportional hazards model from either OS or PFS, as implemented in the *lifeline* Python package. Then, the final C-index score was defined as:

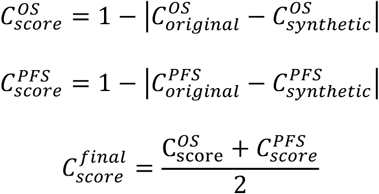

Where 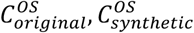 denote the C-index of the original and synthetic datasets based on OS; 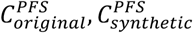 denote the C-index of the original and synthetic datasets based on PFS.

In terms of Kaplan-Meier curve analysis, *P* value of a log-rank test was calculated to compare survival curves of two observed groups.

### Privacy evaluation

According to the European Data Protection Board (EDPB), there are three criteria to assess the anonymization methods.^27^ They are singling-out, linkablity and inference:

- Singling out is the ability to isolate a specific individual within a dataset.
- Linkability is the ability to connect two records belonged to the same patient.
- Inference is the ability to deduce, with significant probability, the value of an individual using the anonymized dataset.

In our benchmarking study, we performed attack-based evaluations via the Annonymeter framework.^18^ This framework enables us to simulate attack scenarios and evaluate directly singling out, linkablity and inference risk, which is coherent and aligned with EDPB. We designed a privacy assessment adapted to our data modality, which is described clearly below.

#### Singling out risk

Anonymeter evaluates singling out risk by attempting to isolate individual records within the original dataset using unique predicates derived from patterns in the synthetic data. There are two algorithms: univariate PredicateFromAttribute and the multivariate MultivariatePredicate, to generate singling-out predicates. The univariate algorithm generates predicates from single features, while the multivariate configuration leverages attribute combinations to identify individuals within the dataset.

To evaluate univariate singling-out attacks, we tested different configurations by randomly sampling 25%, 50%, 75%, and 100% of the dataset’s attributes. For each of these configurations, Anonymeter generated 10,000 predicates (as defined by the *n*_*attacks*_ parameter) to simulate adversary attempts.

For multivariate singling-out attacks, attackers were provided with combinations of multiple attributes, with the number of attributes per attack set to {2,3,5,7,10,20,50}. Risk estimation followed the same randomized attack protocol as in the univariate setting where 10,000 predicates were generated.

#### Linkability

Anonymeter evaluates linkability risk by determining if an attacker can use a synthetic dataset to connect *disjoint* sets of attributes, referred to as auxiliary information, belonging to the same individual. The framework simulates this by identifying the a nearest neighbor in the synthetic data for each attribute set independently, utilizing the Gower coefficient to measure similarity across both categorical and numerical data. That means the input for Anonymeter in linkability assessment is the 2 disjoint sets of features. In our experiment, we want to validate the risk that an attacker can leverage synthetic datasets to connect molecular profile to clinical information of a patient. Therefore, 2 disjoint sets are clinical features and transcriptomic features. We iteratively picked up 25%, 50%, 75% and 100% genes in gene expression matrix to link all clinical variables.

#### Attribute inference risk

This risk was quantified by assessing an adversary’s ability to infer sensitive attributes (secrets) from auxiliary gene expression features. Similar to linkability, Anonymeter utilized the k-nearest neighbor algorithm to predict secret attributes in the original data based on synthetic samples. For inference attacks, each clinical variable was treated as a potential secret, and risks were computed independently for each target attribute.

#### Overall privacy score

In the Anonymeter framework, besides the main attack risk using synthetic data (*R*_*attack*_), the naïve risk (*R*_*naive*_), corresponding to a baseline obtained under random guessing, was also reported. According to the authors, if the main attack risk is lower than or equal to the random naïve baseline (*R*_*attack*_ ≤ *R*_*naive*_), the attack is classified as “failed” and should not be reported as “zero risk”. Therefore, we reported failed attacks following their suggestion (Figure 9). In the overall privacy score calculation, for each attack configuration *c*, we used naïve risk (*R*_*naive*_) instead of using 0 of the attack failed, as

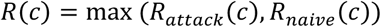

For each privacy category *m*, the final risk for a given synthetic data generation method was obtained by averaging over all configurations:

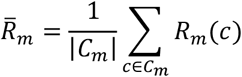

Where *C*_*m*_ denotes the set of attack configuration for category *m*.

To obtain an overall privacy risk, we aggregated the four privacy categories, including univariate singling-out, multivariate singling-out, linkability, and attribute inference, using an unweighted mean:

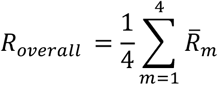

Finally, we report an overall privacy score defined as

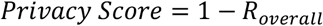

where higher values indicate stronger privacy preservation. All privacy evaluations were repeated across five random seeds to account for stochastic variability in synthetic data generation.

### Rank-derived meta-score

We adapted the ranking framework of Yan et al.^30^ to integrate 8 evaluation dimensions: univariate similarity, bivariate similarity, DGE, GSEA, ssGSEA, dell deconvolution, survival analysis, and privacy.

In the context of each cancer dataset, 6 SDG methods (5 replicates each; 30 candidates) were ranked per dimension. The rank-score of the method *m* under the dimension *a* was defined as the average rank across its five replicates:

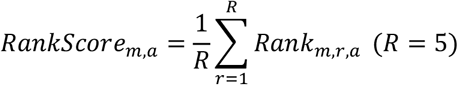

Evaluation dimensions were grouped into:

- Statistical fidelity: univariate, bivariate similarity
- Biological utility: DGE, GSEA, ssGSEA, cell deconvolution, survival analysis
- Privacy

The final meta-score within one cancer dataset was:

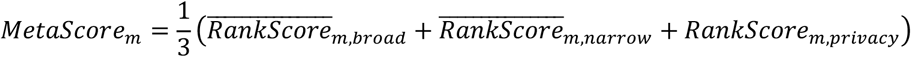

Where each dimension contributes equally.

As for cross-cancer comparison (three cancer types), we summed the dimension-level rank-derived scores across cancers:

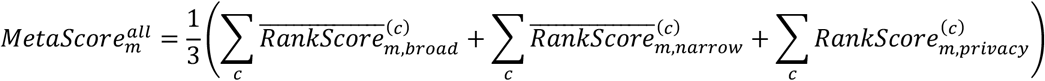

### Statistical tests and Bayesian comparison

All statistical tests were reported alongside the corresponding results in the main text. We used *scipy*.*stats* Python package to perform hypothesis testing.^38^ To compare numerical features between patient categories (e.g., DGE, differential analysis of ssGSEA score, cell type signatures), two-sided non-parametric Wilcoxon rank-sum test (*ranksums* function) or Mann-Whitney U test (*mannwhitneyu* function) was applied, depending on the test used in the original studies. In machine learning model comparison, the Wilcoxon signed-rank test was employed (*wilcoxon* function). Multiple hypothesis testing was controlled using the Benjamini-Hochberg procedure for FDR correction (*multipletests* function from the *statsmodels* Python package).^39^

We performed pairwise Bayesian comparisons following the framework proposed by Benavoli et al., using the *baycomp* Python library.^40^ For each cancer cohort, all SDG methods were compared pairwise based on their performance scores across five independent replicates. A Bayesian correlated t-test was applied to estimate posterior probabilities for three mutually exclusive hypotheses:

- Better probability (*P*(*SDG*_1_ > *SDG*_2_)): the probability that *SDG*_1_ outperforms *SDG*_2_.
- Worse probability (*P*(*SDG*_1_ < *SDG*_2_)): the probability that *SDG*_1_ underperforms *SDG*_2_.
- Practical equivalent probability: the probability that the performance difference lies within a predefined Region of Practical Equivalence (ROPE).

The ROPE threshold was set to 0.01, below which performance differences were considered negligible. The resulting “better” probabilities were visualized as *N* × *N* heatmaps, where each cell represents *P*(*row* > *column*), defined as the posterior probability that the method in the row outperforms the method in the column for the corresponding cohort.

## Data and code availability

The synthetic datasets of three clinical cohorts, their replicates, analysis code, and notebooks generating figures are deposited at https://doi.org/10.5281/zenodo.19742953. SynOmicsBench source code (Python) is provided in our GitHub repository at https://github.com/trinhthechuong/SynOmicsBench. Detailed documentation of SynOmicsBench is available at https://trinhthechuong.github.io/SynOmicsBench/.

## Acknowledge

The authors appreciate the valuable suggestions from Dr. Antoine Boutet about the Anonymeter framework. We also appreciate the enthusiastic support from Morgan Guillaudeux and the Octopize technical team for adapting Avatar with genomic high-dimensional data. The computations in this study were carried out using the GRICAD infrastructure (https://gricad.univ-grenoble-alpes.fr), which is supported by Grenoble research communities. This work was supported by the DIGPHAT project (Multi-scale and longitudinal data modeling in pharmacology: toward digital pharmacological twins), which has received funding from the French research initiative “France 2030” through the program PEPR Digital Health under ANR grant agreement no. 22-PESN-0017, by the KATY project, which has received funding from the European Union’s Horizon 2020 research and innovation program under grant agreement no. 101017453, and by the CANVAS project, which has received funding from the Horizon Europe twinning program under grant agreement no. 101079510.

## Author contributions

T-C.T., JB.W., G.U., and C.B. conceived the study. T-C.T. developed the code and performed the analysis. T-C.T., JB.W., G.U., and C.B. wrote the manuscript.

