## Supplementary Data for "A unified benchmark of synthetic data generation for clinical transcriptomic cancer cohorts"

### A. Supplementary Methods

#### 1. Standardized preprocessing pipeline

To control the confounding impact of data quality and comparison without biases, we designed a standardized data preprocessing pipeline applied to the original data before training the generative models. The pipeline consists of five sequential stages:

- **Raw Data Filtering:** Quality control was initially conducted on the molecular and clinical data. Duplicate entries and entries without unique identifiers were removed, leaving a single primary record. For transcriptomics data, genes that were not expressed (zero counts) across all samples or exhibited near-zero variance were excluded. Additionally, a threshold on the degree of missingness was employed to remove features or patients with too much missing data, to ensure enough data density for subsequent analyses.
- **Transcriptomic Harmonization:** The MyGeneInfo Python library was used to map Ensembl IDs to HUGO gene symbols.[1] For datasets already reporting gene expression in HUGO symbols, this step was omitted. In cases of many-to-one Ensembl-to-gene mapping, expression values for redundant Ensembl IDs were averaged. Additionally, this step generated a dictionary to group genes exhibiting identical expression profiles across the cohorts.
- Feature type classification: A metadata schema was constructed for each input dataset, specifying the data type of every feature. Features were categorized as categorical, ordinal, or numerical. Categorical features included non-numeric variables, binary variables, or numeric variables with fewer than ten unique values, and were automatically inferred by the pipeline (e.g., sex, treatment arm). Ordinal features represented categorical variables with an inherent order but undefined distances between levels, such as disease stage or number of prior therapies. These were typically pre-encoded, but in ambiguous cases, expert clinical annotation was required. All remaining continuous variables, including age, survival metrics, and gene expression, were classified as numerical.
- Multivariate imputations: To improve the quality of imputation, categorical variables were one hot encoded, ordinal variables were encoded using an ordinal encoder and numerical variables were scaled using Min-Max scaler. Missing values were filled in by using Multivariate Imputation by Chained

Equations (MICE).[2] MICE was preferred because it is flexible in handling the heterogeneity of this data modality and can accommodate missing completely at random (MCAR) and missing at random (MAR) mechanisms.[2] When facing non-convergence or multicollinearity, MICE didn't work, so K-nearest neighbors (KNN) was used as an alternative.[3] In order to explicitly maintain patterns of missingness in the synthetic data, binary indicators of missingness were created for those features that had missing values before imputation. These flags indicated whether a value was originally observed or missing, and under the generative framework also allow for learning of feature-specific missingness patterns during training and synthesis of new data. Once the imputation was done, all the features were inverse transformed to the original scale in order to retain the biologically meaningful distributions before the synthetic data is generated.

- Data Integration: The clinical and transcriptomic data were linked through patient-specific identifiers to create an integrated dataset that could be used for subsequent synthetic data modelling and analysis.

**2. Method-specific adaptations for high-dimensional SDG.**

Naïve fitting techniques did not work because the data was high-dimensional and heterogeneous. Instead, we used method-specific procedures to generate synthetic data at scale:

***<u>Gaussian Copula:</u>*** We used one-hot encoding for categorical features. For ordinal categorical features, we used the OrdinalEncoder from Scikit-learn.[18] To manage high-dimensional data, we fit univariate marginal distributions in parallel across multiple CPU cores using joblib. We grouped column-wise operations into batches with a vectorized chunking approach to reduce communication overhead. When inverting, we mapped one-hot encoded variables back to their original categories by picking the category with the highest probability. For ordinal variables, we rounded synthetic values to match the original labels.

***<u>Avatars:</u>*** Avatars is a proprietary tool developed by Octopize that requires a license. We used the official Python client to upload the original data to the API and receive the synthetic data. Because of memory limits with large, high-dimensional datasets, we could not upload the full dataset at once. Based on advice from the Avatars technical team, we split each dataset into blocks of fewer than 4,000 records. We anonymized these blocks one at a time using the API, then combined the synthetic blocks to create a single synthetic dataset. To partition features, we measured pairwise associations. We used Spearman's rank correlation for numerical features and Cramér's V for categorical features. For associations between numerical and categorical features, we divided numerical variables into ten bins and treated them as categorical. We then created a distance matrix from the correlation matrix as follows:

$$distance\ matrix = 1 - \ |correlation\ matrix|$$

A lower distance value means a stronger association between a pair of features. Hierarchical clustering with average linkage and the distance matrix to cluster features. Heatmaps of the clustered distance matrices for two cancer types are shown in the Supplementary.

**Synthpop:** Synthpop generates synthetic data by iterating through a predefined sequence of variables, where each new variable is synthesized using a conditional predictive model trained on the previously generated variables in the sequence. If the default configuration is used, the order aligns with the input data and, each new variable is synthesized using all previously generated features. As the synthesis moves further down the sequence, the number of predictors increases in an arithmetic progression, since each model incorporates all previously synthesized variables as inputs, leading to the memory issue and significant increase in computation time. To miltigate it, we designed the predictor matrix to define the synthesis order, importantly, the number of predictors to model each feature.

Particularly, pairwise associations were computed by the framework applied to Avatars (Spearman for numerical features and Cramér's V for categorical features). The correlation matrix was transformed into a distance matrix $d_{ij} = 1 - |\rho_{ij}|$, where $\rho_{ij}$ denotes the association between feature $i$ and $j$. A binary adjacency matrix was then constructed by thresholding this distance matrix ($d_{ij} \leq 0.7$). To further constrain computational resources, the number of candidate predictors for each feature was capped at 500 by retaining the strongest associations.

The resulting binary matrix was interpreted as an undirected graph, where nodes represent features edges correspond to pairwise associations. To arrange the synthesis order, we ranked features using a two-criteria centrality strategy based on degree centrality (primary criteria) and eigenvector centrality (second criteria in the case of ties). The final predictor matrix was constructed as a lower-triangular structure, allowing each feature to depend only on highly correlated features that appear earlier in the sequence. This ordering encourages influential variables to be generated first, allowing downstream variables to condition on the most informative predictors while preventing cumulative predictor expansion in high-dimensional settings.

### B. Supplementary Figure

#### 1. Standardized preprocessing pipeline

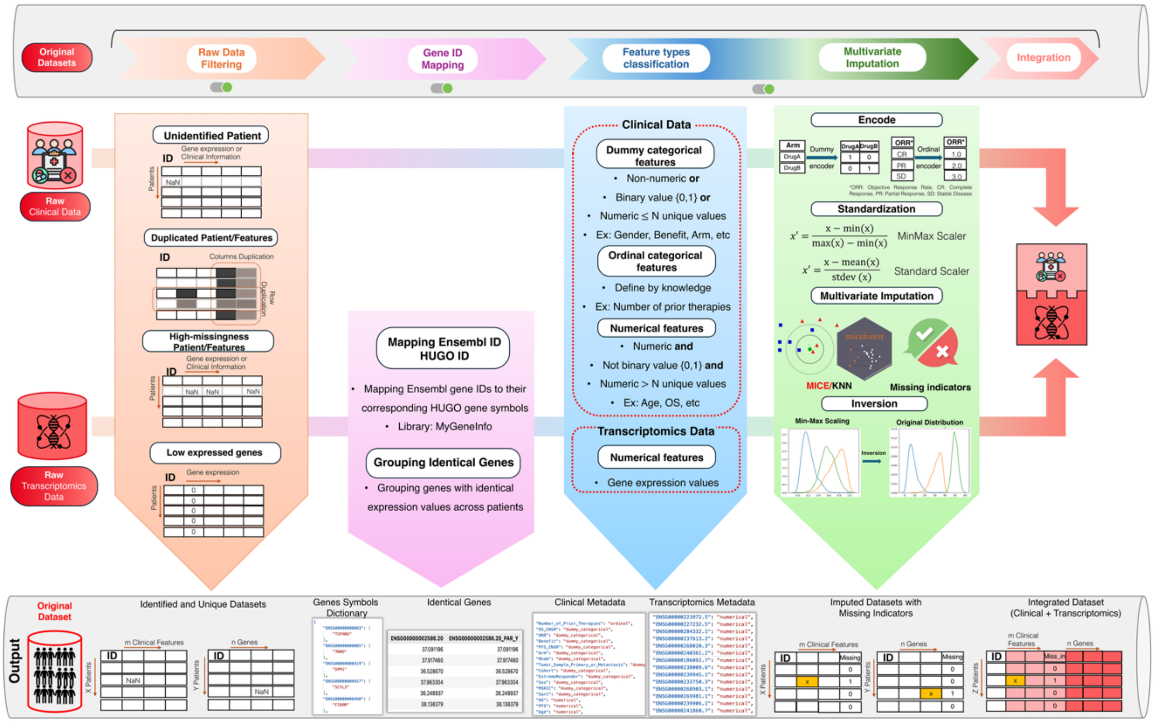

**Figure S1.** A schematic overview of the data processing pipeline.

#### 2. Broad utility

##### 2.1 Univariate similarity

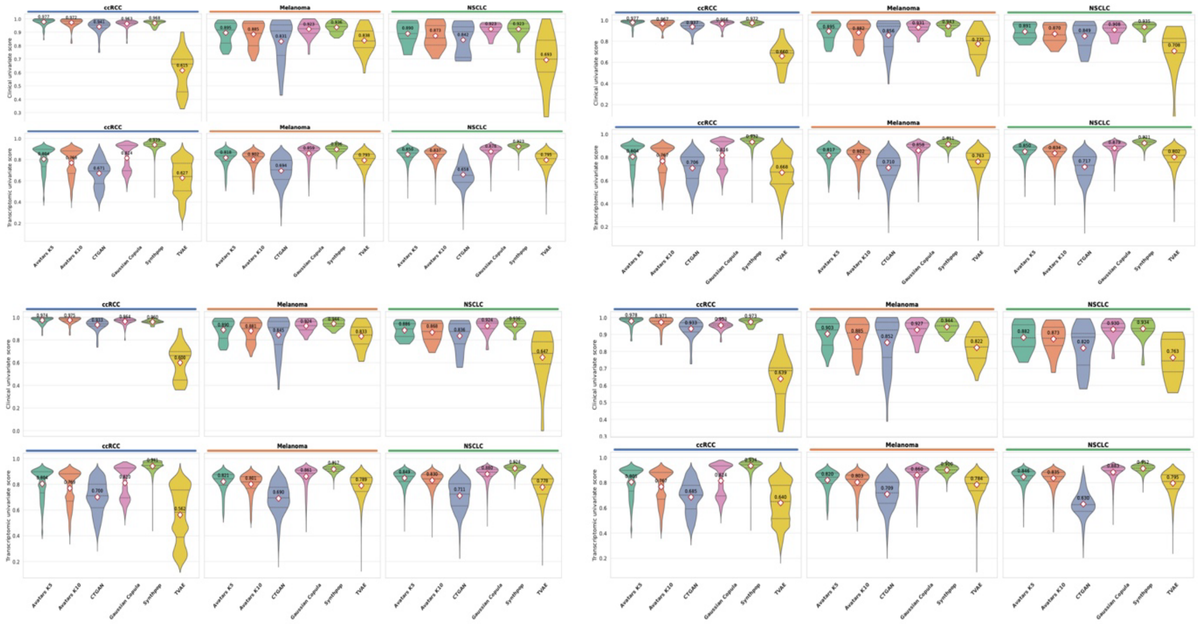

**Figure S2.** Violin plot distribution of univariate similarity scores across replicates. Violin plots display the distribution of univariate similarity scores for clinical variables and transcriptomic features across the ccRCC, melanoma, and NSCLC cohorts. Numerical attributes were evaluated using the KS statistic, whereas categorical attributes were assessed using TVD. Each violin represents the feature-level distribution of similarity scores within a cohort and data modality.

#### 2.2 Bivariate similarity

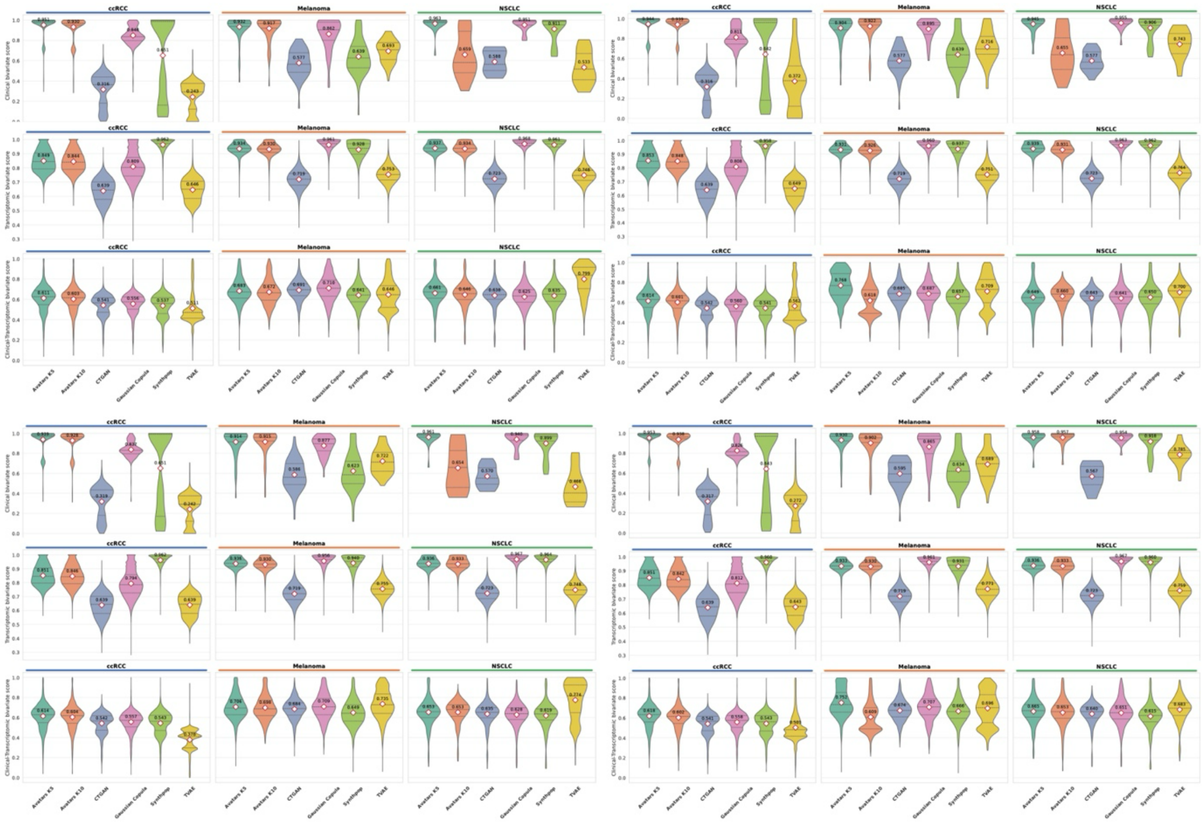

**Figure S3.** Violin plot distribution of pairwise similarity for strongly correlated feature pairs across synthetic replicates. Violin plots depict the distribution of bivariate similarity scores for feature pairs with high correlation in the original data ($|\rho| \geq 0.5$), stratified by clinical-clinical, transcriptomic-transcriptomic, and clinical-transcriptomic associations across the ccRCC, melanoma, and NSCLC cohorts. Each violin represents the distribution of similarity scores across selected feature pairs, with central markers indicating mean values.

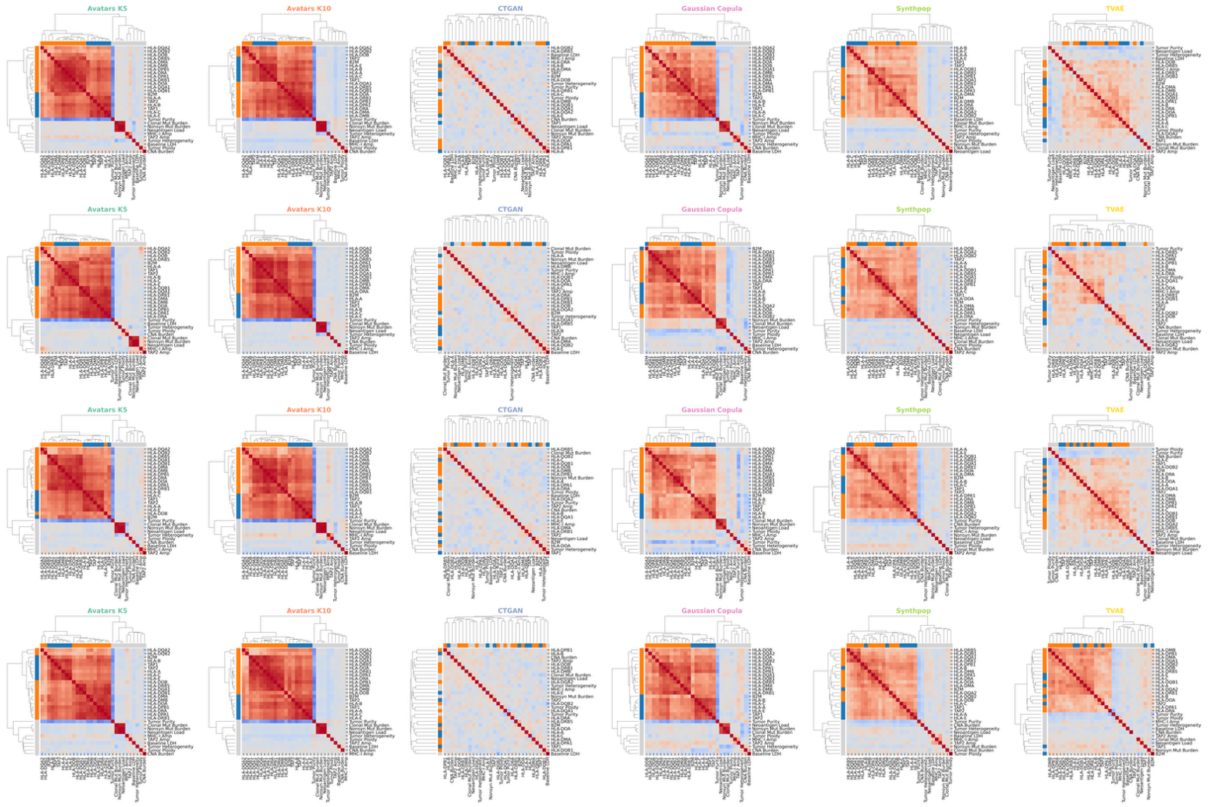

**Figure S4.** Hierarchical clustering of feature correlation matrices in the melanoma cohort across synthetic replicates. Correlation heatmaps with hierarchical clustering depict the structural organization of feature-feature associations in the melanoma dataset.

#### 3. Narrrow utility

##### 3.1 Differential gene expression (DGE)

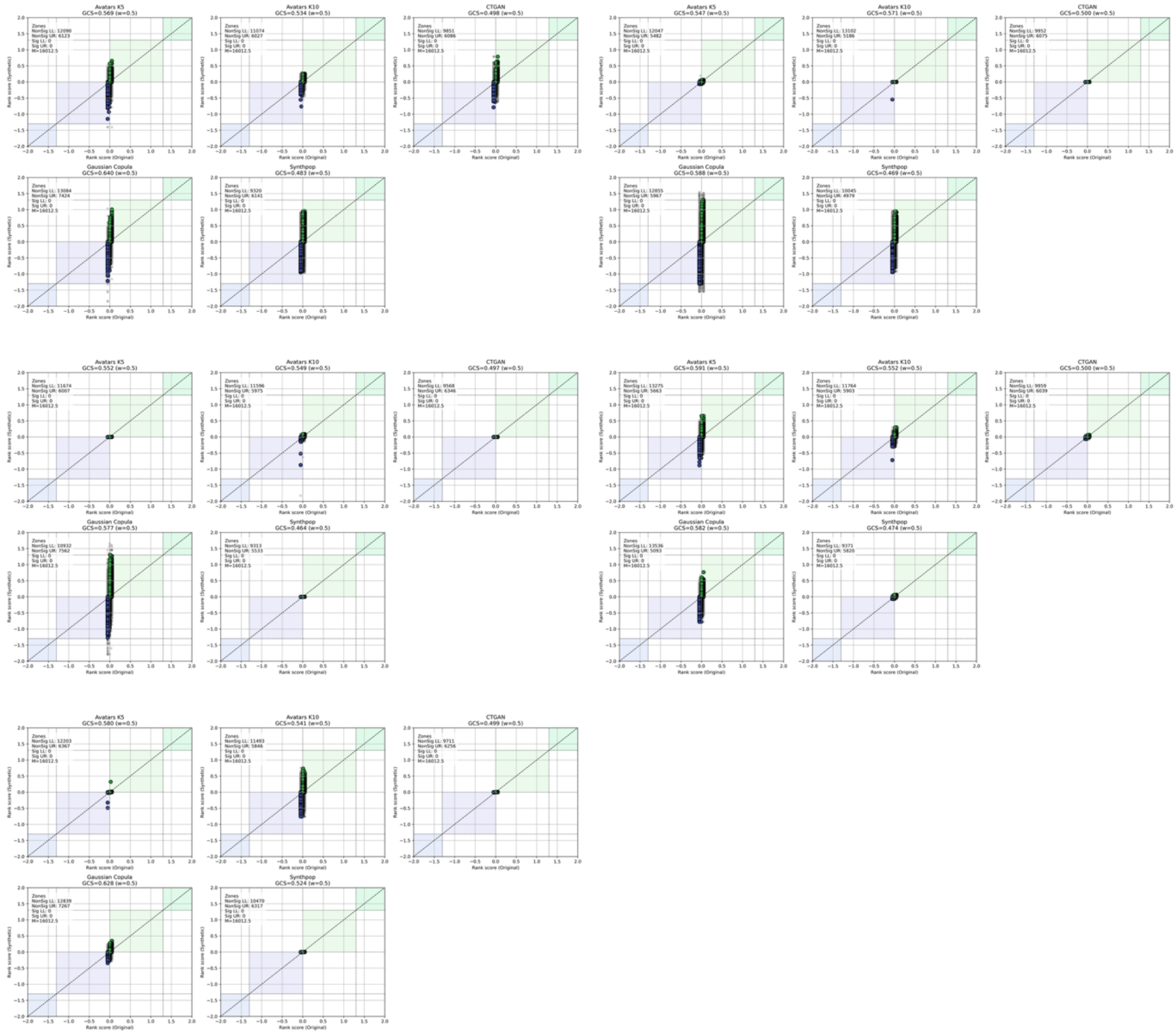

**Figure S5.** GCS scatter plots of ccRCC cohort. These scatter plots compare gene rank score between the original data and synthetic datasets of each SDG method. Rank scores combine the sign of the $log_2(FC)$ and the, $P$ value (Wilcoxon rank sum test). Pathways located in the upper-right (UR, light green) and lower-left (LL, light purple) quadrants indicate concordant regulation direction and the level of significance ($P < 0.05$).

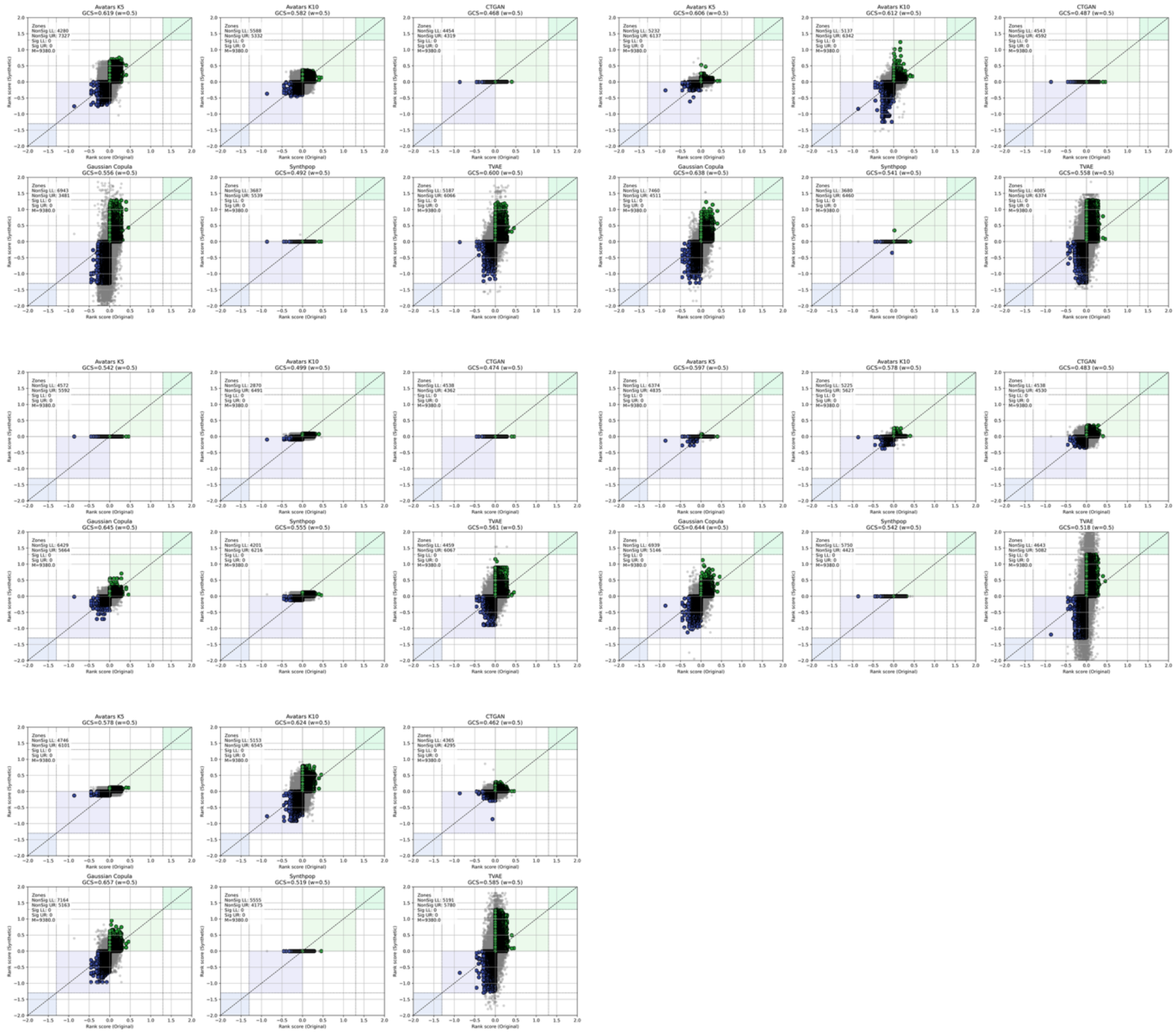

**Figure S6.** GCS scatter plots of Melanoma cohort. These scatter plots compare gene rank score between the original data and synthetic datasets of each SDG method. Rank scores combine the sign of the $log_2(FC)$ and the, $P$ value (Wilcoxon rank sum test). Pathways located in the upper-right (UR, light green) and lower-left (LL, light purple) quadrants indicate concordant regulation direction and the level of significance ($P < 0.05$).

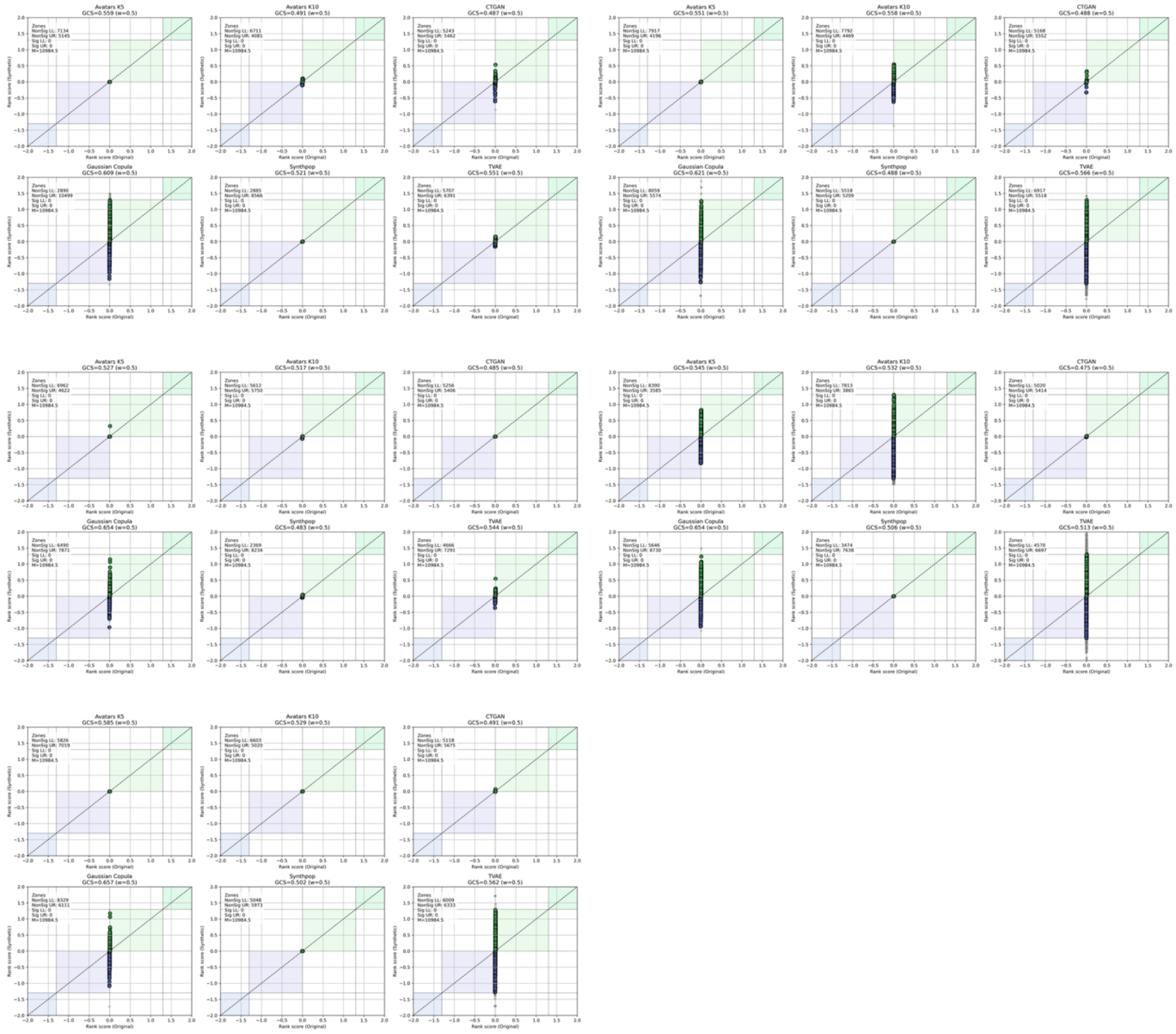

**Figure S7.** GCS scatter plots of NSCLC cohort. These scatter plots compare gene rank score between the original data and synthetic datasets of each SDG method. Rank scores combine the sign of the $log_2(FC)$ and the, $P$ value (Wilcoxon rank sum test). Pathways located in the upper-right (UR, light green) and lower-left (LL, light purple) quadrants indicate concordant regulation direction and the level of significance ($P < 0.05$).

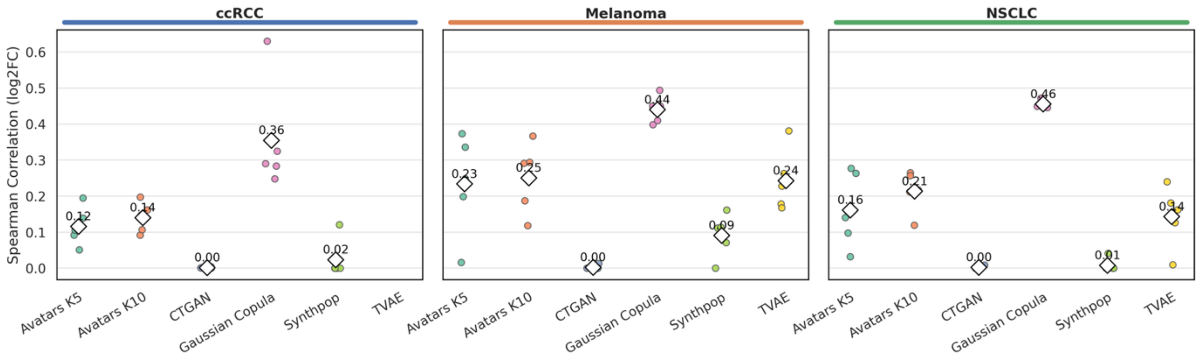

**Figure S8.** Strip plots of Spearman rank correlations of gene-wise $\boldsymbol{log_2(FC)}$ between synthetic datasets and original data across three cancer cohorts.

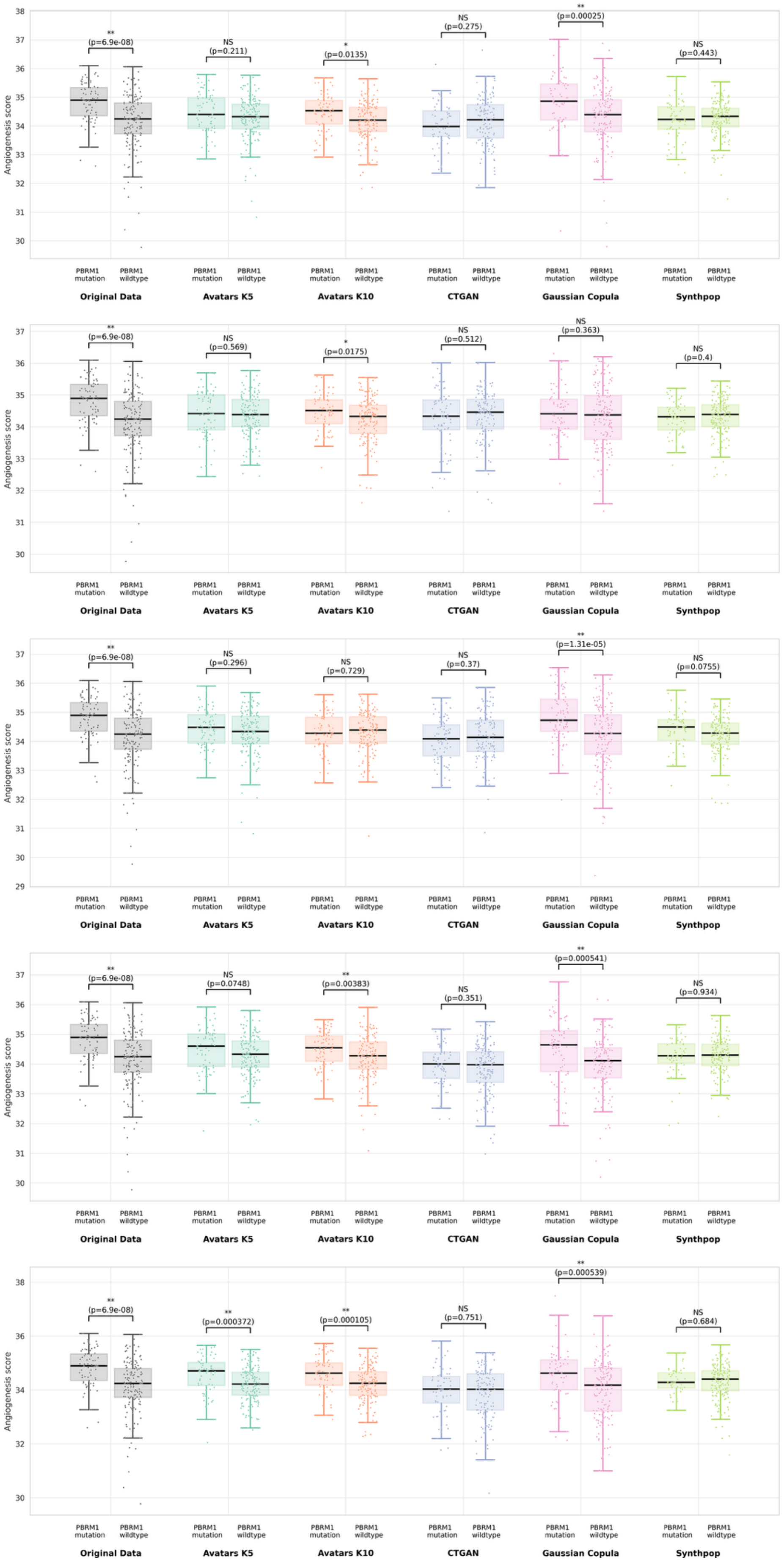
Angiogenesis score
PBRM1 mutation
PBRM1 wildtype
Original Data
Avatars K5
Avatars K10
CTGAN
Gaussian Copula
Synthpop
** (p=6.9e-08)
NS (p=0.211)
* (p=0.0135)
NS (p=0.275)
** (p=0.00025)
NS (p=0.443)
** (p=6.9e-08)
NS (p=0.569)
* (p=0.0175)
NS (p=0.512)
NS (p=0.363)
NS (p=0.4)
** (p=6.9e-08)
NS (p=0.296)
NS (p=0.729)
NS (p=0.37)
** (p=1.31e-05)
NS (p=0.0755)
** (p=6.9e-08)
NS (p=0.0748)
** (p=0.00383)
NS (p=0.351)
** (p=0.000541)
NS (p=0.934)
** (p=6.9e-08)
** (p=0.000372)
** (p=0.000105)
NS (p=0.751)
** (p=0.000539)
NS (p=0.684)

**Figure S9.** PBRM1 mutation-associated angiogenesis signal in ccRCC across synthetic replicates. Angiogenesis scores stratified by PBRM1 mutation status are shown. Statistical significance was assessed using a two-sided Wilcoxon rank-sum test.

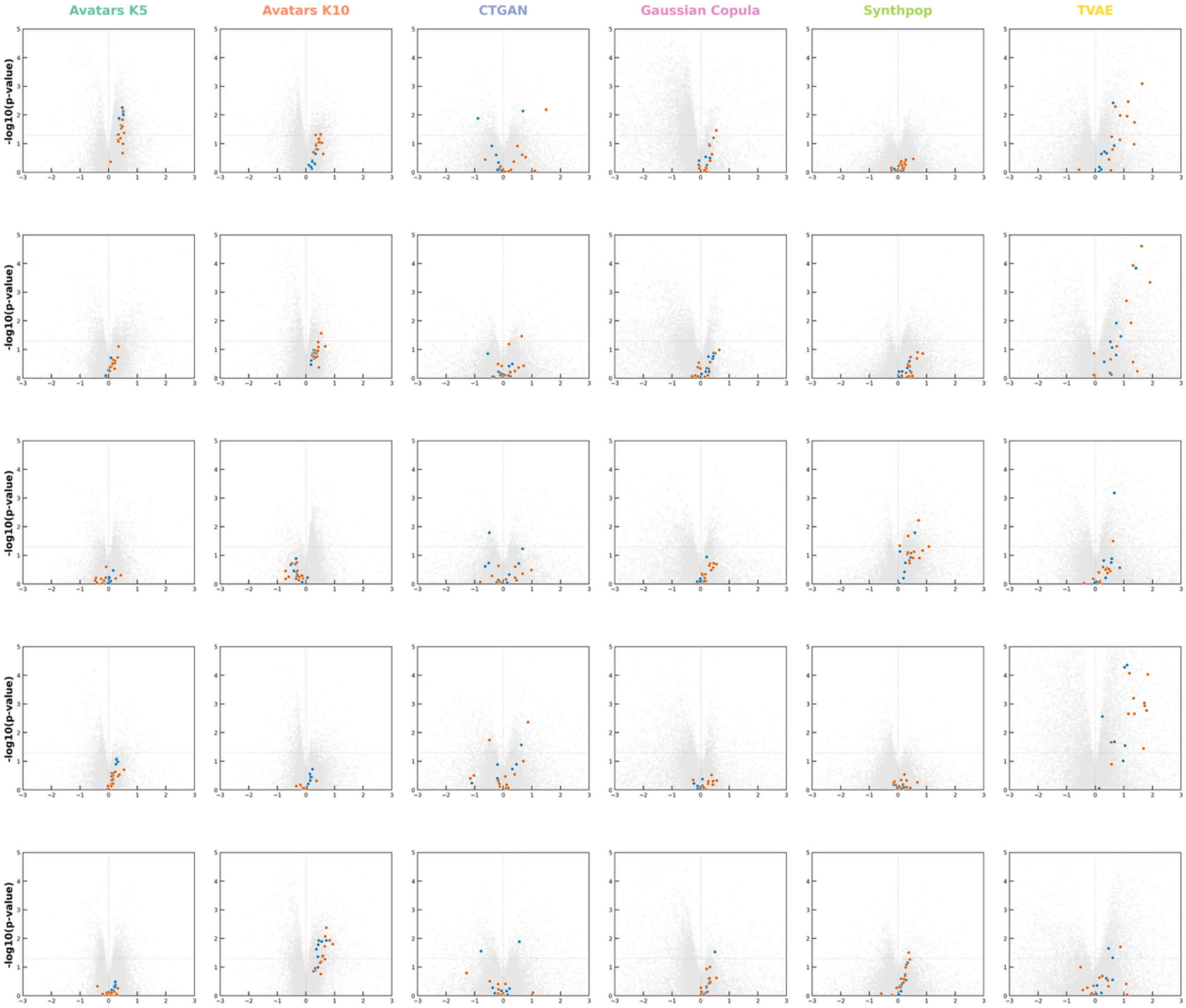

**Figure S10.** Preservation of responder-associated MHC gene signatures in melanoma across synthetic replicates. Volcano plots compare responders and non-responders. Highlighted points are MHC class I and MHC class II HLA genes. Statistical significance was assessed using a two-sided Wilcoxon rank-sum test.

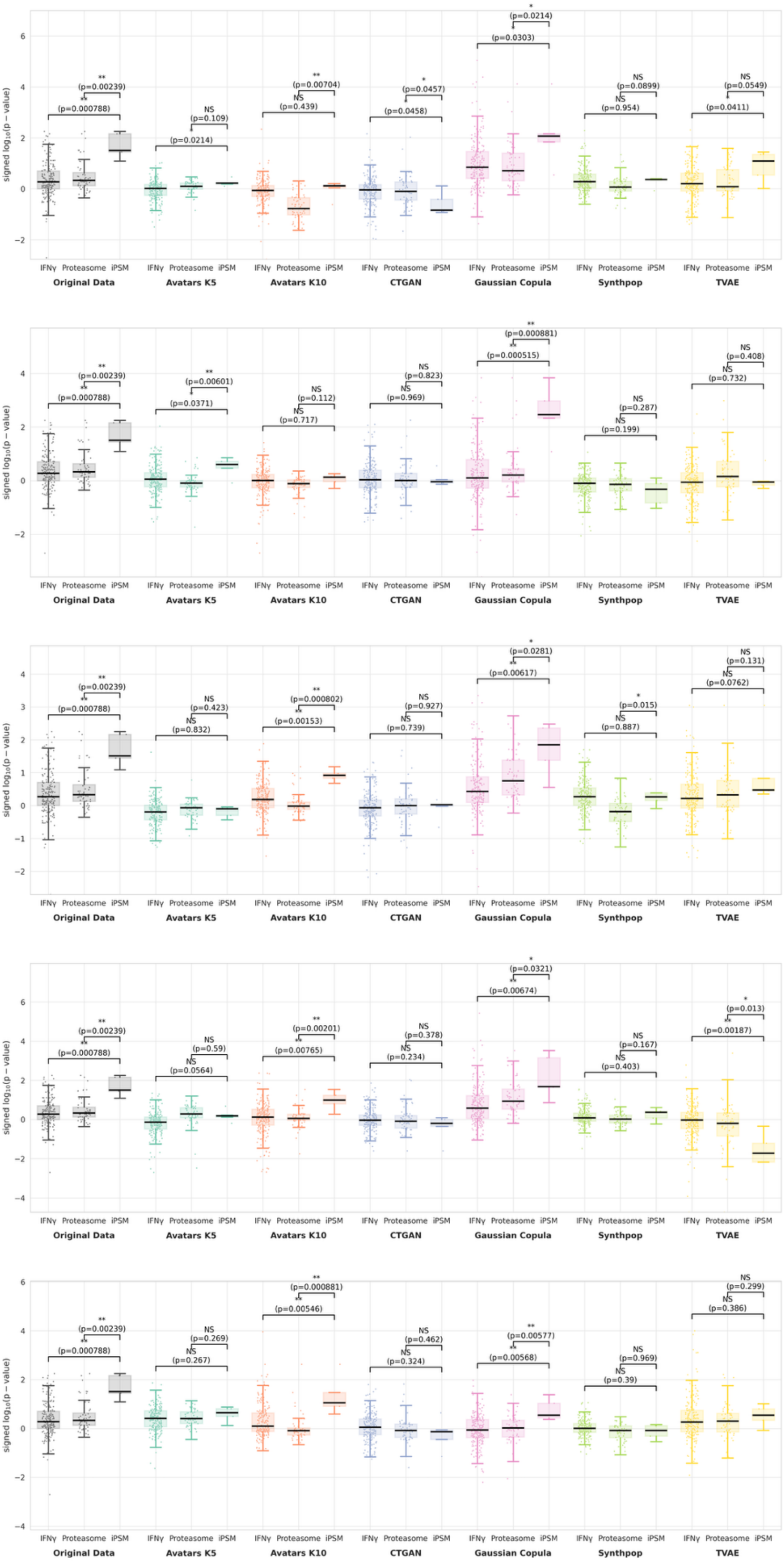

**Figure S11.** Re-discovery of immunoproteasome enrichment in NSCLC across synthetic replicates. Signed $-\log_{10}(p - value)$ distributions compare IFN-$\gamma$ response genes, immunoproteasome (iPSM) components, and canonical proteasome genes between responders and non-responders. Statistical significance was assessed using a two-sided Wilcoxon rank-sum test.

#### 3.2 Gene set enrichment analysis (GSEA)

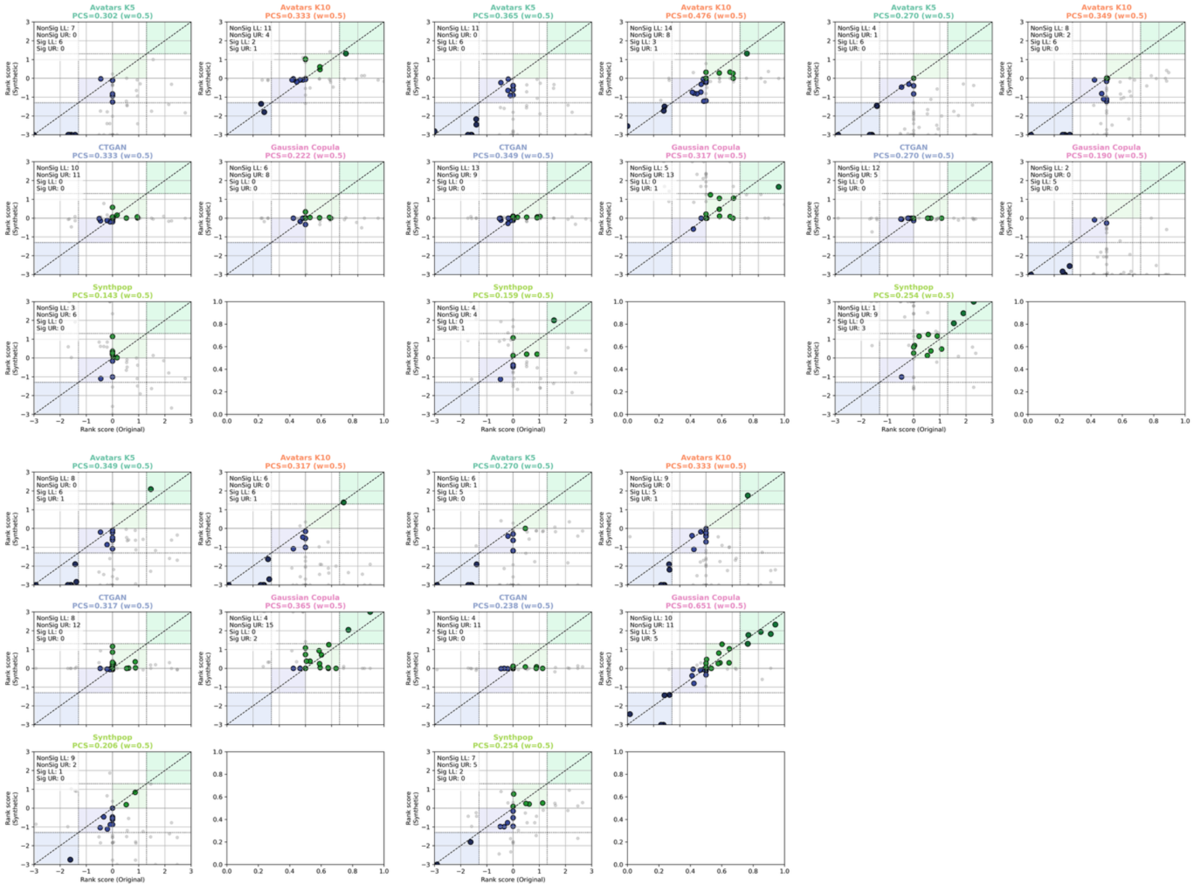

**Figure S12.** PCS scatter plots of ccRCC cohort across synthetic replicates. These scatter plots compare pathway rank score between the original data and synthetic datasets of each SDG method. Rank scores combine the sign of the NES and the FDR, $Q$ value. Pathways located in the upper-right (UR, light green) and lower-left (LL, light purple) quadrants indicate concordant regulation direction and the level of significance (FDR, $Q < 0.05$).

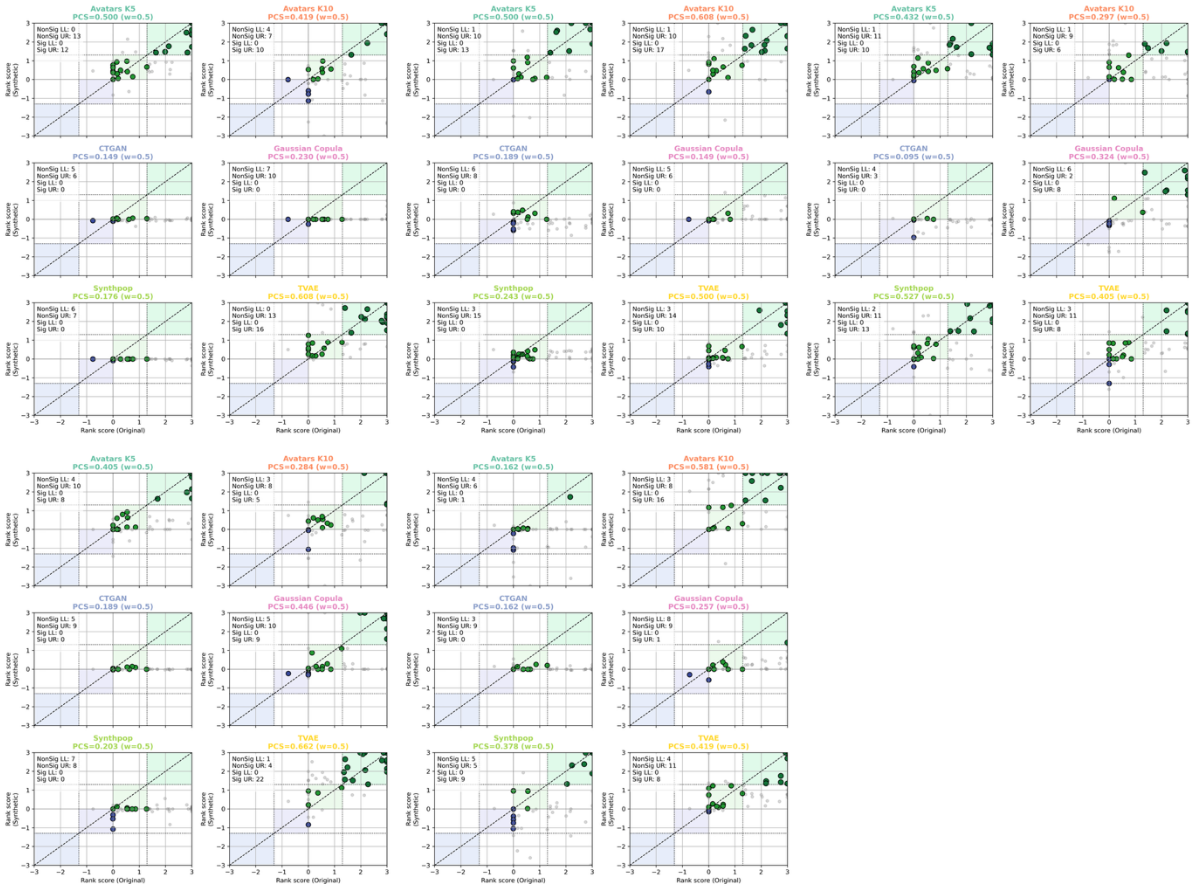

**Figure S13.** PCS scatter plots of Melanoma cohort across synthetic replicates. These scatter plots compare pathway rank score between the original data and synthetic datasets of each SDG method. Rank scores combine the sign of the NES and the FDR, $Q$ value. Pathways located in the upper-right (UR, light green) and lower-left (LL, light purple) quadrants indicate concordant regulation direction and the level of significance (FDR, $Q < 0.05$).

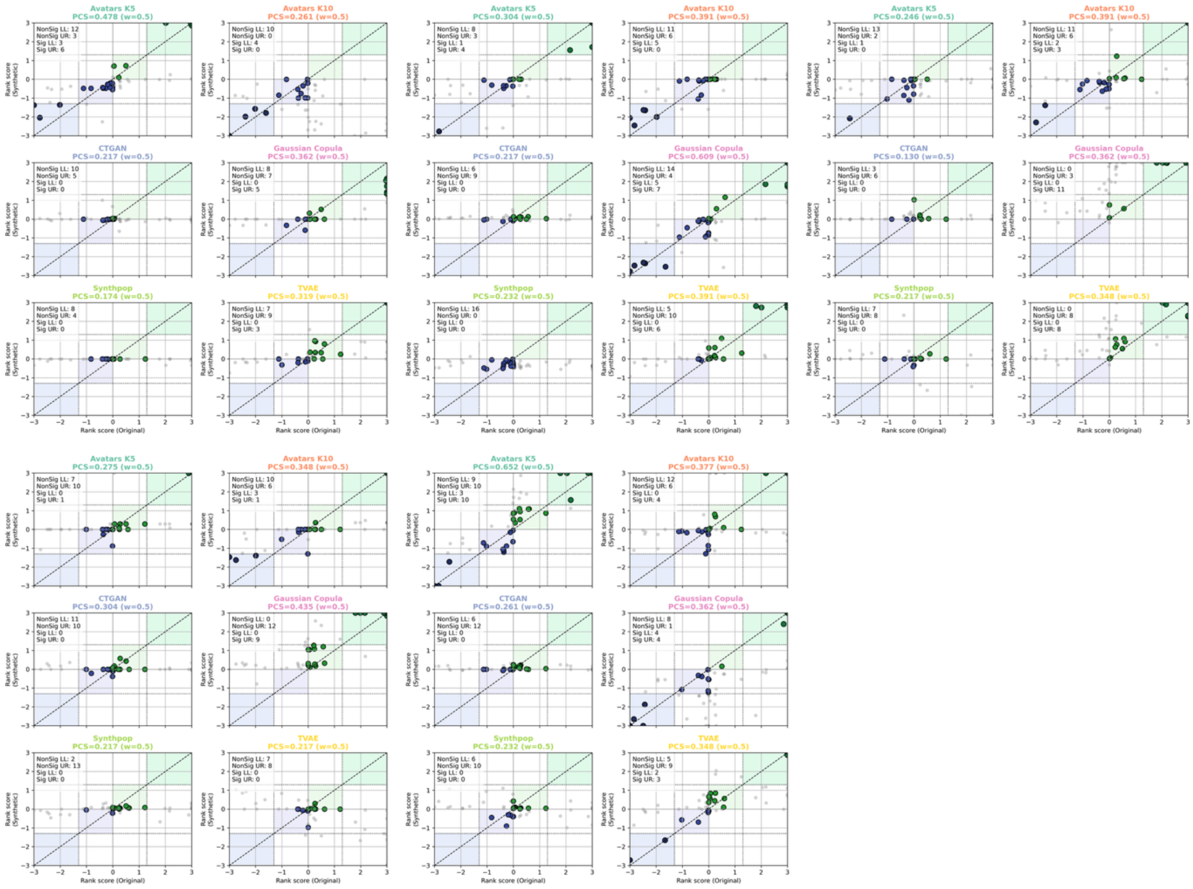

**Figure S14.** PCS scatter plots of NSCLC cohort across synthetic replicates. These scatter plots compare pathway rank score between the original data and synthetic datasets of each SDG method. Rank scores combine the sign of the NES and the FDR, $Q$ value. Pathways located in the upper-right (UR, light green) and lower-left (LL, light purple) quadrants indicate concordant regulation direction and the level of significance (FDR, $Q < 0.05$).

#### 3.3 Single-sample GSEA

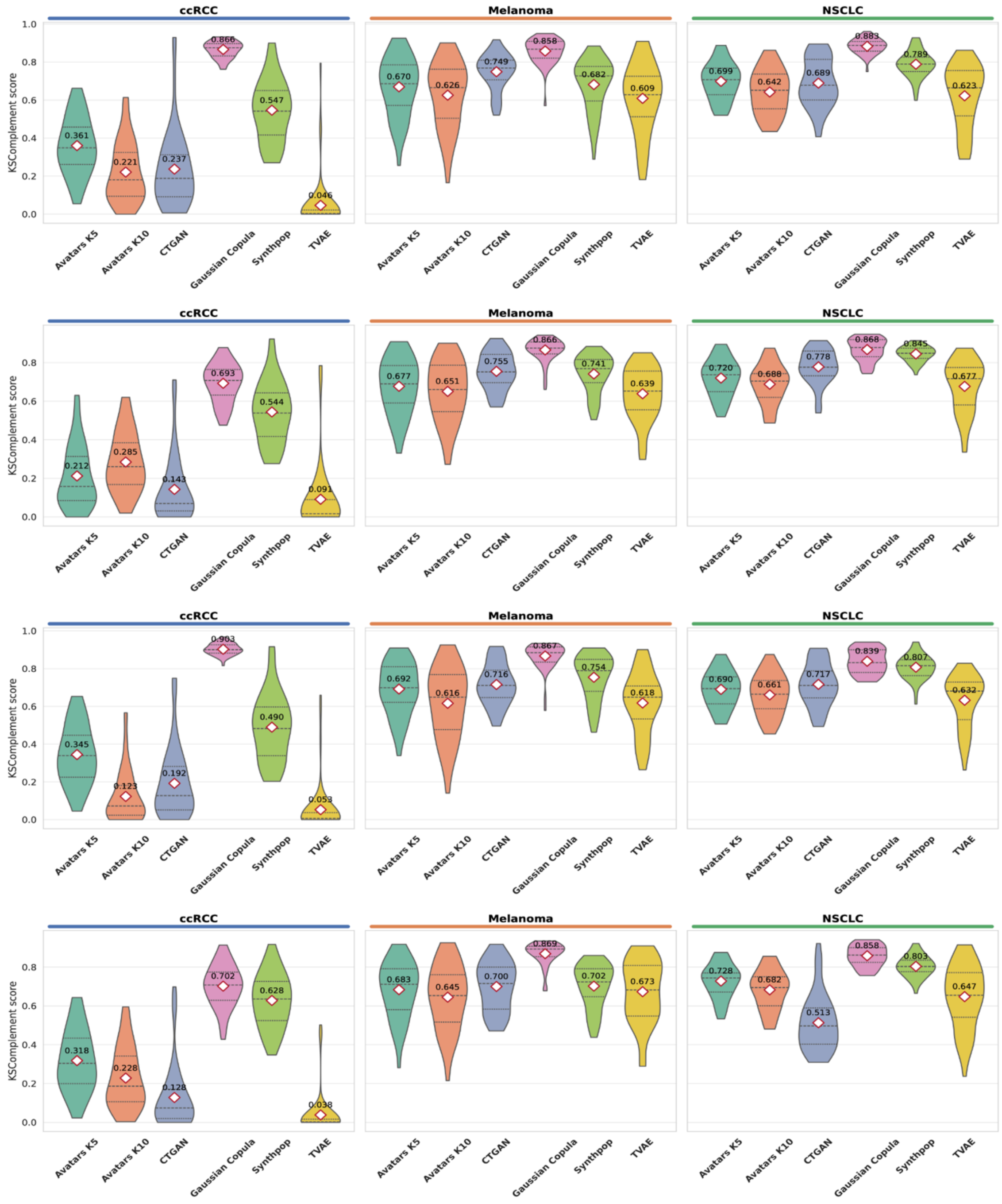

**Figure S15.** Distribution of KS-Complement scores of 50 Hallmark pathways across synthetic replicates. Violin plots summarize pathway-level KS-Complement values across datasets. Higher values reflect greater similarity of ssGSEA score distributions between original and synthetic data.

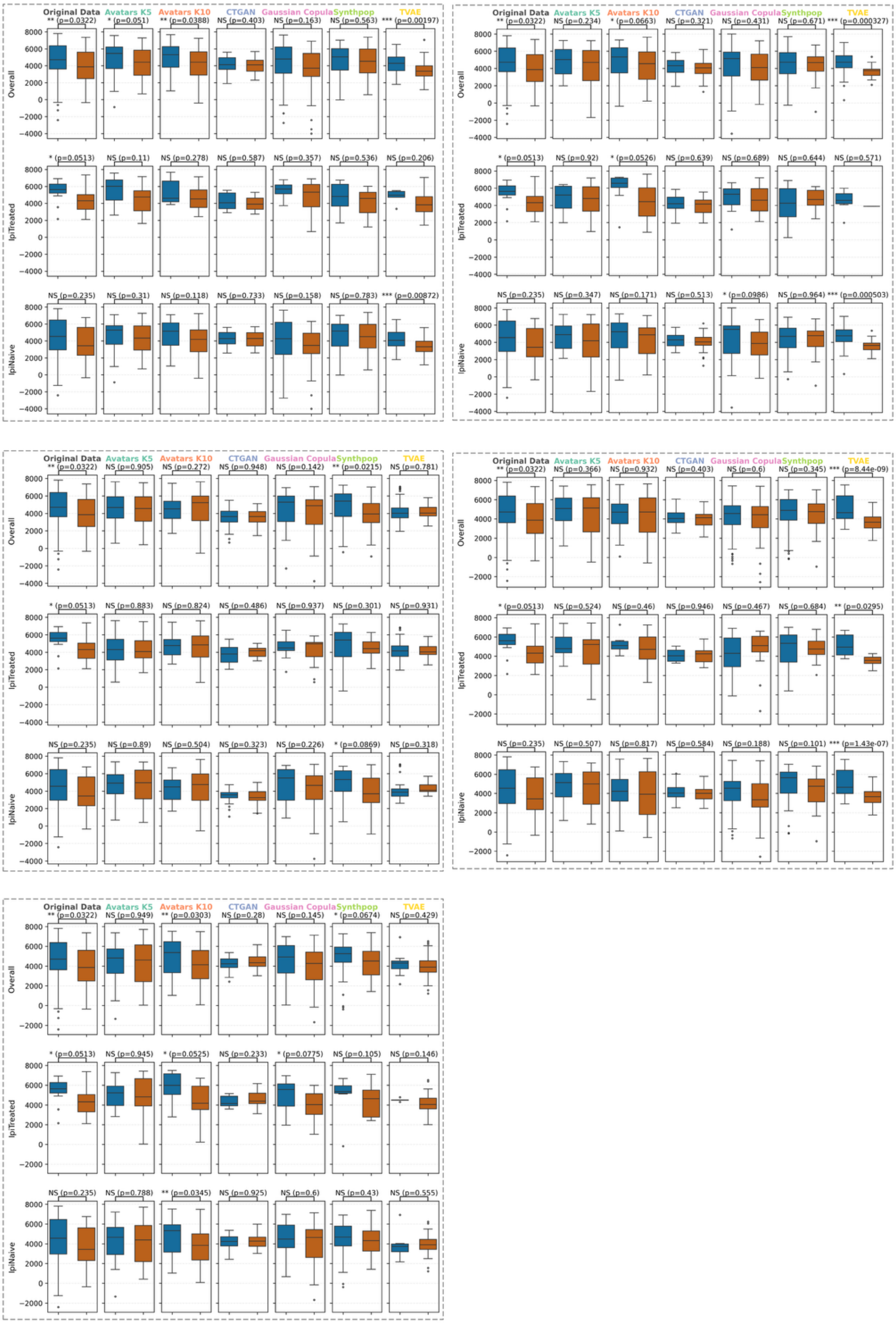

**Figure S16.** Comparison of MHC-II scores across clinical subgroups in original and synthetic Melanoma cohorts across multiple replicates. Boxplots compare MHC-II scores between responders (blue) and progressors (orange) across three clinical subgroups: Overall, Ipilimumab-treated (IpiTreated), and Ipilimumab-naïve (IpiNaive). The original data exhibited higher MHC-II scores in responders within the IpiTreated group ($\boldsymbol{P} < \mathbf{0.1}$)and comparable MHC-II scores between responders and progressors in the IpiNaive group ($\boldsymbol{P} > \mathbf{0.1}$). Avatars K10 and Gaussian Copula successfully captured the expected pattern, though this recovery lacks robustness across all replicates. Statistical significance was assessed using a two-sided Mann-Whitney U test.

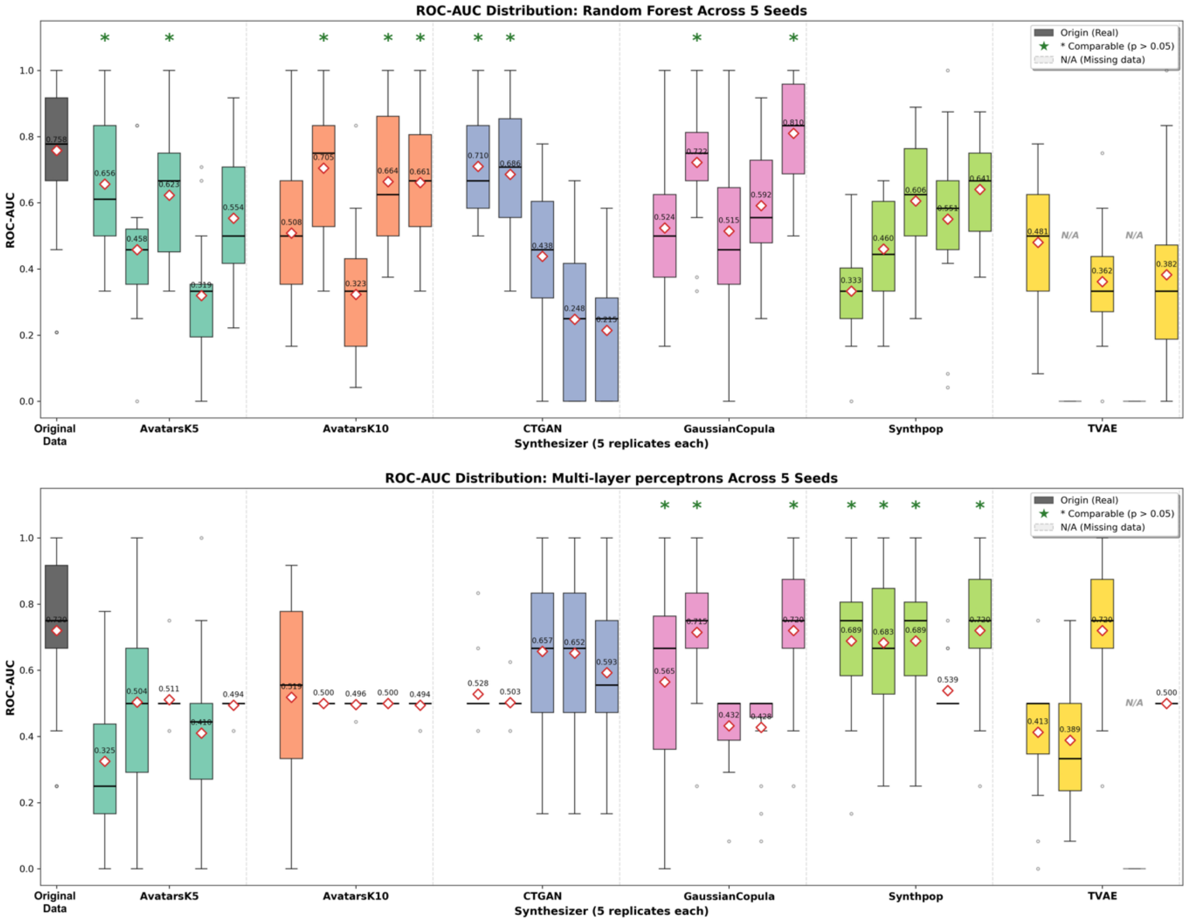

**Figure S17.** Predictive modeling transferability in Melanoma cohort. Boxplots depict cross-validated ROC-AUC for Random Forest and Multi-layer Perceptron models trained on synthetic data replicates and evaluated on held-out original data (5-fold cross-validation, repeated three times). The original data benchmark is shown for reference. Colored dots denote statistically non-inferior performance versus the original model (Wilcoxon signed-rank test, $P < 0.05$).

#### 3.4 Cell type deconvolution

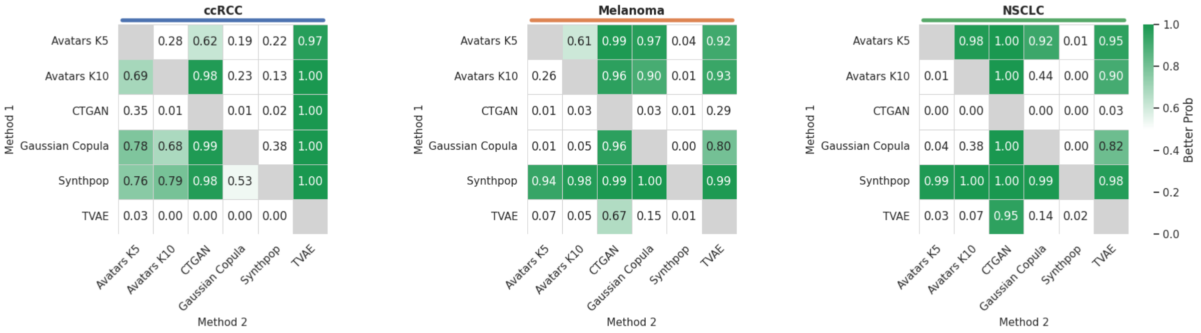

**Figure S18.** Bayesian comparison Aitchison distance score across cancers. Pairwise probability heatmaps display the posterior probability P(row>column) that a SDG method in a given row achieves a higher C-index score than the method in the corresponding column.

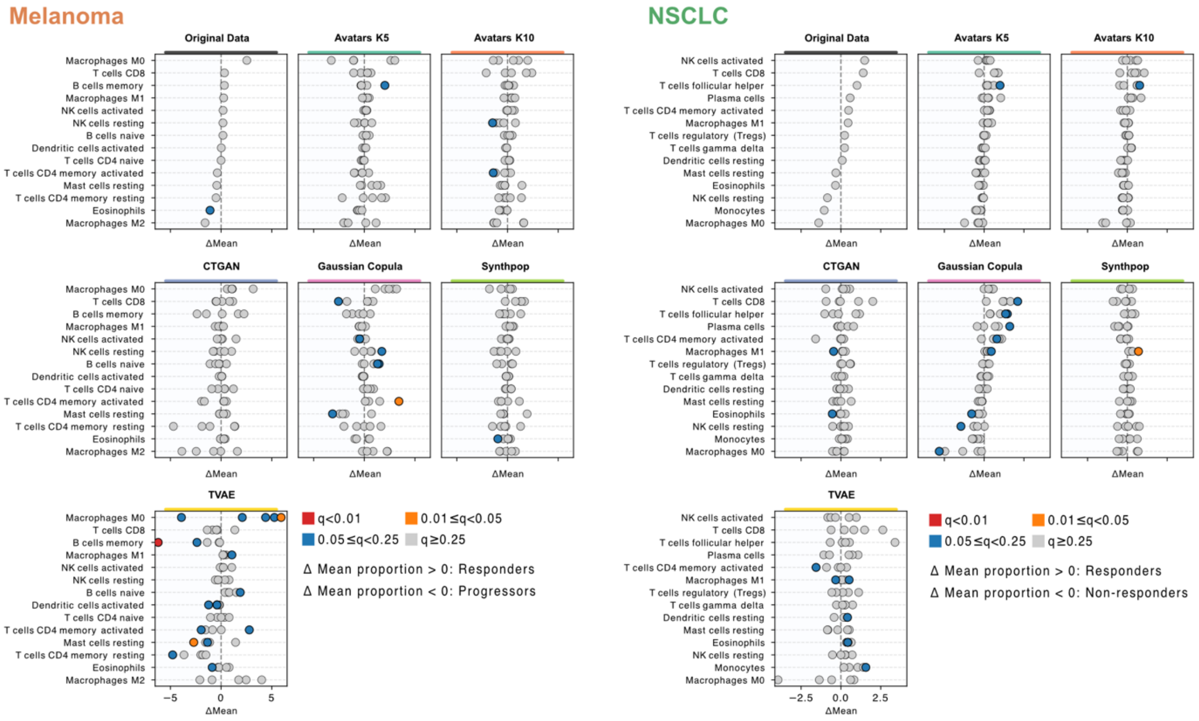

**Figure S19.** Differential LM22 immune cell proportions in Melanoma and NSCLC cohorts. Dot plots compare mean differences in immune cell fractions inferred by CIBERSORTx between Responders and Progressors/Non-responders across the original and synthetic datasets generated by each SDG method. Colors indicate statistical significance based on a two-sided Wilcoxon rank-sum test with FDR adjustment.

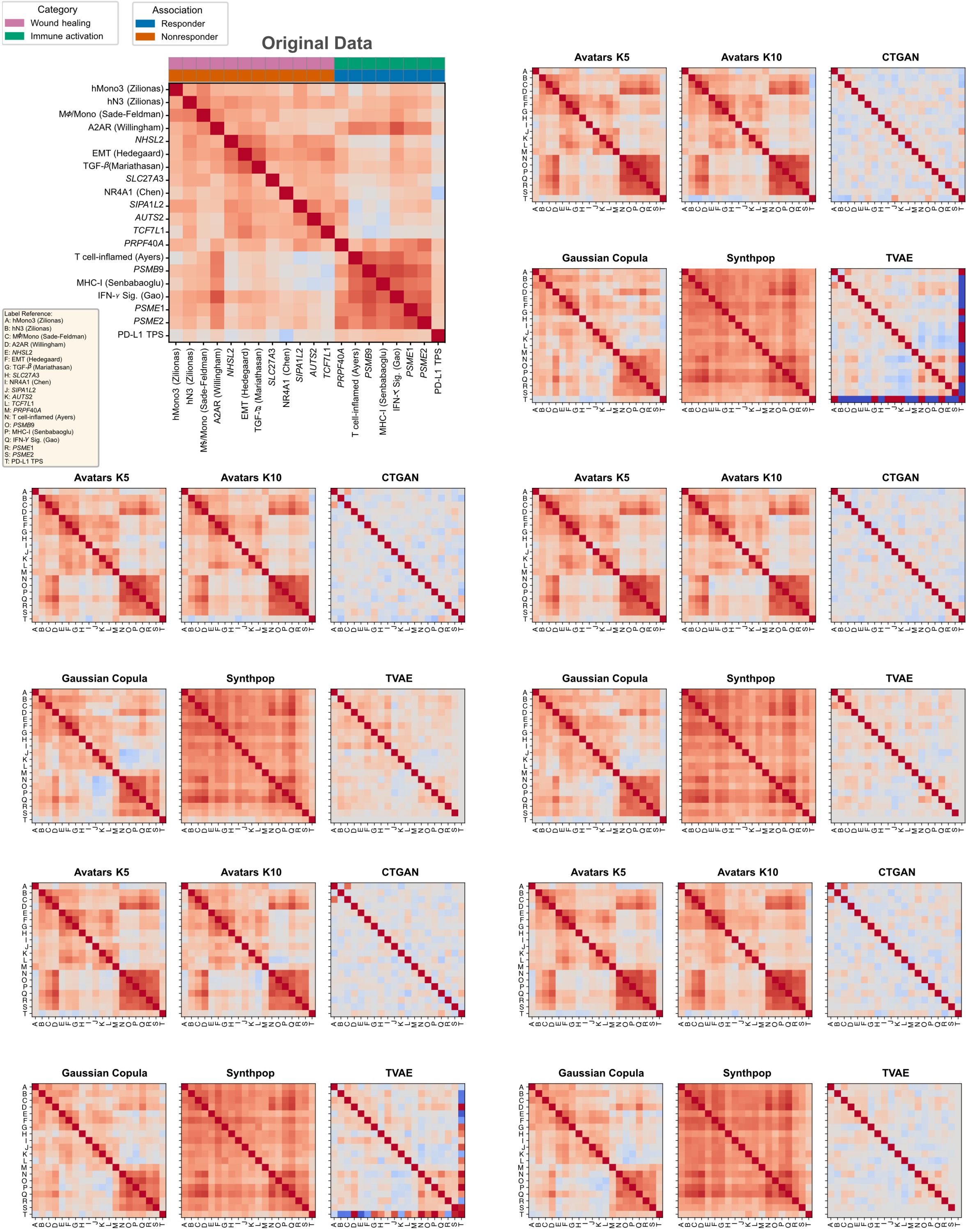

**Figure S20.** Biological signature correlation matrices in the NSCLC cohort across synthetic replicates. Correlation heatmaps display pairwise Pearson correlations among signatures in the original data and each synthetic dataset. The color scale ranges from negative (blue) to positive (red) correlations. Clustering highlights two principal modules: a resistance-associated wound-healing/immunosuppressive, stromal cluster (C1) and a response-associated immune activation/exhaustion cluster (C2).

#### 3.5 Survival analysis

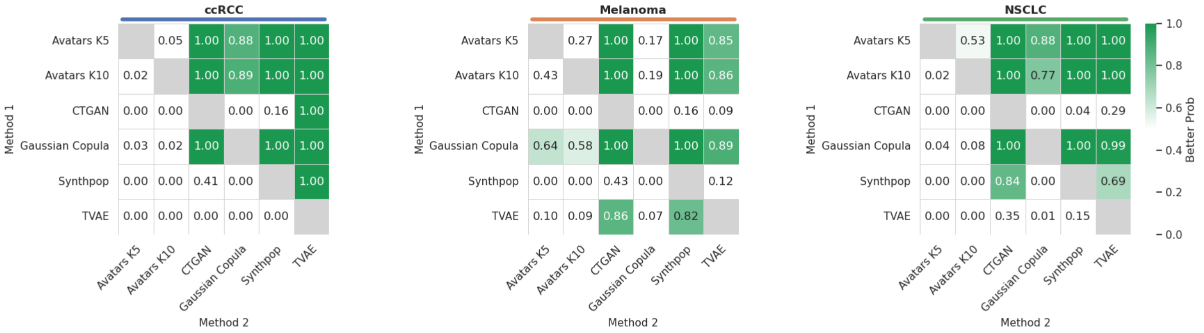

**Figure S21.** Bayesian comparison of C-index score across cancers. Pairwise probability heatmaps display the posterior probability $P(row > column)$ that a SDG method in a given row achieves a higher C-index score than the method in the corresponding column.

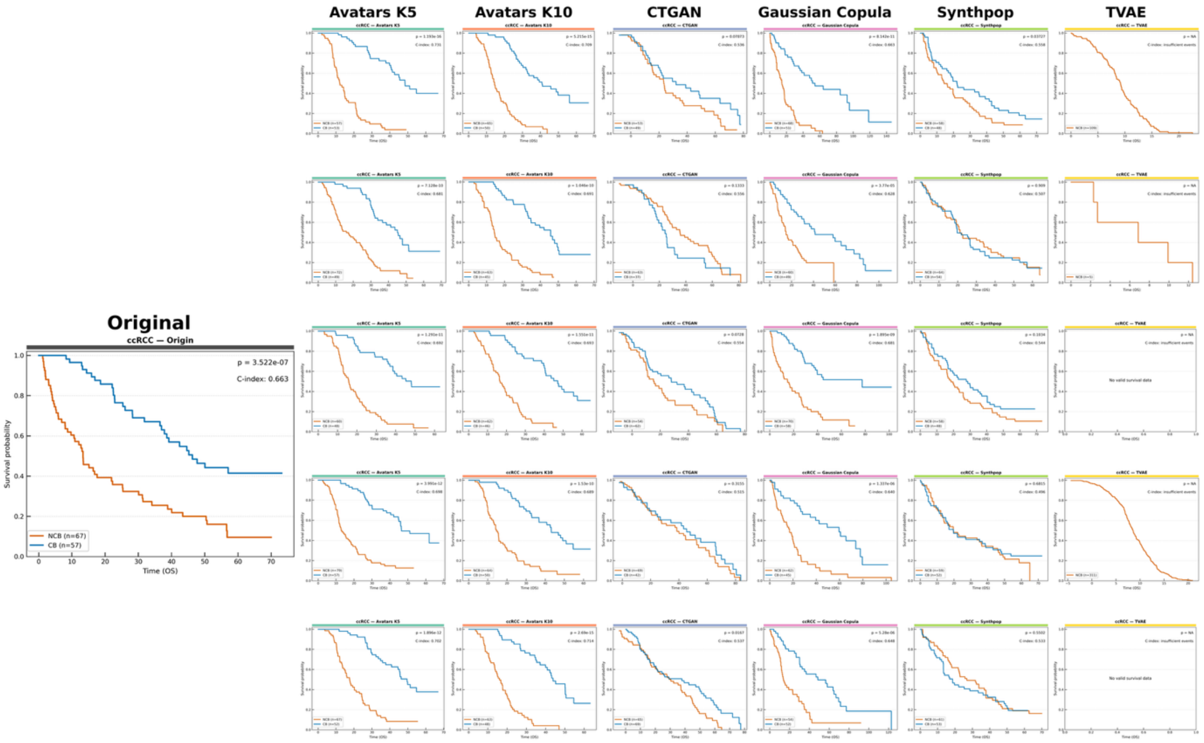

**Figure S22.** Kaplan-Meier Overall Survival (OS) curves of CB vs NCB in ccRCC cohorts between original data and synthetic replicates. Kaplan-Meier (KM) curves comparing OS between clinical benefit (CB) and non-clinical benefit (NCB) groups in patients with ccRCC treated with Nivolumab. The left panel shows the original cohort, while the remaining panels display results from multiple replicates of synthetic cohorts generated using six SDG methods. Survival probability is plotted over time (months), with blue lines representing CB and orange lines representing NCB. Group differences were evaluated using the log-rank test.

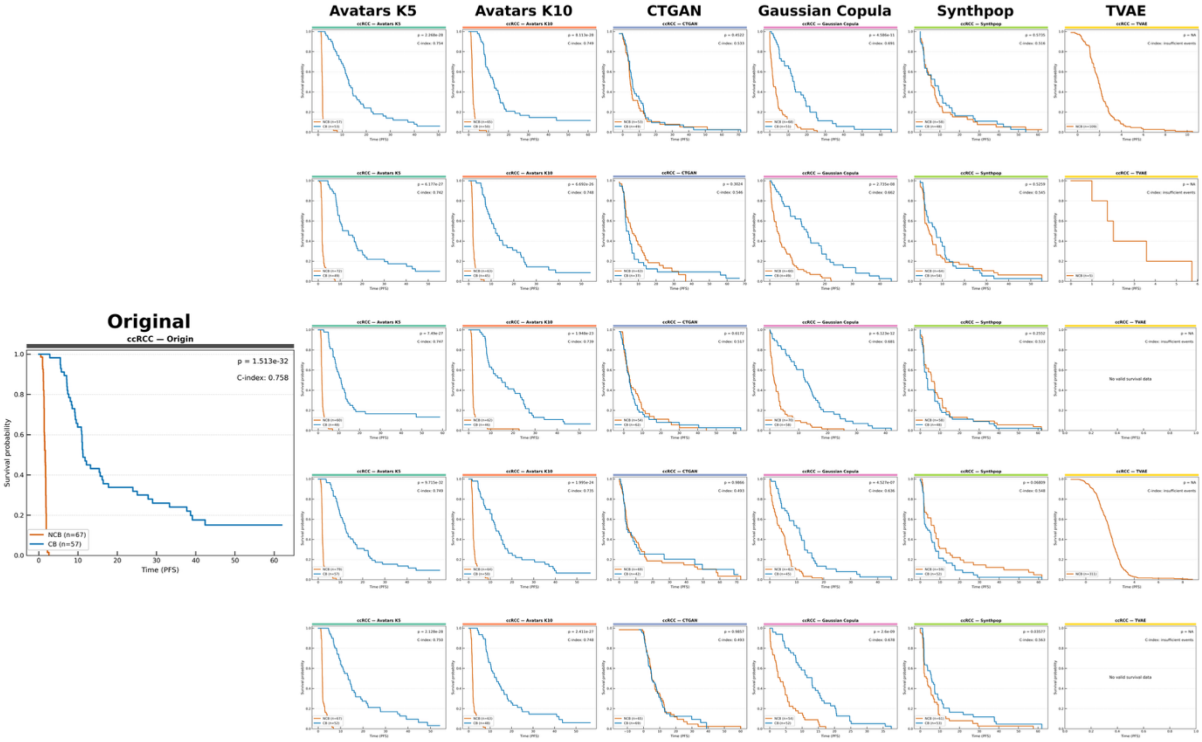

**Figure S23.** Kaplan-Meier Progression-Free Survival (PFS) curves of CB vs NCB in ccRCC cohorts between original data and synthetic replicates. Kaplan-Meier (KM) curves comparing PFS between clinical benefit (CB) and non–clinical benefit (NCB) groups in patients with ccRCC treated with Nivolumab. Group differences were evaluated using the log-rank test.

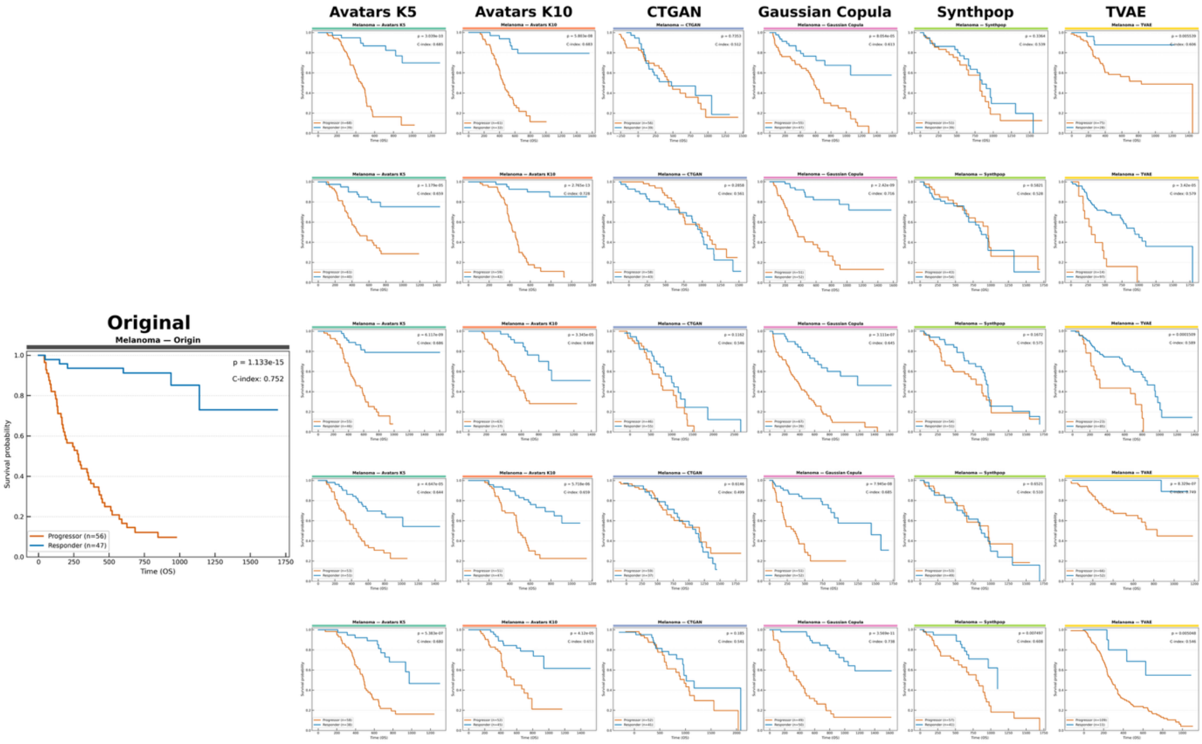

**Figure S24.** Kaplan-Meier Overall Survival (OS) curves of Responders vs Progressors in Melanoma cohorts between original data and synthetic replicates. Kaplan-Meier (KM) curves comparing OS between Responders (CR/PR) and Progressors (PD) groups in patients with Melanoma. Group differences were evaluated using the log-rank test.

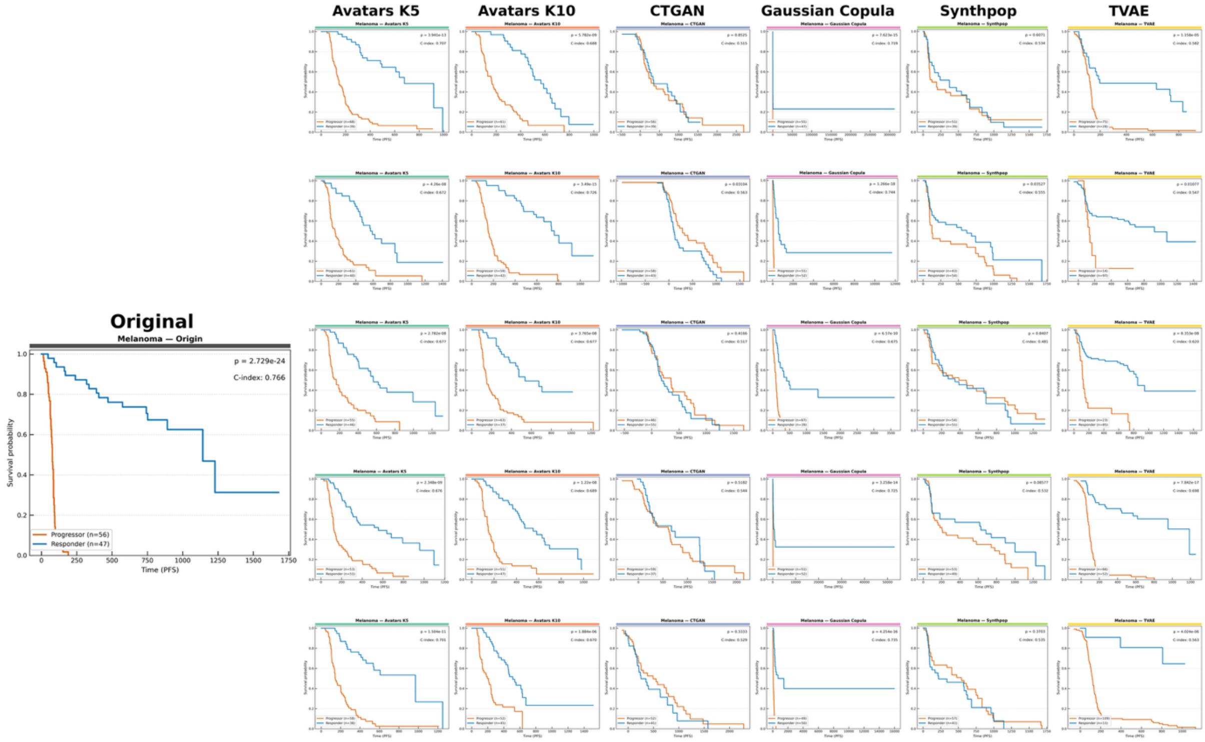

**Figure S25.** Kaplan-Meier Progression-Free Survival (PFS) curves of Responders vs Progressors in Melanoma cohorts between original data and synthetic replicates. Kaplan-Meier (KM) curves comparing PFS between Responders (CR/PR) and Progressors (PD) groups in patients with Melanoma. Group differences were evaluated using the log-rank test.

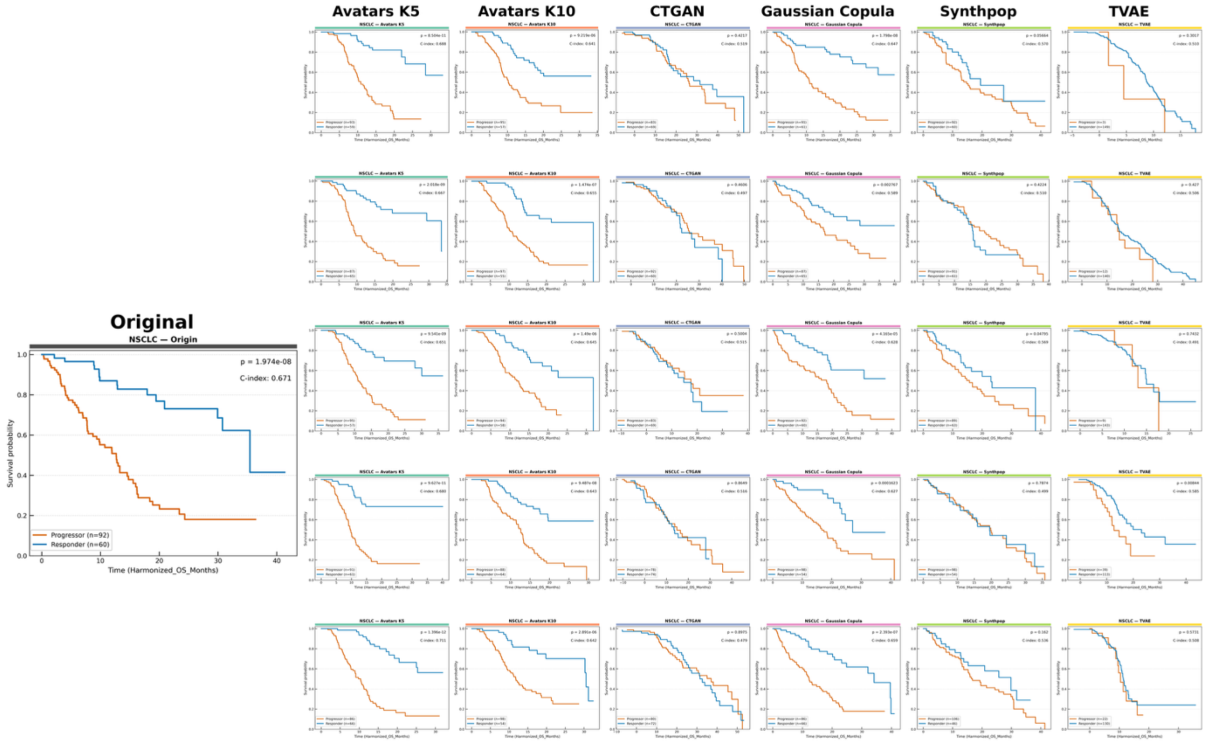

**Figure S26.** Kaplan-Meier Overall Survival (OS) curves of Responders vs non-Responders in NSCLC cohorts between original data and synthetic replicates. Kaplan-Meier (KM) curves comparing OS between Responders (CR/PR) and non-Responders (PD/SD) groups in patients with NSCLC. Group differences were evaluated using the log-rank test.

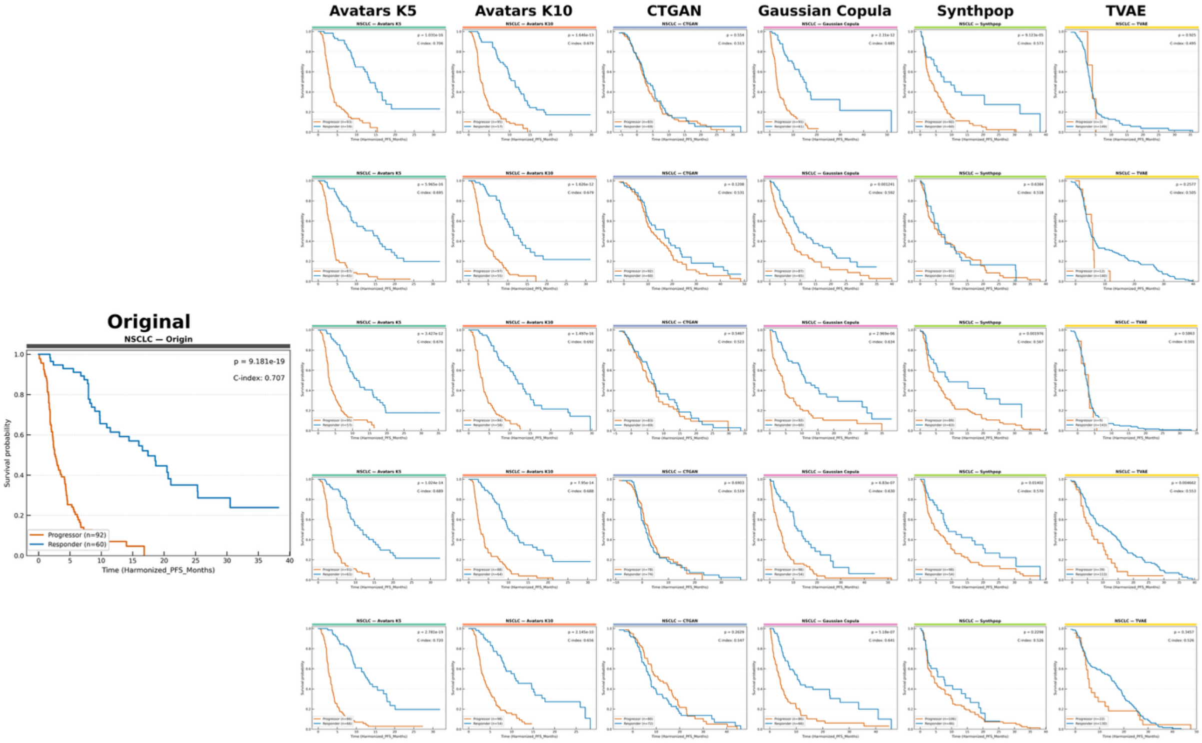

**Figure S27.** Kaplan-Meier Progression-Free Survival (PFS) curves of Responders vs non-Responders in NSCLC cohorts between original data and synthetic replicates. Kaplan-Meier (KM) curves comparing PFS between Responders (CR/PR) and non-Responders (PD/SD) groups in patients with NSCLC. Group differences were evaluated using the log-rank test.

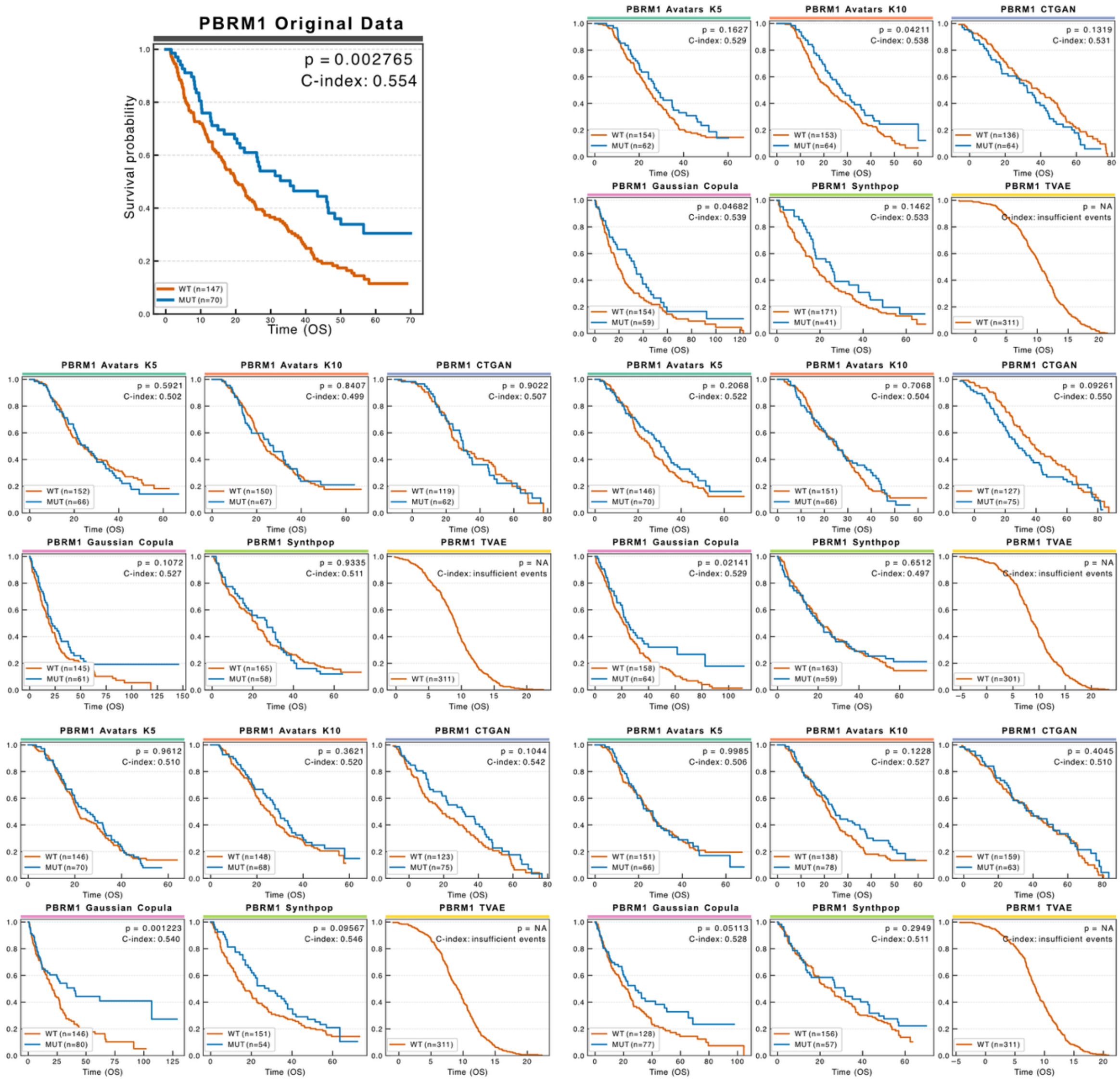

**Figure S28.** PBRM1-associated overall survival in ccRCC across synthetic replicates. Kaplan-Meier curves compare OS between PBRM1-mutant (blue) and wild-type (orange) groups in the original dataset and across synthetic cohorts generated by each SDG method. The original survival advantage of PBRM1-mutant patients is reproduced in selected synthetic datasets, with log-rank $P$ values C-indices reported in each panel.

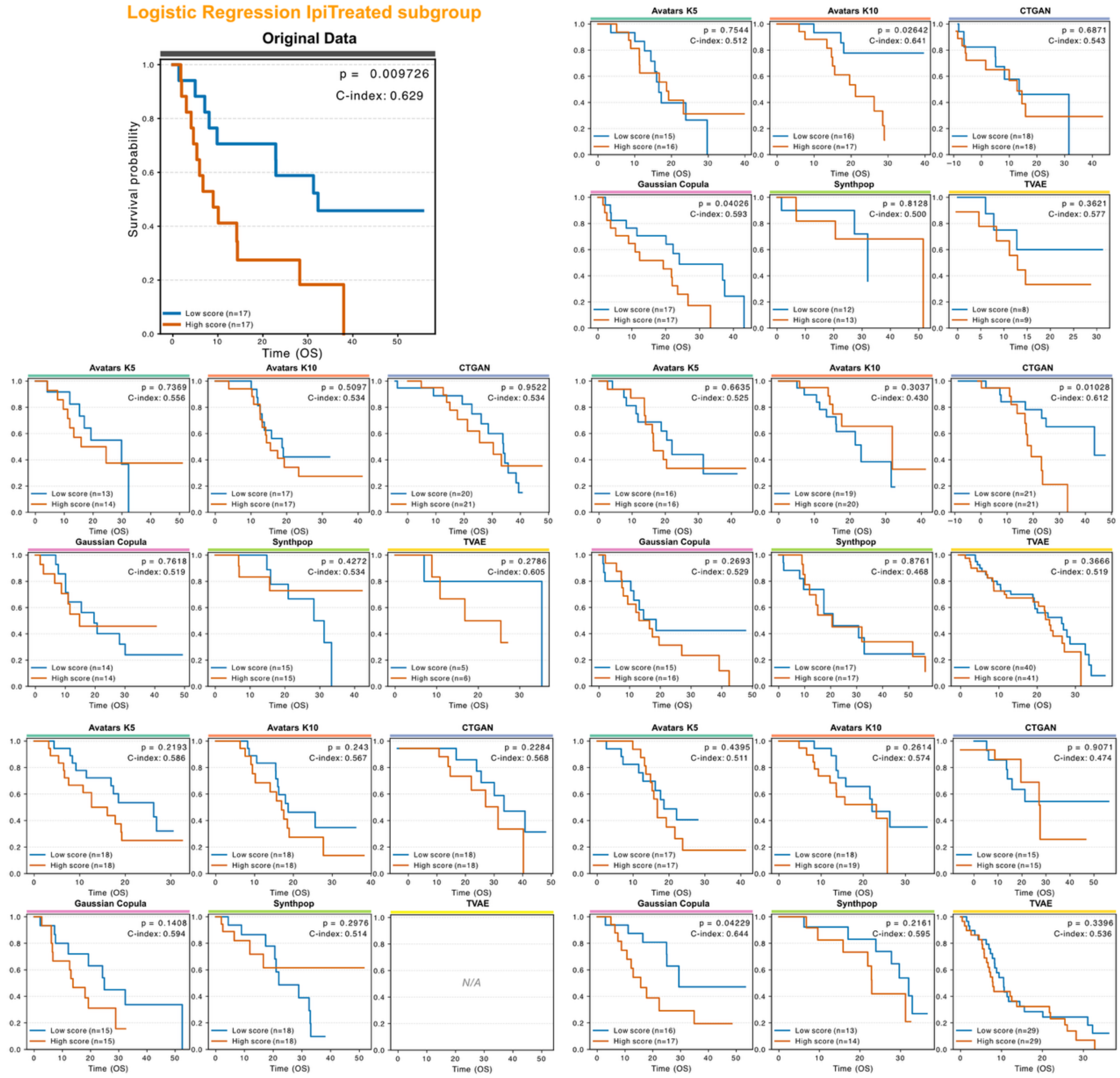

**Figure S29.** Preservation of survival stratification based on a predictive model in the ipilimumab-treated Melanoma cohort across synthetic replicates. Kaplan-Meier curves of OS stratified by predicted risk groups (low risk: blue; high risk: orange) derived from a logistic regression model classifying progressors versus non-progressors. The model was retrained on each synthetic dataset, and predicted risk scores were used for survival stratification. Log-rank $P$ values and C-indices are reported in each panel.

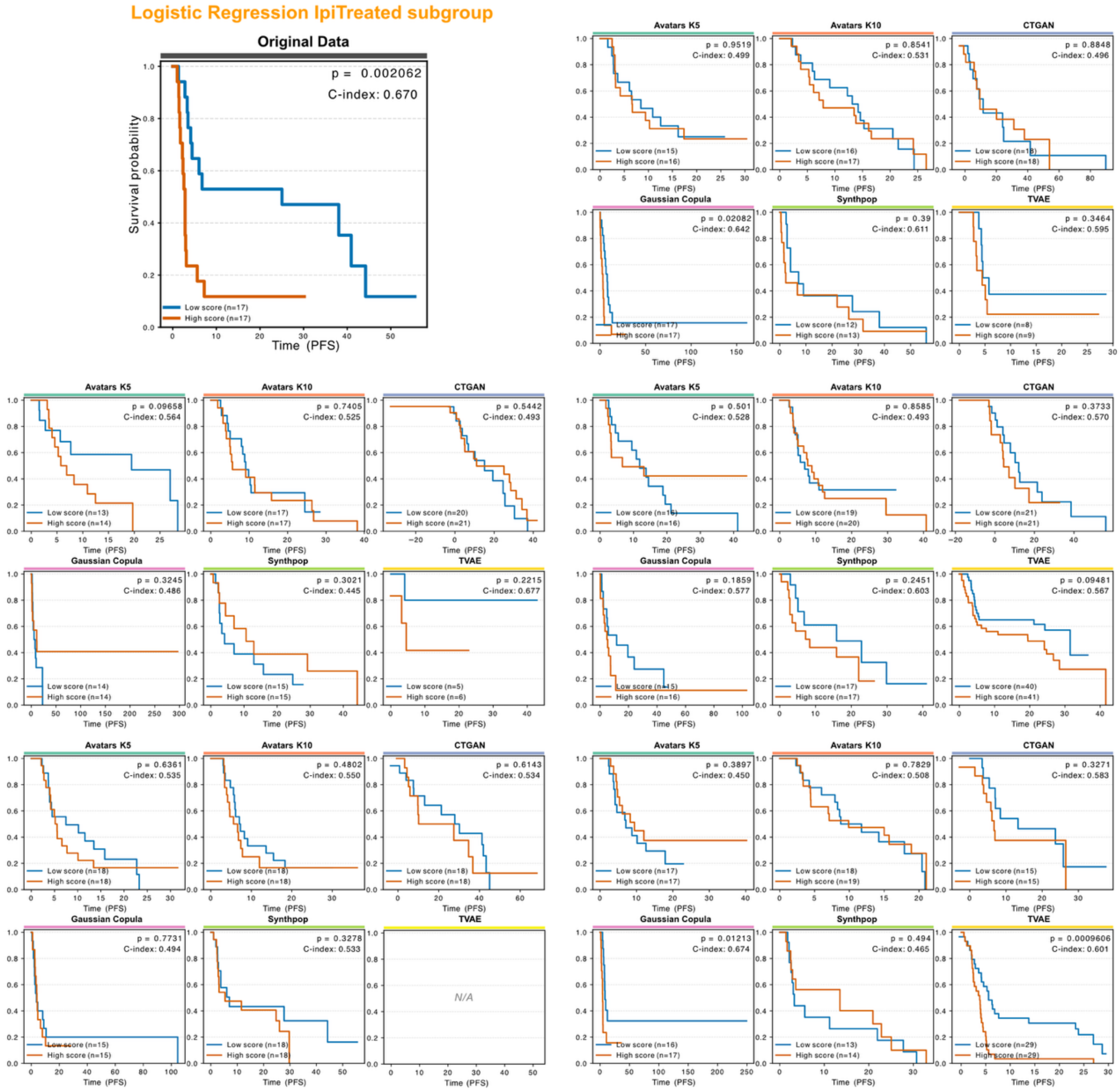

**Figure S30.** Preservation of survival stratification based on a predictive model in the ipilimumab-treated Melanoma cohort across synthetic replicates. Kaplan-Meier curves of PFS stratified by predicted risk groups (low risk: blue; high risk: orange) derived from a logistic regression model classifying progressors versus non-progressors. The model was retrained on each synthetic dataset, and predicted risk scores were used for survival stratification. Log-rank $P$ values and C-indices are reported in each panel.

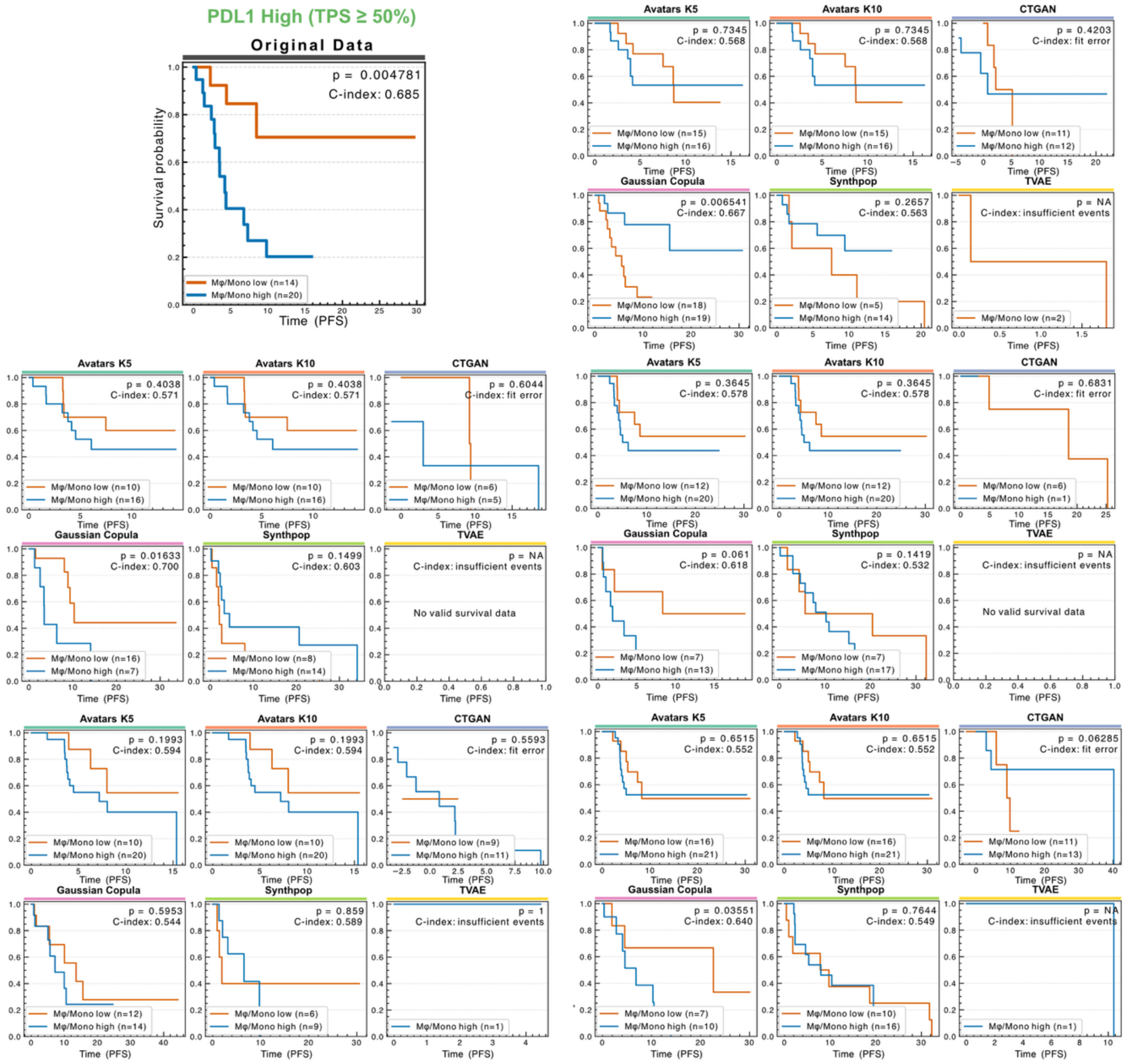

**Figure S31.** Patient stratification using a macrophage/monocyte signature in the PD-L1 high NSCLC cohort across synthetic replicates. Kaplan-Meier curves evaluate PFS differences between tumors with low (orange) versus high (blue) macrophage/monocyte aignaturescores within the PD-L1 high (TPS ≥ 50%) subgroup. Recovery of the adverse prognostic association observed in the original dataset is assessed across SDG methods. Log-rank P values and C-indices are shown in each panel.

### C. Supplementary Table

#### 1. Broad utility

##### 1.1 Univariate similarity

**Table S1.** Pairwise Bayesian comparison of SDG methods based on univariate similarity scores in the ccRCC dataset.

| Method 1 | Method 2 | Better Prob | Worse Prob | Equivalent Prob |
|---|---|---|---|---|
| Avatars K5 | Avatars K10 | 0.998 | 0.000 | 0.002 |
| Avatars K5 | CTGAN | 0.999 | 0.000 | 0.001 |
| Avatars K5 | Gaussian Copula | 0.025 | 0.013 | 0.962 |
| Avatars K5 | Synthpop | 0.000 | 1.000 | 0.000 |
| Avatars K5 | TVAE | 1.000 | 0.000 | 0.000 |
| Avatars K10 | Avatars K5 | 0.000 | 0.998 | 0.002 |
| Avatars K10 | CTGAN | 0.996 | 0.001 | 0.003 |
| Avatars K10 | Gaussian Copula | 0.001 | 0.973 | 0.026 |
| Avatars K10 | Synthpop | 0.000 | 1.000 | 0.000 |
| Avatars K10 | TVAE | 1.000 | 0.000 | 0.000 |
| CTGAN | Avatars K5 | 0.000 | 0.999 | 0.001 |
| CTGAN | Avatars K10 | 0.001 | 0.996 | 0.003 |
| CTGAN | Gaussian Copula | 0.000 | 1.000 | 0.000 |
| CTGAN | Synthpop | 0.000 | 1.000 | 0.000 |
| CTGAN | TVAE | 1.000 | 0.000 | 0.000 |
| Gaussian Copula | Avatars K5 | 0.013 | 0.025 | 0.962 |
| Gaussian Copula | Avatars K10 | 0.973 | 0.001 | 0.026 |
| Gaussian Copula | CTGAN | 1.000 | 0.000 | 0.000 |
| Gaussian Copula | Synthpop | 0.000 | 1.000 | 0.000 |
| Gaussian Copula | TVAE | 1.000 | 0.000 | 0.000 |
| Synthpop | Avatars K5 | 1.000 | 0.000 | 0.000 |
| Synthpop | Avatars K10 | 1.000 | 0.000 | 0.000 |
| Synthpop | CTGAN | 1.000 | 0.000 | 0.000 |
| Synthpop | Gaussian Copula | 1.000 | 0.000 | 0.000 |
| Synthpop | TVAE | 1.000 | 0.000 | 0.000 |

| | | | | |
|---|---|---|---|---|
| TVAE | Avatars K5 | 0.000 | 1.000 | 0.000 |
| TVAE | Avatars K10 | 0.000 | 1.000 | 0.000 |
| TVAE | CTGAN | 0.000 | 1.000 | 0.000 |
| TVAE | Gaussian Copula | 0.000 | 1.000 | 0.000 |
| TVAE | Synthpop | 0.000 | 1.000 | 0.000 |

**Table S2.** Pairwise Bayesian comparison of SDG methods based on univariate similarity scores in the Melanoma dataset.

| Method 1 | Method 2 | Better Prob | Worse Prob | Equivalent Prob |
|---|---|---|---|---|
| Avatars K5 | Avatars K10 | 0.942 | 0.000 | 0.058 |
| Avatars K5 | CTGAN | 0.999 | 0.000 | 0.001 |
| Avatars K5 | Gaussian Copula | 0.000 | 1.000 | 0.000 |
| Avatars K5 | Synthpop | 0.000 | 1.000 | 0.000 |
| Avatars K5 | TVAE | 0.988 | 0.004 | 0.009 |
| Avatars K10 | Avatars K5 | 0.000 | 0.942 | 0.058 |
| Avatars K10 | CTGAN | 0.997 | 0.001 | 0.002 |
| Avatars K10 | Gaussian Copula | 0.000 | 1.000 | 0.000 |
| Avatars K10 | Synthpop | 0.000 | 1.000 | 0.000 |
| Avatars K10 | TVAE | 0.966 | 0.007 | 0.026 |
| CTGAN | Avatars K5 | 0.000 | 0.999 | 0.001 |
| CTGAN | Avatars K10 | 0.001 | 0.997 | 0.002 |
| CTGAN | Gaussian Copula | 0.000 | 1.000 | 0.000 |
| CTGAN | Synthpop | 0.000 | 1.000 | 0.000 |
| CTGAN | TVAE | 0.055 | 0.843 | 0.102 |
| Gaussian Copula | Avatars K5 | 1.000 | 0.000 | 0.000 |
| Gaussian Copula | Avatars K10 | 1.000 | 0.000 | 0.000 |
| Gaussian Copula | CTGAN | 1.000 | 0.000 | 0.000 |
| Gaussian Copula | Synthpop | 0.000 | 0.998 | 0.002 |
| Gaussian Copula | TVAE | 0.998 | 0.001 | 0.001 |
| Synthpop | Avatars K5 | 1.000 | 0.000 | 0.000 |
| Synthpop | Avatars K10 | 1.000 | 0.000 | 0.000 |

| | | | | |
|---|---|---|---|---|
| Synthpop | CTGAN | 1.000 | 0.000 | 0.000 |
| Synthpop | Gaussian Copula | 0.998 | 0.000 | 0.002 |
| Synthpop | TVAE | 0.999 | 0.000 | 0.000 |
| TVAE | Avatars K5 | 0.004 | 0.988 | 0.009 |
| TVAE | Avatars K10 | 0.007 | 0.966 | 0.026 |
| TVAE | CTGAN | 0.843 | 0.055 | 0.102 |
| TVAE | Gaussian Copula | 0.001 | 0.998 | 0.001 |
| TVAE | Synthpop | 0.000 | 0.999 | 0.000 |

**Table S3.** Pairwise Bayesian comparison of SDG methods based on univariate similarity scores in the NSCLC dataset.

| Method 1 | Method 2 | Better Prob | Worse Prob | Equivalent Prob |
|---|---|---|---|---|
| Avatars K5 | Avatars K10 | 0.893 | 0.001 | 0.107 |
| Avatars K5 | CTGAN | 0.998 | 0.001 | 0.001 |
| Avatars K5 | Gaussian Copula | 0.001 | 0.994 | 0.005 |
| Avatars K5 | Synthpop | 0.000 | 1.000 | 0.000 |
| Avatars K5 | TVAE | 0.999 | 0.001 | 0.001 |
| Avatars K10 | Avatars K5 | 0.001 | 0.893 | 0.107 |
| Avatars K10 | CTGAN | 0.996 | 0.002 | 0.002 |
| Avatars K10 | Gaussian Copula | 0.000 | 1.000 | 0.000 |
| Avatars K10 | Synthpop | 0.000 | 1.000 | 0.000 |
| Avatars K10 | TVAE | 0.998 | 0.001 | 0.001 |
| CTGAN | Avatars K5 | 0.001 | 0.998 | 0.001 |
| CTGAN | Avatars K10 | 0.002 | 0.996 | 0.002 |
| CTGAN | Gaussian Copula | 0.001 | 0.999 | 0.000 |
| CTGAN | Synthpop | 0.000 | 1.000 | 0.000 |
| CTGAN | TVAE | 0.556 | 0.224 | 0.220 |
| Gaussian Copula | Avatars K5 | 0.994 | 0.001 | 0.005 |
| Gaussian Copula | Avatars K10 | 1.000 | 0.000 | 0.000 |
| Gaussian Copula | CTGAN | 0.999 | 0.001 | 0.000 |
| Gaussian Copula | Synthpop | 0.002 | 0.955 | 0.042 |

| | | | | |
|---|---|---|---|---|
| Gaussian Copula | TVAE | 1.000 | 0.000 | 0.000 |
| Synthpop | Avatars K5 | 1.000 | 0.000 | 0.000 |
| Synthpop | Avatars K10 | 1.000 | 0.000 | 0.000 |
| Synthpop | CTGAN | 1.000 | 0.000 | 0.000 |
| Synthpop | Gaussian Copula | 0.955 | 0.002 | 0.042 |
| Synthpop | TVAE | 1.000 | 0.000 | 0.000 |
| TVAE | Avatars K5 | 0.001 | 0.999 | 0.001 |
| TVAE | Avatars K10 | 0.001 | 0.998 | 0.001 |
| TVAE | CTGAN | 0.224 | 0.556 | 0.220 |
| TVAE | Gaussian Copula | 0.000 | 1.000 | 0.000 |
| TVAE | Synthpop | 0.000 | 1.000 | 0.000 |

#### 1.2 Bivariate similarity

**Table S4.** Pairwise Bayesian comparison of SDG methods based on bivariate similarity scores in the ccRCC dataset.

| Method 1 | Method 2 | Better Prob | Worse Prob | Equivalent Prob |
|---|---|---|---|---|
| Avatars K5 | Avatars K10 | 0.000 | 0.000 | 1.000 |
| Avatars K5 | CTGAN | 1.000 | 0.000 | 0.000 |
| Avatars K5 | Gaussian Copula | 0.998 | 0.000 | 0.002 |
| Avatars K5 | Synthpop | 1.000 | 0.000 | 0.000 |
| Avatars K5 | TVAE | 1.000 | 0.000 | 0.000 |
| Avatars K10 | Avatars K5 | 0.000 | 0.000 | 1.000 |
| Avatars K10 | CTGAN | 1.000 | 0.000 | 0.000 |
| Avatars K10 | Gaussian Copula | 0.982 | 0.000 | 0.018 |
| Avatars K10 | Synthpop | 1.000 | 0.000 | 0.000 |
| Avatars K10 | TVAE | 1.000 | 0.000 | 0.000 |
| CTGAN | Avatars K5 | 0.000 | 1.000 | 0.000 |
| CTGAN | Avatars K10 | 0.000 | 1.000 | 0.000 |
| CTGAN | Gaussian Copula | 0.000 | 1.000 | 0.000 |
| CTGAN | Synthpop | 0.000 | 1.000 | 0.000 |
| CTGAN | TVAE | 0.991 | 0.002 | 0.007 |
| Gaussian Copula | Avatars K5 | 0.000 | 0.998 | 0.002 |
| Gaussian Copula | Avatars K10 | 0.000 | 0.982 | 0.018 |

| | | | | |
|---|---|---|---|---|
| Gaussian Copula | CTGAN | 1.000 | 0.000 | 0.000 |
| Gaussian Copula | Synthpop | 1.000 | 0.000 | 0.000 |
| Gaussian Copula | TVAE | 1.000 | 0.000 | 0.000 |
| Synthpop | Avatars K5 | 0.000 | 1.000 | 0.000 |
| Synthpop | Avatars K10 | 0.000 | 1.000 | 0.000 |
| Synthpop | CTGAN | 1.000 | 0.000 | 0.000 |
| Synthpop | Gaussian Copula | 0.000 | 1.000 | 0.000 |
| Synthpop | TVAE | 1.000 | 0.000 | 0.000 |
| TVAE | Avatars K5 | 0.000 | 1.000 | 0.000 |
| TVAE | Avatars K10 | 0.000 | 1.000 | 0.000 |
| TVAE | CTGAN | 0.002 | 0.991 | 0.007 |
| TVAE | Gaussian Copula | 0.000 | 1.000 | 0.000 |
| TVAE | Synthpop | 0.000 | 1.000 | 0.000 |

**Table S5.** Pairwise Bayesian comparison of SDG methods based on bivariate similarity scores in the Melanoma dataset.

| Method 1 | Method 2 | Better Prob | Worse Prob | Equivalent Prob |
|---|---|---|---|---|
| Avatars K5 | Avatars K10 | 0.000 | 0.000 | 1.000 |
| Avatars K5 | CTGAN | 1.000 | 0.000 | 0.000 |
| Avatars K5 | Gaussian Copula | 0.000 | 0.002 | 0.998 |
| Avatars K5 | Synthpop | 0.280 | 0.000 | 0.720 |
| Avatars K5 | TVAE | 0.997 | 0.000 | 0.003 |
| Avatars K10 | Avatars K5 | 0.000 | 0.000 | 1.000 |
| Avatars K10 | CTGAN | 1.000 | 0.000 | 0.000 |
| Avatars K10 | Gaussian Copula | 0.000 | 0.000 | 1.000 |
| Avatars K10 | Synthpop | 0.022 | 0.000 | 0.978 |
| Avatars K10 | TVAE | 0.996 | 0.000 | 0.003 |
| CTGAN | Avatars K5 | 0.000 | 1.000 | 0.000 |
| CTGAN | Avatars K10 | 0.000 | 1.000 | 0.000 |
| CTGAN | Gaussian Copula | 0.000 | 1.000 | 0.000 |
| CTGAN | Synthpop | 0.000 | 0.164 | 0.836 |
| CTGAN | TVAE | 0.309 | 0.002 | 0.689 |

| | | | | |
|---|---|---|---|---|
| Gaussian Copula | Avatars K5 | 0.002 | 0.000 | 0.998 |
| Gaussian Copula | Avatars K10 | 0.000 | 0.000 | 1.000 |
| Gaussian Copula | CTGAN | 1.000 | 0.000 | 0.000 |
| Gaussian Copula | Synthpop | 0.975 | 0.000 | 0.025 |
| Gaussian Copula | TVAE | 0.999 | 0.000 | 0.001 |
| Synthpop | Avatars K5 | 0.000 | 0.280 | 0.720 |
| Synthpop | Avatars K10 | 0.000 | 0.022 | 0.978 |
| Synthpop | CTGAN | 0.164 | 0.000 | 0.836 |
| Synthpop | Gaussian Copula | 0.000 | 0.975 | 0.025 |
| Synthpop | TVAE | 0.963 | 0.000 | 0.037 |
| TVAE | Avatars K5 | 0.000 | 0.997 | 0.003 |
| TVAE | Avatars K10 | 0.000 | 0.996 | 0.003 |
| TVAE | CTGAN | 0.002 | 0.309 | 0.689 |
| TVAE | Gaussian Copula | 0.000 | 0.999 | 0.001 |
| TVAE | Synthpop | 0.000 | 0.963 | 0.037 |

**Table S6.** Pairwise Bayesian comparison of SDG methods based on bivariate similarity scores in the NSCLC dataset.

| Method 1 | Method 2 | Better Prob | Worse Prob | Equivalent Prob |
|---|---|---|---|---|
| Avatars K5 | Avatars K10 | 0.792 | 0.033 | 0.175 |
| Avatars K5 | CTGAN | 1.000 | 0.000 | 0.000 |
| Avatars K5 | Gaussian Copula | 0.002 | 0.001 | 0.997 |
| Avatars K5 | Synthpop | 1.000 | 0.000 | 0.000 |
| Avatars K5 | TVAE | 0.994 | 0.002 | 0.004 |
| Avatars K10 | Avatars K5 | 0.033 | 0.792 | 0.175 |
| Avatars K10 | CTGAN | 0.976 | 0.005 | 0.019 |
| Avatars K10 | Gaussian Copula | 0.035 | 0.762 | 0.203 |
| Avatars K10 | Synthpop | 0.779 | 0.025 | 0.196 |
| Avatars K10 | TVAE | 0.985 | 0.005 | 0.011 |
| CTGAN | Avatars K5 | 0.000 | 1.000 | 0.000 |
| CTGAN | Avatars K10 | 0.005 | 0.976 | 0.019 |

| | | | | |
|---|---|---|---|---|
| CTGAN | Gaussian Copula | 0.000 | 1.000 | 0.000 |
| CTGAN | Synthpop | 0.000 | 0.995 | 0.004 |
| CTGAN | TVAE | 0.567 | 0.106 | 0.328 |
| Gaussian Copula | Avatars K5 | 0.001 | 0.002 | 0.997 |
| Gaussian Copula | Avatars K10 | 0.762 | 0.035 | 0.203 |
| Gaussian Copula | CTGAN | 1.000 | 0.000 | 0.000 |
| Gaussian Copula | Synthpop | 1.000 | 0.000 | 0.000 |
| Gaussian Copula | TVAE | 0.995 | 0.002 | 0.003 |
| Synthpop | Avatars K5 | 0.000 | 1.000 | 0.000 |
| Synthpop | Avatars K10 | 0.025 | 0.779 | 0.196 |
| Synthpop | CTGAN | 0.995 | 0.000 | 0.004 |
| Synthpop | Gaussian Copula | 0.000 | 1.000 | 0.000 |
| Synthpop | TVAE | 0.939 | 0.013 | 0.047 |
| TVAE | Avatars K5 | 0.002 | 0.994 | 0.004 |
| TVAE | Avatars K10 | 0.005 | 0.985 | 0.011 |
| TVAE | CTGAN | 0.106 | 0.567 | 0.328 |
| TVAE | Gaussian Copula | 0.002 | 0.995 | 0.003 |
| TVAE | Synthpop | 0.013 | 0.939 | 0.047 |

### 2. Narrow utility

#### 2.1 Differential gene expression (DGE)

**Table S7.** Pairwise Bayesian comparison of SDG methods based on GCS in the ccRCC dataset.

| Method 1 | Method 2 | Better Prob | Worse Prob | Equivalent Prob |
|---|---|---|---|---|
| Avatars K5 | Avatars K10 | 0.664 | 0.101 | 0.235 |
| Avatars K5 | CTGAN | 0.996 | 0.002 | 0.003 |
| Avatars K5 | Gaussian Copula | 0.045 | 0.859 | 0.096 |
| Avatars K5 | Synthpop | 0.996 | 0.002 | 0.002 |
| Avatars K10 | Avatars K5 | 0.101 | 0.664 | 0.235 |
| Avatars K10 | CTGAN | 0.995 | 0.001 | 0.004 |
| Avatars K10 | Gaussian Copula | 0.038 | 0.911 | 0.051 |
| Avatars K10 | Synthpop | 0.968 | 0.013 | 0.019 |
| CTGAN | Avatars K5 | 0.002 | 0.996 | 0.003 |
| CTGAN | Avatars K10 | 0.001 | 0.995 | 0.004 |
| CTGAN | Gaussian Copula | 0.002 | 0.996 | 0.002 |
| CTGAN | Synthpop | 0.631 | 0.094 | 0.275 |
| Gaussian Copula | Avatars K5 | 0.859 | 0.045 | 0.096 |
| Gaussian Copula | Avatars K10 | 0.911 | 0.038 | 0.051 |
| Gaussian Copula | CTGAN | 0.996 | 0.002 | 0.002 |
| Gaussian Copula | Synthpop | 0.999 | 0.000 | 0.000 |
| Synthpop | Avatars K5 | 0.002 | 0.996 | 0.002 |
| Synthpop | Avatars K10 | 0.013 | 0.968 | 0.019 |
| Synthpop | CTGAN | 0.094 | 0.631 | 0.275 |
| Synthpop | Gaussian Copula | 0.000 | 0.999 | 0.000 |

**Table S8.** Pairwise Bayesian comparison of SDG methods based on GCS in the Melanoma dataset.

| Method 1 | Method 2 | Better Prob | Worse Prob | Equivalent Prob |
|---|---|---|---|---|
| Avatars K5 | Avatars K10 | 0.491 | 0.233 | 0.276 |
| Avatars K5 | CTGAN | 0.997 | 0.002 | 0.002 |
| Avatars K5 | Gaussian Copula | 0.155 | 0.736 | 0.108 |
| Avatars K5 | Synthpop | 0.891 | 0.054 | 0.055 |

| | | | | |
|---|---|---|---|---|
| Avatars K5 | TVAE | 0.686 | 0.137 | 0.177 |
| Avatars K10 | Avatars K5 | 0.233 | 0.491 | 0.276 |
| Avatars K10 | CTGAN | 0.975 | 0.014 | 0.011 |
| Avatars K10 | Gaussian Copula | 0.119 | 0.794 | 0.087 |
| Avatars K10 | Synthpop | 0.793 | 0.121 | 0.086 |
| Avatars K10 | TVAE | 0.549 | 0.262 | 0.188 |
| CTGAN | Avatars K5 | 0.002 | 0.997 | 0.002 |
| CTGAN | Avatars K10 | 0.014 | 0.975 | 0.011 |
| CTGAN | Gaussian Copula | 0.002 | 0.997 | 0.001 |
| CTGAN | Synthpop | 0.004 | 0.985 | 0.011 |
| CTGAN | TVAE | 0.010 | 0.980 | 0.010 |
| Gaussian Copula | Avatars K5 | 0.736 | 0.155 | 0.108 |
| Gaussian Copula | Avatars K10 | 0.794 | 0.119 | 0.087 |
| Gaussian Copula | CTGAN | 0.997 | 0.002 | 0.001 |
| Gaussian Copula | Synthpop | 0.996 | 0.002 | 0.002 |
| Gaussian Copula | TVAE | 0.861 | 0.080 | 0.059 |
| Synthpop | Avatars K5 | 0.054 | 0.891 | 0.055 |
| Synthpop | Avatars K10 | 0.121 | 0.793 | 0.086 |
| Synthpop | CTGAN | 0.985 | 0.004 | 0.011 |
| Synthpop | Gaussian Copula | 0.002 | 0.996 | 0.002 |
| Synthpop | TVAE | 0.136 | 0.739 | 0.125 |
| TVAE | Avatars K5 | 0.137 | 0.686 | 0.177 |
| TVAE | Avatars K10 | 0.262 | 0.549 | 0.188 |
| TVAE | CTGAN | 0.980 | 0.010 | 0.010 |
| TVAE | Gaussian Copula | 0.080 | 0.861 | 0.059 |
| TVAE | Synthpop | 0.739 | 0.136 | 0.125 |

**Table S9.** Pairwise Bayesian comparison of SDG methods based on GCS in the NSCLC dataset.

| Method 1 | Method 2 | Better Prob | Worse Prob | Equivalent Prob |
|---|---|---|---|---|
| Avatars K5 | Avatars K10 | 0.777 | 0.075 | 0.148 |
| Avatars K5 | CTGAN | 0.995 | 0.002 | 0.003 |

| | | | | |
|---|---|---|---|---|
| Avatars K5 | Gaussian Copula | 0.005 | 0.989 | 0.006 |
| Avatars K5 | Synthpop | 0.986 | 0.004 | 0.010 |
| Avatars K5 | TVAE | 0.408 | 0.165 | 0.427 |
| Avatars K10 | Avatars K5 | 0.075 | 0.777 | 0.148 |
| Avatars K10 | CTGAN | 0.925 | 0.021 | 0.054 |
| Avatars K10 | Gaussian Copula | 0.002 | 0.997 | 0.001 |
| Avatars K10 | Synthpop | 0.723 | 0.107 | 0.170 |
| Avatars K10 | TVAE | 0.090 | 0.709 | 0.201 |
| CTGAN | Avatars K5 | 0.002 | 0.995 | 0.003 |
| CTGAN | Avatars K10 | 0.021 | 0.925 | 0.054 |
| CTGAN | Gaussian Copula | 0.000 | 1.000 | 0.000 |
| CTGAN | Synthpop | 0.049 | 0.645 | 0.306 |
| CTGAN | TVAE | 0.001 | 0.996 | 0.002 |
| Gaussian Copula | Avatars K5 | 0.989 | 0.005 | 0.006 |
| Gaussian Copula | Avatars K10 | 0.997 | 0.002 | 0.001 |
| Gaussian Copula | CTGAN | 1.000 | 0.000 | 0.000 |
| Gaussian Copula | Synthpop | 0.998 | 0.001 | 0.001 |
| Gaussian Copula | TVAE | 0.986 | 0.007 | 0.007 |
| Synthpop | Avatars K5 | 0.004 | 0.986 | 0.010 |
| Synthpop | Avatars K10 | 0.107 | 0.723 | 0.170 |
| Synthpop | CTGAN | 0.645 | 0.049 | 0.306 |
| Synthpop | Gaussian Copula | 0.001 | 0.998 | 0.001 |
| Synthpop | TVAE | 0.020 | 0.939 | 0.041 |
| TVAE | Avatars K5 | 0.165 | 0.408 | 0.427 |
| TVAE | Avatars K10 | 0.709 | 0.090 | 0.201 |
| TVAE | CTGAN | 0.996 | 0.001 | 0.002 |
| TVAE | Gaussian Copula | 0.007 | 0.986 | 0.007 |
| TVAE | Synthpop | 0.939 | 0.020 | 0.041 |

**Table S10**. Spearman rank correlations between gene-wise $log_2(FC)$ of synthetic datasets and original data across three cancer cohorts.

| Cancer | Methods | Replicate 1 | Replicate 2 | Replicate 3 | Replicate 4 | Replicate 5 |
|---|---|---|---|---|---|---|
| ccRCC | Avatars K5 | 0.19 | 0.05 | 0.09 | 0.14 | 0.11 |
| | Avatars K10 | 0.2 | 0.16 | 0.09 | 0.11 | 0.15 |
| | CTGAN | 0.0 | 0.01 | 0.0 | 0.0 | 0.0 |
| | Gaussian Copula | 0.63 | 0.32 | 0.25 | 0.28 | 0.29 |
| | Synthpop | 0.12 | 0.0 | 0.0 | 0.0 | 0.0 |
| Melanoma | Avatars K5 | 0.37 | 0.25 | 0.34 | 0.02 | 0.2 |
| | Avatars K10 | 0.19 | 0.37 | 0.29 | 0.12 | 0.29 |
| | CTGAN | 0.0 | 0.0 | 0.01 | 0.0 | 0.0 |
| | Gaussian Copula | 0.49 | 0.4 | 0.45 | 0.41 | 0.45 |
| | Synthpop | 0.07 | 0.0 | 0.11 | 0.16 | 0.11 |
| | TVAE | 0.23 | 0.26 | 0.18 | 0.17 | 0.38 |
| NSCLC | Avatars K5 | 0.14 | 0.03 | 0.26 | 0.1 | 0.28 |
| | Avatars K10 | 0.26 | 0.22 | 0.21 | 0.12 | 0.26 |
| | CTGAN | 0.01 | 0.01 | 0.0 | 0.0 | 0.0 |
| | Gaussian Copula | 0.47 | 0.46 | 0.45 | 0.45 | 0.45 |
| | Synthpop | 0.0 | 0.04 | 0.0 | 0.0 | 0.0 |
| | TVAE | 0.18 | 0.16 | 0.24 | 0.13 | 0.01 |

#### 2.2 Gene set enrichment analysis (GSEA)

**Table S11.** Pairwise Bayesian comparison of SDG methods based on PCS in the ccRCC dataset.

| Method 1 | Method 2 | Better Prob | Worse Prob | Equivalent Prob |
|---|---|---|---|---|
| Avatars K5 | Avatars K10 | 0.085 | 0.837 | 0.077 |
| Avatars K5 | CTGAN | 0.490 | 0.167 | 0.343 |
| Avatars K5 | Gaussian Copula | 0.366 | 0.580 | 0.054 |
| Avatars K5 | Synthpop | 0.915 | 0.057 | 0.028 |
| Avatars K10 | Avatars K5 | 0.837 | 0.085 | 0.077 |
| Avatars K10 | CTGAN | 0.868 | 0.072 | 0.060 |
| Avatars K10 | Gaussian Copula | 0.507 | 0.438 | 0.055 |
| Avatars K10 | Synthpop | 0.956 | 0.032 | 0.012 |
| CTGAN | Avatars K5 | 0.167 | 0.490 | 0.343 |
| CTGAN | Avatars K10 | 0.072 | 0.868 | 0.060 |

| | | | | |
|---|---|---|---|---|
| CTGAN | Gaussian Copula | 0.353 | 0.597 | 0.049 |
| CTGAN | Synthpop | 0.879 | 0.084 | 0.037 |
| Gaussian Copula | Avatars K5 | 0.580 | 0.366 | 0.054 |
| Gaussian Copula | Avatars K10 | 0.438 | 0.507 | 0.055 |
| Gaussian Copula | CTGAN | 0.597 | 0.353 | 0.049 |
| Gaussian Copula | Synthpop | 0.854 | 0.118 | 0.028 |
| Synthpop | Avatars K5 | 0.057 | 0.915 | 0.028 |
| Synthpop | Avatars K10 | 0.032 | 0.956 | 0.012 |
| Synthpop | CTGAN | 0.084 | 0.879 | 0.037 |
| Synthpop | Gaussian Copula | 0.118 | 0.854 | 0.028 |

**Table S12.** Pairwise Bayesian comparison of SDG methods based on PCS in the Melanoma dataset.

| Method 1 | Method 2 | Better Prob | Worse Prob | Equivalent Prob |
|---|---|---|---|---|
| Avatars K5 | Avatars K10 | 0.388 | 0.566 | 0.046 |
| Avatars K5 | CTGAN | 0.962 | 0.030 | 0.007 |
| Avatars K5 | Gaussian Copula | 0.777 | 0.187 | 0.036 |
| Avatars K5 | Synthpop | 0.689 | 0.273 | 0.038 |
| Avatars K5 | TVAE | 0.115 | 0.851 | 0.034 |
| Avatars K10 | Avatars K5 | 0.566 | 0.388 | 0.046 |
| Avatars K10 | CTGAN | 0.977 | 0.018 | 0.004 |
| Avatars K10 | Gaussian Copula | 0.782 | 0.191 | 0.027 |
| Avatars K10 | Synthpop | 0.767 | 0.201 | 0.032 |
| Avatars K10 | TVAE | 0.286 | 0.672 | 0.042 |
| CTGAN | Avatars K5 | 0.030 | 0.962 | 0.007 |
| CTGAN | Avatars K10 | 0.018 | 0.977 | 0.004 |
| CTGAN | Gaussian Copula | 0.087 | 0.884 | 0.029 |
| CTGAN | Synthpop | 0.128 | 0.845 | 0.028 |
| CTGAN | TVAE | 0.002 | 0.997 | 0.001 |
| Gaussian Copula | Avatars K5 | 0.187 | 0.777 | 0.036 |
| Gaussian Copula | Avatars K10 | 0.191 | 0.782 | 0.027 |
| Gaussian Copula | CTGAN | 0.884 | 0.087 | 0.029 |
| Gaussian Copula | Synthpop | 0.393 | 0.545 | 0.062 |
| Gaussian Copula | TVAE | 0.021 | 0.973 | 0.006 |
| Synthpop | Avatars K5 | 0.273 | 0.689 | 0.038 |
| Synthpop | Avatars K10 | 0.201 | 0.767 | 0.032 |
| Synthpop | CTGAN | 0.845 | 0.128 | 0.028 |
| Synthpop | Gaussian Copula | 0.545 | 0.393 | 0.062 |
| Synthpop | TVAE | 0.128 | 0.853 | 0.019 |
| TVAE | Avatars K5 | 0.851 | 0.115 | 0.034 |
| TVAE | Avatars K10 | 0.672 | 0.286 | 0.042 |
| TVAE | CTGAN | 0.997 | 0.002 | 0.001 |

| | | | | |
|---|---|---|---|---|
| TVAE | Gaussian Copula | 0.973 | 0.021 | 0.006 |
| TVAE | Synthpop | 0.853 | 0.128 | 0.019 |

**Table S13.** Pairwise Bayesian comparison of SDG methods based on PCS in the NSCLC dataset.

| Method 1 | Method 2 | Better Prob | Worse Prob | Equivalent Prob |
|---|---|---|---|---|
| Avatars K5 | Avatars K10 | 0.579 | 0.366 | 0.055 |
| Avatars K5 | CTGAN | 0.885 | 0.092 | 0.022 |
| Avatars K5 | Gaussian Copula | 0.396 | 0.558 | 0.046 |
| Avatars K5 | Synthpop | 0.886 | 0.093 | 0.021 |
| Avatars K5 | TVAE | 0.676 | 0.270 | 0.053 |
| Avatars K10 | Avatars K5 | 0.366 | 0.579 | 0.055 |
| Avatars K10 | CTGAN | 0.935 | 0.045 | 0.020 |
| Avatars K10 | Gaussian Copula | 0.143 | 0.798 | 0.059 |
| Avatars K10 | Synthpop | 0.998 | 0.001 | 0.001 |
| Avatars K10 | TVAE | 0.649 | 0.223 | 0.128 |
| CTGAN | Avatars K5 | 0.092 | 0.885 | 0.022 |
| CTGAN | Avatars K10 | 0.045 | 0.935 | 0.020 |
| CTGAN | Gaussian Copula | 0.028 | 0.963 | 0.009 |
| CTGAN | Synthpop | 0.514 | 0.325 | 0.161 |
| CTGAN | TVAE | 0.119 | 0.840 | 0.042 |
| Gaussian Copula | Avatars K5 | 0.558 | 0.396 | 0.046 |
| Gaussian Copula | Avatars K10 | 0.798 | 0.143 | 0.059 |
| Gaussian Copula | CTGAN | 0.963 | 0.028 | 0.009 |
| Gaussian Copula | Synthpop | 0.981 | 0.014 | 0.005 |
| Gaussian Copula | TVAE | 0.865 | 0.097 | 0.038 |
| Synthpop | Avatars K5 | 0.093 | 0.886 | 0.021 |
| Synthpop | Avatars K10 | 0.001 | 0.998 | 0.001 |
| Synthpop | CTGAN | 0.325 | 0.514 | 0.161 |
| Synthpop | Gaussian Copula | 0.014 | 0.981 | 0.005 |
| Synthpop | TVAE | 0.024 | 0.961 | 0.015 |
| TVAE | Avatars K5 | 0.270 | 0.676 | 0.053 |
| TVAE | Avatars K10 | 0.223 | 0.649 | 0.128 |
| TVAE | CTGAN | 0.840 | 0.119 | 0.042 |

| | | | | |
|---|---|---|---|---|
| TVAE | Gaussian Copula | 0.097 | 0.865 | 0.038 |
| TVAE | Synthpop | 0.961 | 0.024 | 0.015 |

#### 2.3 Single-sample GSEA

**Table S14.** Pairwise Bayesian comparison of SDG methods based on KS-complement score in ssGSEA in the ccRCC dataset.

| Method 1 | Method 2 | Better Prob | Worse Prob | Equivalent Prob |
|---|---|---|---|---|
| Avatars K5 | Avatars K10 | 0.788 | 0.157 | 0.055 |
| Avatars K5 | CTGAN | 0.900 | 0.067 | 0.033 |
| Avatars K5 | Gaussian Copula | 0.000 | 1.000 | 0.000 |
| Avatars K5 | Synthpop | 0.005 | 0.994 | 0.001 |
| Avatars K5 | TVAE | 0.990 | 0.008 | 0.003 |
| Avatars K10 | Avatars K5 | 0.157 | 0.788 | 0.055 |
| Avatars K10 | CTGAN | 0.572 | 0.318 | 0.110 |
| Avatars K10 | Gaussian Copula | 0.002 | 0.998 | 0.000 |
| Avatars K10 | Synthpop | 0.000 | 1.000 | 0.000 |
| Avatars K10 | TVAE | 0.993 | 0.004 | 0.002 |
| CTGAN | Avatars K5 | 0.067 | 0.900 | 0.033 |
| CTGAN | Avatars K10 | 0.318 | 0.572 | 0.110 |
| CTGAN | Gaussian Copula | 0.000 | 1.000 | 0.000 |
| CTGAN | Synthpop | 0.002 | 0.997 | 0.000 |
| CTGAN | TVAE | 0.980 | 0.013 | 0.008 |
| Gaussian Copula | Avatars K5 | 1.000 | 0.000 | 0.000 |
| Gaussian Copula | Avatars K10 | 0.998 | 0.002 | 0.000 |
| Gaussian Copula | CTGAN | 1.000 | 0.000 | 0.000 |
| Gaussian Copula | Synthpop | 0.971 | 0.023 | 0.006 |
| Gaussian Copula | TVAE | 1.000 | 0.000 | 0.000 |
| Synthpop | Avatars K5 | 0.994 | 0.005 | 0.001 |
| Synthpop | Avatars K10 | 1.000 | 0.000 | 0.000 |
| Synthpop | CTGAN | 0.997 | 0.002 | 0.000 |
| Synthpop | Gaussian Copula | 0.023 | 0.971 | 0.006 |
| Synthpop | TVAE | 1.000 | 0.000 | 0.000 |
| TVAE | Avatars K5 | 0.008 | 0.990 | 0.003 |
| TVAE | Avatars K10 | 0.004 | 0.993 | 0.002 |

| | | | | |
|---|---|---|---|---|
| TVAE | CTGAN | 0.013 | 0.980 | 0.008 |
| TVAE | Gaussian Copula | 0.000 | 1.000 | 0.000 |
| TVAE | Synthpop | 0.000 | 1.000 | 0.000 |

**Table S15.** Pairwise Bayesian comparison of SDG methods based on KS-complement score in ssGSEA in the Melanoma dataset.

| Method 1 | Method 2 | Better Prob | Worse Prob | Equivalent Prob |
|---|---|---|---|---|
| Avatars K5 | Avatars K10 | 0.981 | 0.005 | 0.014 |
| Avatars K5 | CTGAN | 0.074 | 0.820 | 0.106 |
| Avatars K5 | Gaussian Copula | 0.000 | 1.000 | 0.000 |
| Avatars K5 | Synthpop | 0.019 | 0.936 | 0.045 |
| Avatars K5 | TVAE | 0.770 | 0.096 | 0.134 |
| Avatars K10 | Avatars K5 | 0.005 | 0.981 | 0.014 |
| Avatars K10 | CTGAN | 0.007 | 0.985 | 0.008 |
| Avatars K10 | Gaussian Copula | 0.000 | 1.000 | 0.000 |
| Avatars K10 | Synthpop | 0.007 | 0.986 | 0.007 |
| Avatars K10 | TVAE | 0.189 | 0.586 | 0.224 |
| CTGAN | Avatars K5 | 0.820 | 0.074 | 0.106 |
| CTGAN | Avatars K10 | 0.985 | 0.007 | 0.008 |
| CTGAN | Gaussian Copula | 0.001 | 0.998 | 0.001 |
| CTGAN | Synthpop | 0.333 | 0.464 | 0.203 |
| CTGAN | TVAE | 0.857 | 0.088 | 0.056 |
| Gaussian Copula | Avatars K5 | 1.000 | 0.000 | 0.000 |
| Gaussian Copula | Avatars K10 | 1.000 | 0.000 | 0.000 |
| Gaussian Copula | CTGAN | 0.998 | 0.001 | 0.001 |
| Gaussian Copula | Synthpop | 0.998 | 0.001 | 0.001 |

| | | | | |
|---|---|---|---|---|
| Gaussian Copula | TVAE | 0.999 | 0.001 | 0.000 |
| Synthpop | Avatars K5 | 0.936 | 0.019 | 0.045 |
| Synthpop | Avatars K10 | 0.986 | 0.007 | 0.007 |
| Synthpop | CTGAN | 0.464 | 0.333 | 0.203 |
| Synthpop | Gaussian Copula | 0.001 | 0.998 | 0.001 |
| Synthpop | TVAE | 0.953 | 0.023 | 0.024 |
| TVAE | Avatars K5 | 0.096 | 0.770 | 0.134 |
| TVAE | Avatars K10 | 0.586 | 0.189 | 0.224 |
| TVAE | CTGAN | 0.088 | 0.857 | 0.056 |
| TVAE | Gaussian Copula | 0.001 | 0.999 | 0.000 |
| TVAE | Synthpop | 0.023 | 0.953 | 0.024 |

**Table S16.** Pairwise Bayesian comparison of SDG methods based on KS-complement score in ssGSEA in the NSCLC dataset.

| Method 1 | Method 2 | Better Prob | Worse Prob | Equivalent Prob |
|---|---|---|---|---|
| Avatars K5 | Avatars K10 | 0.993 | 0.001 | 0.005 |
| Avatars K5 | CTGAN | 0.742 | 0.206 | 0.052 |
| Avatars K5 | Gaussian Copula | 0.000 | 0.999 | 0.000 |
| Avatars K5 | Synthpop | 0.001 | 0.999 | 0.001 |
| Avatars K5 | TVAE | 0.997 | 0.001 | 0.002 |
| Avatars K10 | Avatars K5 | 0.001 | 0.993 | 0.005 |
| Avatars K10 | CTGAN | 0.607 | 0.321 | 0.072 |
| Avatars K10 | Gaussian Copula | 0.000 | 1.000 | 0.000 |
| Avatars K10 | Synthpop | 0.000 | 1.000 | 0.000 |
| Avatars K10 | TVAE | 0.905 | 0.006 | 0.090 |
| CTGAN | Avatars K5 | 0.206 | 0.742 | 0.052 |
| CTGAN | Avatars K10 | 0.321 | 0.607 | 0.072 |
| CTGAN | Gaussian Copula | 0.027 | 0.966 | 0.007 |
| CTGAN | Synthpop | 0.052 | 0.933 | 0.015 |
| CTGAN | TVAE | 0.397 | 0.525 | 0.078 |
| Gaussian Copula | Avatars K5 | 0.999 | 0.000 | 0.000 |

| | | | | |
|---|---|---|---|---|
| Gaussian Copula | Avatars K10 | 1.000 | 0.000 | 0.000 |
| Gaussian Copula | CTGAN | 0.966 | 0.027 | 0.007 |
| Gaussian Copula | Synthpop | 0.867 | 0.050 | 0.083 |
| Gaussian Copula | TVAE | 1.000 | 0.000 | 0.000 |
| Synthpop | Avatars K5 | 0.999 | 0.001 | 0.001 |
| Synthpop | Avatars K10 | 1.000 | 0.000 | 0.000 |
| Synthpop | CTGAN | 0.933 | 0.052 | 0.015 |
| Synthpop | Gaussian Copula | 0.050 | 0.867 | 0.083 |
| Synthpop | TVAE | 1.000 | 0.000 | 0.000 |
| TVAE | Avatars K5 | 0.001 | 0.997 | 0.002 |
| TVAE | Avatars K10 | 0.006 | 0.905 | 0.090 |
| TVAE | CTGAN | 0.525 | 0.397 | 0.078 |
| TVAE | Gaussian Copula | 0.000 | 1.000 | 0.000 |
| TVAE | Synthpop | 0.000 | 1.000 | 0.000 |

#### 2.4 Cell type deconvolution

**Table S17.** Pairwise Bayesian comparison of SDG methods based on Aitchison distance in cell type deconvolution in the ccRCC dataset.

| Method 1 | Method 2 | Better Prob | Worse Prob | Equivalent Prob |
|---|---|---|---|---|
| Avatars K5 | Avatars K10 | 0.276 | 0.694 | 0.030 |
| Avatars K5 | CTGAN | 0.625 | 0.347 | 0.028 |
| Avatars K5 | Gaussian Copula | 0.192 | 0.781 | 0.026 |
| Avatars K5 | Synthpop | 0.221 | 0.756 | 0.023 |
| Avatars K5 | TVAE | 0.971 | 0.026 | 0.003 |
| Avatars K10 | Avatars K5 | 0.694 | 0.276 | 0.030 |
| Avatars K10 | CTGAN | 0.982 | 0.013 | 0.004 |
| Avatars K10 | Gaussian Copula | 0.226 | 0.682 | 0.092 |
| Avatars K10 | Synthpop | 0.134 | 0.794 | 0.072 |
| Avatars K10 | TVAE | 1.000 | 0.000 | 0.000 |
| CTGAN | Avatars K5 | 0.347 | 0.625 | 0.028 |
| CTGAN | Avatars K10 | 0.013 | 0.982 | 0.004 |

| | | | | |
|---|---|---|---|---|
| CTGAN | Gaussian Copula | 0.011 | 0.986 | 0.003 |
| CTGAN | Synthpop | 0.017 | 0.978 | 0.004 |
| CTGAN | TVAE | 0.998 | 0.002 | 0.000 |
| Gaussian Copula | Avatars K5 | 0.781 | 0.192 | 0.026 |
| Gaussian Copula | Avatars K10 | 0.682 | 0.226 | 0.092 |
| Gaussian Copula | CTGAN | 0.986 | 0.011 | 0.003 |
| Gaussian Copula | Synthpop | 0.383 | 0.532 | 0.085 |
| Gaussian Copula | TVAE | 1.000 | 0.000 | 0.000 |
| Synthpop | Avatars K5 | 0.756 | 0.221 | 0.023 |
| Synthpop | Avatars K10 | 0.794 | 0.134 | 0.072 |
| Synthpop | CTGAN | 0.978 | 0.017 | 0.004 |
| Synthpop | Gaussian Copula | 0.532 | 0.383 | 0.085 |
| Synthpop | TVAE | 1.000 | 0.000 | 0.000 |
| TVAE | Avatars K5 | 0.026 | 0.971 | 0.003 |
| TVAE | Avatars K10 | 0.000 | 1.000 | 0.000 |
| TVAE | CTGAN | 0.002 | 0.998 | 0.000 |
| TVAE | Gaussian Copula | 0.000 | 1.000 | 0.000 |
| TVAE | Synthpop | 0.000 | 1.000 | 0.000 |

**Table S18.** Pairwise Bayesian comparison of SDG methods based on Aitchison distance in cell type deconvolution in the Melanoma dataset.

| Method 1 | Method 2 | Better Prob | Worse Prob | Equivalent Prob |
|---|---|---|---|---|
| Avatars K5 | Avatars K10 | 0.611 | 0.259 | 0.130 |
| Avatars K5 | CTGAN | 0.989 | 0.009 | 0.002 |
| Avatars K5 | Gaussian Copula | 0.972 | 0.014 | 0.014 |
| Avatars K5 | Synthpop | 0.039 | 0.937 | 0.023 |
| Avatars K5 | TVAE | 0.917 | 0.067 | 0.017 |
| Avatars K10 | Avatars K5 | 0.259 | 0.611 | 0.130 |
| Avatars K10 | CTGAN | 0.958 | 0.034 | 0.008 |
| Avatars K10 | Gaussian Copula | 0.899 | 0.051 | 0.050 |
| Avatars K10 | Synthpop | 0.013 | 0.979 | 0.008 |

| | | | | |
|---|---|---|---|---|
| Avatars K10 | TVAE | 0.933 | 0.051 | 0.017 |
| CTGAN | Avatars K5 | 0.009 | 0.989 | 0.002 |
| CTGAN | Avatars K10 | 0.034 | 0.958 | 0.008 |
| CTGAN | Gaussian Copula | 0.031 | 0.959 | 0.010 |
| CTGAN | Synthpop | 0.006 | 0.993 | 0.001 |
| CTGAN | TVAE | 0.289 | 0.671 | 0.040 |
| Gaussian Copula | Avatars K5 | 0.014 | 0.972 | 0.014 |
| Gaussian Copula | Avatars K10 | 0.051 | 0.899 | 0.050 |
| Gaussian Copula | CTGAN | 0.959 | 0.031 | 0.010 |
| Gaussian Copula | Synthpop | 0.001 | 0.999 | 0.000 |
| Gaussian Copula | TVAE | 0.802 | 0.154 | 0.043 |
| Synthpop | Avatars K5 | 0.937 | 0.039 | 0.023 |
| Synthpop | Avatars K10 | 0.979 | 0.013 | 0.008 |
| Synthpop | CTGAN | 0.993 | 0.006 | 0.001 |
| Synthpop | Gaussian Copula | 0.999 | 0.001 | 0.000 |
| Synthpop | TVAE | 0.989 | 0.009 | 0.002 |
| TVAE | Avatars K5 | 0.067 | 0.917 | 0.017 |
| TVAE | Avatars K10 | 0.051 | 0.933 | 0.017 |
| TVAE | CTGAN | 0.671 | 0.289 | 0.040 |
| TVAE | Gaussian Copula | 0.154 | 0.802 | 0.043 |
| TVAE | Synthpop | 0.009 | 0.989 | 0.002 |

**Table S19.** Pairwise Bayesian comparison of SDG methods based on Aitchison distance in cell type deconvolution in the NSCLC dataset.

| Method 1 | Method 2 | Better Prob | Worse Prob | Equivalent Prob |
|---|---|---|---|---|
| Avatars K5 | Avatars K10 | 0.981 | 0.008 | 0.012 |
| Avatars K5 | CTGAN | 1.000 | 0.000 | 0.000 |
| Avatars K5 | Gaussian Copula | 0.923 | 0.041 | 0.036 |
| Avatars K5 | Synthpop | 0.006 | 0.986 | 0.008 |
| Avatars K5 | TVAE | 0.954 | 0.034 | 0.011 |
| Avatars K10 | Avatars K5 | 0.008 | 0.981 | 0.012 |

| | | | | |
|---|---|---|---|---|
| Avatars K10 | CTGAN | 1.000 | 0.000 | 0.000 |
| Avatars K10 | Gaussian Copula | 0.440 | 0.376 | 0.184 |
| Avatars K10 | Synthpop | 0.002 | 0.996 | 0.001 |
| Avatars K10 | TVAE | 0.905 | 0.066 | 0.030 |
| CTGAN | Avatars K5 | 0.000 | 1.000 | 0.000 |
| CTGAN | Avatars K10 | 0.000 | 1.000 | 0.000 |
| CTGAN | Gaussian Copula | 0.003 | 0.997 | 0.001 |
| CTGAN | Synthpop | 0.000 | 1.000 | 0.000 |
| CTGAN | TVAE | 0.034 | 0.955 | 0.011 |
| Gaussian Copula | Avatars K5 | 0.041 | 0.923 | 0.036 |
| Gaussian Copula | Avatars K10 | 0.376 | 0.440 | 0.184 |
| Gaussian Copula | CTGAN | 0.997 | 0.003 | 0.001 |
| Gaussian Copula | Synthpop | 0.005 | 0.993 | 0.003 |
| Gaussian Copula | TVAE | 0.817 | 0.142 | 0.040 |
| Synthpop | Avatars K5 | 0.986 | 0.006 | 0.008 |
| Synthpop | Avatars K10 | 0.996 | 0.002 | 0.001 |
| Synthpop | CTGAN | 1.000 | 0.000 | 0.000 |
| Synthpop | Gaussian Copula | 0.993 | 0.005 | 0.003 |
| Synthpop | TVAE | 0.976 | 0.019 | 0.005 |
| TVAE | Avatars K5 | 0.034 | 0.954 | 0.011 |
| TVAE | Avatars K10 | 0.066 | 0.905 | 0.030 |
| TVAE | CTGAN | 0.955 | 0.034 | 0.011 |
| TVAE | Gaussian Copula | 0.142 | 0.817 | 0.040 |
| TVAE | Synthpop | 0.019 | 0.976 | 0.005 |

#### 2.5 Survival analysis

**Table S20.** Pairwise Bayesian comparison of SDG methods based on C-index score in survival analysis in the ccRCC dataset.

| Method 1 | Method 2 | Better Prob | Worse Prob | Equivalent Prob |
|---|---|---|---|---|
| Avatars K5 | Avatars K10 | 0.050 | 0.022 | 0.928 |

| | | | | |
|---|---|---|---|---|
| Avatars K5 | CTGAN | 1.000 | 0.000 | 0.000 |
| Avatars K5 | Gaussian Copula | 0.877 | 0.026 | 0.097 |
| Avatars K5 | Synthpop | 1.000 | 0.000 | 0.000 |
| Avatars K5 | TVAE | 1.000 | 0.000 | 0.000 |
| Avatars K10 | Avatars K5 | 0.022 | 0.050 | 0.928 |
| Avatars K10 | CTGAN | 1.000 | 0.000 | 0.000 |
| Avatars K10 | Gaussian Copula | 0.891 | 0.019 | 0.091 |
| Avatars K10 | Synthpop | 1.000 | 0.000 | 0.000 |
| Avatars K10 | TVAE | 1.000 | 0.000 | 0.000 |
| CTGAN | Avatars K5 | 0.000 | 1.000 | 0.000 |
| CTGAN | Avatars K10 | 0.000 | 1.000 | 0.000 |
| CTGAN | Gaussian Copula | 0.000 | 0.999 | 0.000 |
| CTGAN | Synthpop | 0.161 | 0.405 | 0.434 |
| CTGAN | TVAE | 1.000 | 0.000 | 0.000 |
| Gaussian Copula | Avatars K5 | 0.026 | 0.877 | 0.097 |
| Gaussian Copula | Avatars K10 | 0.019 | 0.891 | 0.091 |
| Gaussian Copula | CTGAN | 0.999 | 0.000 | 0.000 |
| Gaussian Copula | Synthpop | 1.000 | 0.000 | 0.000 |
| Gaussian Copula | TVAE | 1.000 | 0.000 | 0.000 |
| Synthpop | Avatars K5 | 0.000 | 1.000 | 0.000 |
| Synthpop | Avatars K10 | 0.000 | 1.000 | 0.000 |
| Synthpop | CTGAN | 0.405 | 0.161 | 0.434 |
| Synthpop | Gaussian Copula | 0.000 | 1.000 | 0.000 |
| Synthpop | TVAE | 1.000 | 0.000 | 0.000 |
| TVAE | Avatars K5 | 0.000 | 1.000 | 0.000 |
| TVAE | Avatars K10 | 0.000 | 1.000 | 0.000 |
| TVAE | CTGAN | 0.000 | 1.000 | 0.000 |
| TVAE | Gaussian Copula | 0.000 | 1.000 | 0.000 |
| TVAE | Synthpop | 0.000 | 1.000 | 0.000 |

**Table S21.** Pairwise Bayesian comparison of SDG methods based on C-index score in survival analysis in the Melanoma dataset.

| Method 1 | Method 2 | Better Prob | Worse Prob | Equivalent Prob |
|---|---|---|---|---|
| Avatars K5 | Avatars K10 | 0.273 | 0.427 | 0.300 |
| Avatars K5 | CTGAN | 0.999 | 0.001 | 0.000 |
| Avatars K5 | Gaussian Copula | 0.174 | 0.636 | 0.190 |
| Avatars K5 | Synthpop | 1.000 | 0.000 | 0.000 |
| Avatars K5 | TVAE | 0.845 | 0.099 | 0.056 |
| Avatars K10 | Avatars K5 | 0.427 | 0.273 | 0.300 |
| Avatars K10 | CTGAN | 1.000 | 0.000 | 0.000 |
| Avatars K10 | Gaussian Copula | 0.191 | 0.578 | 0.232 |
| Avatars K10 | Synthpop | 0.998 | 0.001 | 0.001 |
| Avatars K10 | TVAE | 0.861 | 0.089 | 0.050 |
| CTGAN | Avatars K5 | 0.001 | 0.999 | 0.000 |
| CTGAN | Avatars K10 | 0.000 | 1.000 | 0.000 |
| CTGAN | Gaussian Copula | 0.000 | 0.999 | 0.000 |
| CTGAN | Synthpop | 0.163 | 0.426 | 0.411 |
| CTGAN | TVAE | 0.091 | 0.858 | 0.051 |
| Gaussian Copula | Avatars K5 | 0.636 | 0.174 | 0.190 |
| Gaussian Copula | Avatars K10 | 0.578 | 0.191 | 0.232 |
| Gaussian Copula | CTGAN | 0.999 | 0.000 | 0.000 |
| Gaussian Copula | Synthpop | 0.999 | 0.000 | 0.000 |
| Gaussian Copula | TVAE | 0.891 | 0.071 | 0.037 |
| Synthpop | Avatars K5 | 0.000 | 1.000 | 0.000 |
| Synthpop | Avatars K10 | 0.001 | 0.998 | 0.001 |
| Synthpop | CTGAN | 0.426 | 0.163 | 0.411 |
| Synthpop | Gaussian Copula | 0.000 | 0.999 | 0.000 |
| Synthpop | TVAE | 0.117 | 0.822 | 0.061 |
| TVAE | Avatars K5 | 0.099 | 0.845 | 0.056 |
| TVAE | Avatars K10 | 0.089 | 0.861 | 0.050 |
| TVAE | CTGAN | 0.858 | 0.091 | 0.051 |

| | | | | |
|---|---|---|---|---|
| TVAE | Gaussian Copula | 0.071 | 0.891 | 0.037 |
| TVAE | Synthpop | 0.822 | 0.117 | 0.061 |

**Table S22.** Pairwise Bayesian comparison of SDG methods based on C-index score in survival analysis in the NSCLC dataset.

| Method 1 | Method 2 | Better Prob | Worse Prob | Equivalent Prob |
|---|---|---|---|---|
| Avatars K5 | Avatars K10 | 0.530 | 0.016 | 0.455 |
| Avatars K5 | CTGAN | 1.000 | 0.000 | 0.000 |
| Avatars K5 | Gaussian Copula | 0.878 | 0.041 | 0.080 |
| Avatars K5 | Synthpop | 0.998 | 0.001 | 0.001 |
| Avatars K5 | TVAE | 0.999 | 0.001 | 0.000 |
| Avatars K10 | Avatars K5 | 0.016 | 0.530 | 0.455 |
| Avatars K10 | CTGAN | 1.000 | 0.000 | 0.000 |
| Avatars K10 | Gaussian Copula | 0.774 | 0.080 | 0.145 |
| Avatars K10 | Synthpop | 0.998 | 0.001 | 0.001 |
| Avatars K10 | TVAE | 0.999 | 0.001 | 0.001 |
| CTGAN | Avatars K5 | 0.000 | 1.000 | 0.000 |
| CTGAN | Avatars K10 | 0.000 | 1.000 | 0.000 |
| CTGAN | Gaussian Copula | 0.001 | 0.998 | 0.001 |
| CTGAN | Synthpop | 0.037 | 0.841 | 0.123 |
| CTGAN | TVAE | 0.286 | 0.354 | 0.360 |
| Gaussian Copula | Avatars K5 | 0.041 | 0.878 | 0.080 |
| Gaussian Copula | Avatars K10 | 0.080 | 0.774 | 0.145 |
| Gaussian Copula | CTGAN | 0.998 | 0.001 | 0.001 |
| Gaussian Copula | Synthpop | 0.998 | 0.001 | 0.001 |
| Gaussian Copula | TVAE | 0.990 | 0.006 | 0.004 |
| Synthpop | Avatars K5 | 0.001 | 0.998 | 0.001 |
| Synthpop | Avatars K10 | 0.001 | 0.998 | 0.001 |
| Synthpop | CTGAN | 0.841 | 0.037 | 0.123 |
| Synthpop | Gaussian Copula | 0.001 | 0.998 | 0.001 |
| Synthpop | TVAE | 0.686 | 0.150 | 0.164 |

| | | | | |
|---|---|---|---|---|
| TVAE | Avatars K5 | 0.001 | 0.999 | 0.000 |
| TVAE | Avatars K10 | 0.001 | 0.999 | 0.001 |
| TVAE | CTGAN | 0.354 | 0.286 | 0.360 |
| TVAE | Gaussian Copula | 0.006 | 0.990 | 0.004 |
| TVAE | Synthpop | 0.150 | 0.686 | 0.164 |

#### 3. Privacy

**Table S23.** Pairwise Bayesian comparison of SDG methods based on Overall privacy score in privacy risk attacks in the ccRCC dataset.

| Method 1 | Method 2 | Better Prob | Worse Prob | Equivalent Prob |
|---|---|---|---|---|
| Avatars K5 | Avatars K10 | 0.003 | 0.960 | 0.037 |
| Avatars K5 | CTGAN | 0.000 | 0.000 | 1.000 |
| Avatars K5 | Gaussian Copula | 0.000 | 0.000 | 1.000 |
| Avatars K5 | Synthpop | 1.000 | 0.000 | 0.000 |
| Avatars K5 | TVAE | 0.000 | 0.001 | 0.999 |
| Avatars K10 | Avatars K5 | 0.037 | 0.960 | 0.003 |
| Avatars K10 | CTGAN | 0.000 | 0.000 | 1.000 |
| Avatars K10 | Gaussian Copula | 0.000 | 0.000 | 1.000 |
| Avatars K10 | Synthpop | 1.000 | 0.000 | 0.000 |
| Avatars K10 | TVAE | 0.000 | 0.001 | 0.998 |
| CTGAN | Avatars K5 | 1.000 | 0.000 | 0.000 |
| CTGAN | Avatars K10 | 1.000 | 0.000 | 0.000 |
| CTGAN | Gaussian Copula | 0.325 | 0.665 | 0.009 |
| CTGAN | Synthpop | 1.000 | 0.000 | 0.000 |
| CTGAN | TVAE | 0.887 | 0.102 | 0.011 |
| Gaussian Copula | Avatars K5 | 1.000 | 0.000 | 0.000 |
| Gaussian Copula | Avatars K10 | 1.000 | 0.000 | 0.000 |
| Gaussian Copula | CTGAN | 0.009 | 0.665 | 0.325 |
| Gaussian Copula | Synthpop | 1.000 | 0.000 | 0.000 |
| Gaussian Copula | TVAE | 0.756 | 0.234 | 0.010 |
| Synthpop | Avatars K5 | 0.000 | 0.000 | 1.000 |

| | | | | |
|---|---|---|---|---|
| Synthpop | Avatars K10 | 0.000 | 0.000 | 1.000 |
| Synthpop | CTGAN | 0.000 | 0.000 | 1.000 |
| Synthpop | Gaussian Copula | 0.000 | 0.000 | 1.000 |
| Synthpop | TVAE | 0.000 | 0.000 | 1.000 |
| TVAE | Avatars K5 | 0.999 | 0.001 | 0.000 |
| TVAE | Avatars K10 | 0.998 | 0.001 | 0.000 |
| TVAE | CTGAN | 0.011 | 0.102 | 0.887 |
| TVAE | Gaussian Copula | 0.010 | 0.234 | 0.756 |
| TVAE | Synthpop | 1.000 | 0.000 | 0.000 |

**Table S24.** Pairwise Bayesian comparison of SDG methods based on Overall privacy score in privacy risk attacks in the Melanoma dataset.

| Method 1 | Method 2 | Better Prob | Worse Prob | Equivalent Prob |
|---|---|---|---|---|
| Avatars K5 | Avatars K10 | 0.010 | 0.865 | 0.125 |
| Avatars K5 | CTGAN | 0.023 | 0.948 | 0.029 |
| Avatars K5 | Gaussian Copula | 0.000 | 0.171 | 0.829 |
| Avatars K5 | Synthpop | 1.000 | 0.000 | 0.000 |
| Avatars K5 | TVAE | 0.439 | 0.545 | 0.015 |
| Avatars K10 | Avatars K5 | 0.125 | 0.865 | 0.010 |
| Avatars K10 | CTGAN | 0.184 | 0.782 | 0.034 |
| Avatars K10 | Gaussian Copula | 0.007 | 0.823 | 0.171 |
| Avatars K10 | Synthpop | 1.000 | 0.000 | 0.000 |
| Avatars K10 | TVAE | 0.657 | 0.318 | 0.025 |
| CTGAN | Avatars K5 | 0.029 | 0.948 | 0.023 |
| CTGAN | Avatars K10 | 0.034 | 0.782 | 0.184 |
| CTGAN | Gaussian Copula | 0.003 | 0.485 | 0.512 |
| CTGAN | Synthpop | 1.000 | 0.000 | 0.000 |
| CTGAN | TVAE | 0.479 | 0.480 | 0.041 |
| Gaussian Copula | Avatars K5 | 0.829 | 0.171 | 0.000 |
| Gaussian Copula | Avatars K10 | 0.171 | 0.823 | 0.007 |
| Gaussian Copula | CTGAN | 0.512 | 0.485 | 0.003 |

| | | | | |
|---|---|---|---|---|
| Gaussian Copula | Synthpop | 1.000 | 0.000 | 0.000 |
| Gaussian Copula | TVAE | 0.918 | 0.079 | 0.003 |
| Synthpop | Avatars K5 | 0.000 | 0.000 | 1.000 |
| Synthpop | Avatars K10 | 0.000 | 0.000 | 1.000 |
| Synthpop | CTGAN | 0.000 | 0.000 | 1.000 |
| Synthpop | Gaussian Copula | 0.000 | 0.000 | 1.000 |
| Synthpop | TVAE | 0.000 | 0.000 | 1.000 |
| TVAE | Avatars K5 | 0.015 | 0.545 | 0.439 |
| TVAE | Avatars K10 | 0.025 | 0.318 | 0.657 |
| TVAE | CTGAN | 0.041 | 0.480 | 0.479 |
| TVAE | Gaussian Copula | 0.003 | 0.079 | 0.918 |
| TVAE | Synthpop | 1.000 | 0.000 | 0.000 |

**Table S25.** Pairwise Bayesian comparison of SDG methods based on Overall privacy score in privacy risk attacks in the NSCLC dataset.

| Method 1 | Method 2 | Better Prob | Worse Prob | Equivalent Prob |
|---|---|---|---|---|
| Avatars K5 | Avatars K10 | 0.004 | 0.802 | 0.194 |
| Avatars K5 | CTGAN | 0.000 | 0.000 | 1.000 |
| Avatars K5 | Gaussian Copula | 0.000 | 0.000 | 1.000 |
| Avatars K5 | Synthpop | 1.000 | 0.000 | 0.000 |
| Avatars K5 | TVAE | 0.056 | 0.788 | 0.156 |
| Avatars K10 | Avatars K5 | 0.194 | 0.802 | 0.004 |
| Avatars K10 | CTGAN | 0.001 | 0.024 | 0.975 |
| Avatars K10 | Gaussian Copula | 0.000 | 0.011 | 0.989 |
| Avatars K10 | Synthpop | 1.000 | 0.000 | 0.000 |
| Avatars K10 | TVAE | 0.210 | 0.737 | 0.053 |
| CTGAN | Avatars K5 | 1.000 | 0.000 | 0.000 |
| CTGAN | Avatars K10 | 0.975 | 0.024 | 0.001 |
| CTGAN | Gaussian Copula | 0.219 | 0.776 | 0.005 |
| CTGAN | Synthpop | 1.000 | 0.000 | 0.000 |
| CTGAN | TVAE | 0.981 | 0.018 | 0.002 |
| Gaussian Copula | Avatars K5 | 1.000 | 0.000 | 0.000 |

| | | | | |
|---|---|---|---|---|
| Gaussian Copula | Avatars K10 | 0.989 | 0.011 | 0.000 |
| Gaussian Copula | CTGAN | 0.005 | 0.776 | 0.219 |
| Gaussian Copula | Synthpop | 1.000 | 0.000 | 0.000 |
| Gaussian Copula | TVAE | 0.936 | 0.061 | 0.003 |
| Synthpop | Avatars K5 | 0.000 | 0.000 | 1.000 |
| Synthpop | Avatars K10 | 0.000 | 0.000 | 1.000 |
| Synthpop | CTGAN | 0.000 | 0.000 | 1.000 |
| Synthpop | Gaussian Copula | 0.000 | 0.000 | 1.000 |
| Synthpop | TVAE | 0.000 | 0.000 | 1.000 |
| TVAE | Avatars K5 | 0.156 | 0.788 | 0.056 |
| TVAE | Avatars K10 | 0.053 | 0.737 | 0.210 |
| TVAE | CTGAN | 0.002 | 0.018 | 0.981 |
| TVAE | Gaussian Copula | 0.003 | 0.061 | 0.936 |
| TVAE | Synthpop | 1.000 | 0.000 | 0.000 |

412
